# Designing enzymes for new-to-nature chemistry and non-natural substrates with AlphaProtein Novo

**DOI:** 10.64898/2026.10.01.756017

**Authors:** Zachary Wu, Josh Abramson, Thomas Frerix, Alexander E. Chu, Ruijie K. Zhang, Luca Schulz, Amy E. Danson, Tristan O. C. Kwan, Li Kevin Wenliang, Jacob Kelly, Zi-Qi Li, Rosalia G. Schneider, Ashok Thillaisundaram, Harshnira Patani, Vinicius F. Zambaldi, Sukhdeep Singh, David La, Masy Domecillo, Ariane N. Mora, Julia C. Reisenbauer, Yu Zhang, Eliseo Papa, Akvilė Žemgulytė, Yu-Han Wu, Augustin Žídek, Jiaxin Shi, Grace Margand, Naila Assem, Kate Stephen, Charlie Emrich, Peng Liu, Lucy Colwell, Demis Hassabis, Rob Fergus, Frances H. Arnold, Pushmeet Kohli, Jue Wang

## Abstract

Creating highly active enzymes for arbitrary reactions is a transformative goal for the molecular sciences^1^. Traditional protein engineering^2,3^ is constrained by a reliance on existing natural starting points, which are often challenging to discover. *De novo* design offers the potential to overcome this by creating enzymes from first principles^4^, but has yet to achieve practically relevant catalytic properties. Here we present AlphaProtein Novo (AP Novo), a machine-learning pipeline that demonstrates, for the first time, that *de novo* enzyme design can outperform natural sequence mining in addressing challenging chemistry. We used AP Novo to design new-to-nature nitrene transferases to synthesise the pharmacophore piperidine with unprecedented product selectivity, and to create enzymes that degrade the environmental toxin DEHP under conditions that denature natural enzymes. We also obtained state-of-the-art catalytic efficiencies on two well-studied model reactions, and through iterative design and analysis found that key enablers of success were mechanism-inspired metrics based on AlphaFold 3^5^ predictions and sequence-ensemble-based scoring of designed backbones. Our results show that *de novo* design has become a powerful complement to natural enzyme diversity for discovering catalysts for a range of applications.

## Main

The design of enzymes for arbitrary reactions is a longstanding goal in biochemistry, with the potential to address major challenges in medicine, industry, and sustainability^1^. While directed evolution is a proven method to optimise enzymes^2,3^, identifying natural sequences with target activity to serve as starting points for engineering is laborious and failure-prone^6^. *De novo* design could in principle provide active starting points with tailored properties for any reaction^4^, but is not yet a standard tool for biocatalyst discovery.

Supported by recent advances in structure generation and prediction^5,7–9^ *de novo* approaches have matured for protein binder design. Current methods are able to design high-affinity binders against a majority of protein targets^10–13^ and this success is beginning to extend to binding small molecules^8,13–15^. *De novo* approaches for enzyme design have also advanced, achieving significant improvements to catalytic efficiency^16–18^ compared to previous approaches and beginning to access new-to-nature chemical space^19–23^.

However, key limitations have so far hindered the adoption of *de novo* enzyme design. Firstly, much recent progress has focused on reactions for which natural enzymes are already highly optimised^17,18,24,25^, and the catalytic rates of those natural enzymes remain orders of magnitude larger than the best *de novo* designs. Secondly, while *de novo* design with natural sidechains or cofactors has improved for useful new-to-nature chemistry such as carbene transfer^19,21,23^ and Morita-Baylis-Hillman^20,22^, the simple model reaction kemp elimination remains the only reaction where *de novo* enzymes outperform natural sequences or derived mutants^26^. Lastly, analysis of *de novo* enzyme designs has so far been performed on a relatively small number of designs and on even fewer distinct protein scaffolds^7,17,27,28^, limiting the generality of any resulting methodological insights.

In this work we present AlphaProtein Novo (AP Novo; APN), a machine-learning pipeline for *de novo* enzyme design, and show that it outperforms natural sequence or mutant screening for two challenging, practically relevant selectivity problems (Fig. 1). We used APN to design enzymes for synthesising piperidines, a key motif in pharmaceuticals and materials^29^, and for hydrolysing di(2-ethylhexyl) phthalate (DEHP), a pervasive environmental toxin^30^. For the first time to our knowledge, we directly compare the efficacy of *de novo* design to screening natural sequences or mutants, the standard approach to practical biocatalyst discovery^31^. We find that the frequency of functional designs from APN matches or surpasses that of natural sequence screening, and our lead piperidine synthase inverts the pyrrolidine-favouring bias of natural enzymes to achieve near-perfect piperidine regioselectivity.

**Figure 1.**
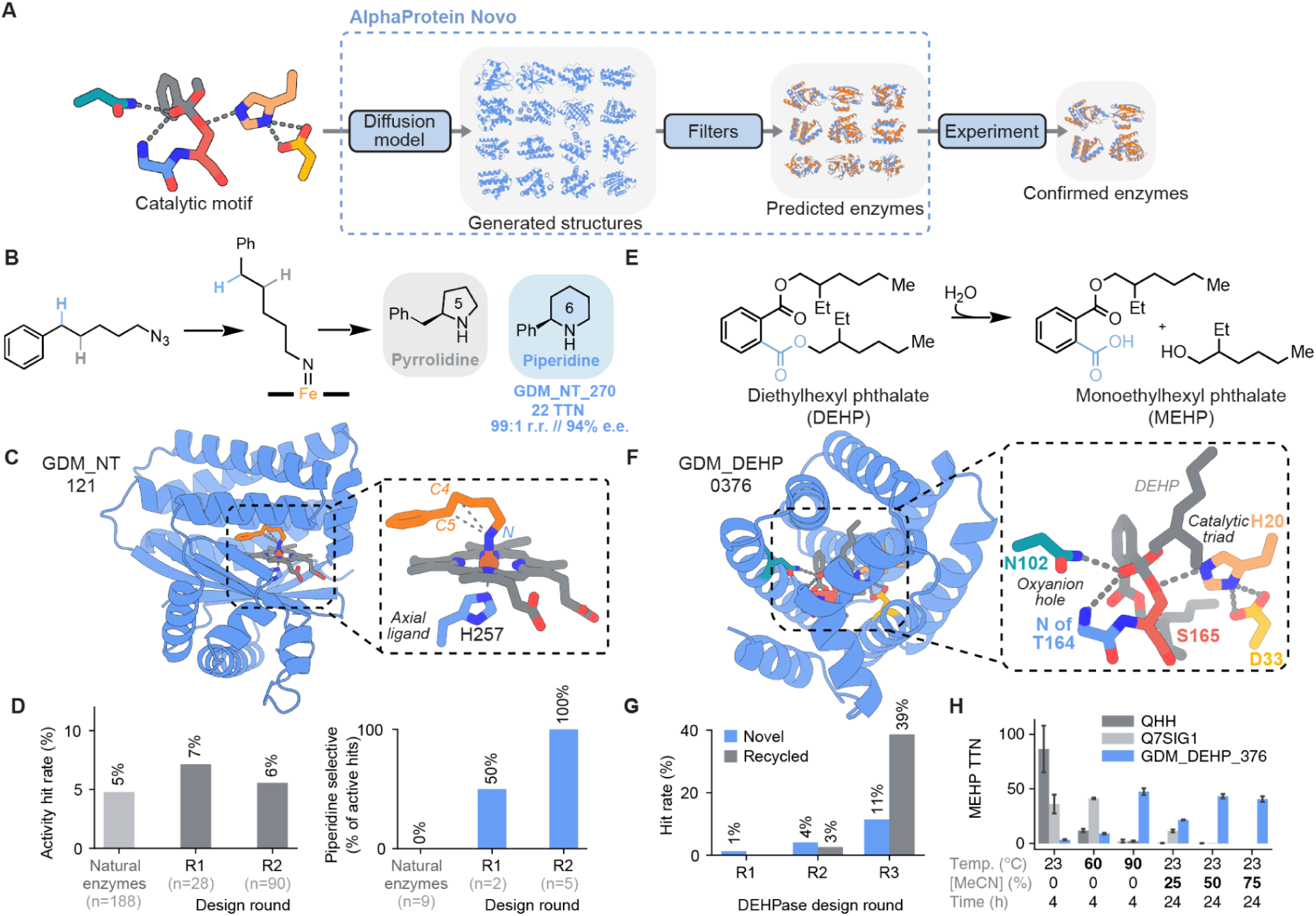
Using AlphaProtein Novo to design enzymes for new-to-nature reactions. A) Schematic of the design pipeline. B) Reaction scheme for the formation of pyrrolidines and piperidines using nitrene transferases. C) Structure view of GDM_NT_121, a regioselective novel-scaffold nitrene transferase. D) Relative hit rates for AP Novo designs for piperidine versus natural diversity screening for the reaction shown in (B). An activity hit (gray) is defined as having at least 5% total C-H amination activity relative to *Tam*Pgb wt. Of these, piperidine-selective hits (blue) are defined by a regioselective preference for piperidine. E) Reaction scheme for the transformation of DEHP to MEHP, the first step in DEHP degradation. F) Structure view of GDM_DEHP_376, a highly-active designed enzyme for DEHP degradation. G) Relative hit rates for designing DEHPases over successive rounds of novel (blue) and recycled (grey) scaffold design (hits classified as having at least 2% activity of control enzyme QHH). Hit rates were calculated using validation data for Rounds 1 and 2, and primary screen data for Round 3. H) Formation of MEHP from DEHP (total turnover number, TTN) at different temperatures, acetonitrile (MeCN) concentrations, and reaction times, for DEHPase design GDM_DEHP_376 and natural esterases QHH and Q7SIG1. Plots show mean values with error bars representing the standard deviation of 2-4 biological replicates.

The APN pipeline designs enzymes by motif scaffolding using a diffusion model, followed by filtering of candidate designs (see Methods). The filtering uses AF3 predictions to assess mechanistically relevant atomic details for each reaction, an approach we verified and improved by analysing data from multiple design-test-learn cycles on three well-studied model reactions. In the course of these design campaigns we obtained state-of-the-art catalytic efficiencies without iteratively optimising known active designs, including a Kemp eliminase with *k*_cat_/K_m_ 45,000 M^-1^s^-1^ at pH 10, another with *k*_cat_/K_m_ 14,600 M^-1^s^-1^ at pH 7.5, a serine esterase with *k*_cat_/K_m_ 360,000 M^-1^s^-1^ that has structural similarity to natural enzymes, and one with 4,900 M^-1^s^-1^ with no natural similarity (Table 1, Table S1). Overall, we found that a key predictor of design success was a novel filtering strategy which evaluates ensembles of sequences derived from the same backbone.

**Table 1.**
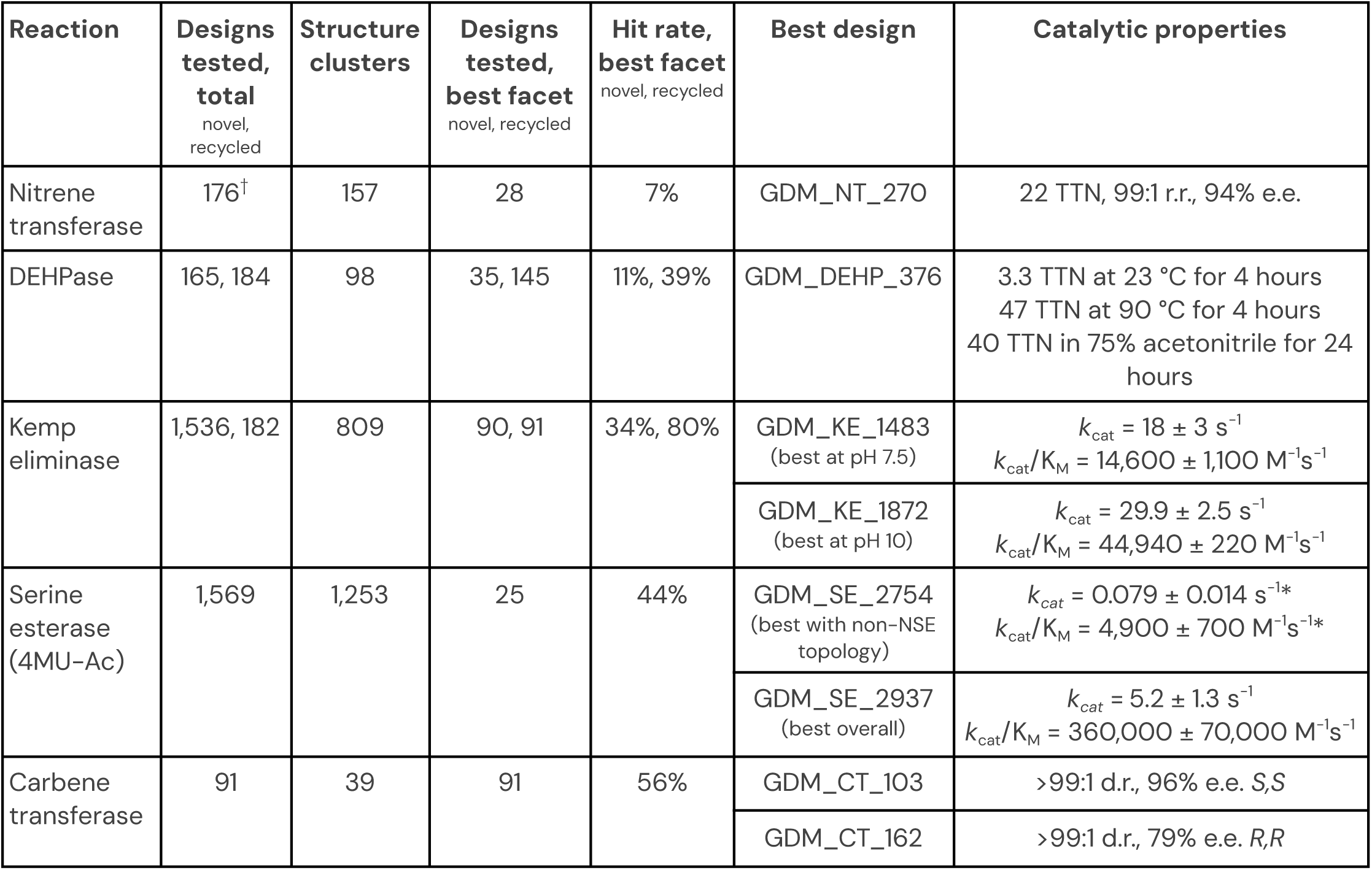
Design outcomes by reaction. All designs are novel-scaffold except in the case of DEHPases and Kemp eliminases, where design numbers and hit rates are shown separately for novel- and recycled-scaffold designs. Each round of design is organised into “facets”, or subsets of designs created using the same pipeline settings. We report the highest per-facet hit rates (and the associated number of designs tested) as a measure of the performance of the most optimised design settings. “Best design” may not come from the “best facet”. Hit rate definitions are reaction-specific and not intended to be compared across reactions. The number of structural clusters is determined by single-linkage greedy clustering with TM-align. Errors on catalytic parameters represent standard deviation. For Kemp eliminases, the best designs at both pH 7.5 and pH 10 are shown (Fig. S10). For serine esterases, the best designs with non-NSE and any topology are shown (see main text, Fig. S11-13). See also Tables S1-2 and Fig. S14 for Michaelis-Menten parameters and hit rates in comparison to previous studies. ^†^The total number of nitrene transferases includes 58 designs from round 1 that were not targeted for piperidine synthesis. *GDM_SE_2754 parameters are fit with substrate inhibition.

| Reaction | Designs tested, total<br>novel, recycled | Structure clusters | Designs tested, best facet<br>novel, recycled | Hit rate, best facet<br>novel, recycled | Best design | Catalytic properties |
| --- | --- | --- | --- | --- | --- | --- |
| Nitrene transferase | 176<sup>†</sup> | 157 | 28 | 7% | GDM_NT_270 | 22 TTN, 99:1 r.r., 94% e.e. |
| DEHPase | 165, 184 | 98 | 35, 145 | 11%, 39% | GDM_DEHP_376 | 3.3 TTN at 23 °C for 4 hours<br>47 TTN at 90 °C for 4 hours<br>40 TTN in 75% acetonitrile for 24 hours |
| Kemp eliminase | 1,536, 182 | 809 | 90, 91 | 34%, 80% | GDM_KE_1483<br>(best at pH 7.5) | $k_{cat} = 18 \pm 3 \text{ s}^{-1}$<br>$k_{cat}/K_M = 14,600 \pm 1,100 \text{ M}^{-1}\text{s}^{-1}$ |
| | | | | | GDM_KE_1872<br>(best at pH 10) | $k_{cat} = 29.9 \pm 2.5 \text{ s}^{-1}$<br>$k_{cat}/K_M = 44,940 \pm 220 \text{ M}^{-1}\text{s}^{-1}$ |
| Serine esterase (4MU-Ac) | 1,569 | 1,253 | 25 | 44% | GDM_SE_2754<br>(best with non-NSE topology) | $k_{cat} = 0.079 \pm 0.014 \text{ s}^{-1*}$<br>$k_{cat}/K_M = 4,900 \pm 700 \text{ M}^{-1}\text{s}^{-1*}$ |
| | | | | | GDM_SE_2937<br>(best overall) | $k_{cat} = 5.2 \pm 1.3 \text{ s}^{-1}$<br>$k_{cat}/K_M = 360,000 \pm 70,000 \text{ M}^{-1}\text{s}^{-1}$ |
| Carbene transferase | 91 | 39 | 91 | 56% | GDM_CT_103 | >99:1 d.r., 96% e.e. S,S |
|  |  |  |  |  | GDM_CT_162 | >99:1 d.r., 79% e.e. R,R |

## Results and discussion

In the results that follow, a design is “novel-scaffold” if the design process only has an input structural catalytic motif, without using a full existing active enzyme. This contrasts with “recycled-scaffold” designs, which come from optimisation of scaffolds with known activity.

### Design of regioselective nitrene transferases for piperidine synthesis

One expected advantage of *de novo* design over natural sequence mining is direct control over scaffold geometry. To test this, we used APN to design enzymes for C-H amination, a reaction which enables efficient access to nitrogen-containing scaffolds^32^, including heterocycles such as pyrrolidines and piperidines. These heterocycles are found in functional materials and bioactive molecules, and piperidines in particular are frequent components in FDA-approved drugs^29^. Previously, it was demonstrated that pyrrolidines can be accessed through new-to-nature intramolecular nitrene transfer by an engineered P450^33^. We aimed to broaden the biocatalytic product scope to include piperidines.

The competitive cyclisation of (5-azidopentyl)benzene can yield either pyrrolidine or piperidine products (Fig. 1B, S3A), which can provide an efficient route to piperidine if regioselectivity is achieved. Pyrrolidine (PYR) formation is achieved via a typically favorable 1,5-hydrogen atom transfer (HAT) geometry^34^, while piperidine (PIP) formation is driven by the lower bond dissociation energy (BDE) of the benzylic C-H bond^35^. Initial attempts at biocatalytic access to piperidine from this substrate focused on screening natural and previously engineered heme proteins. In a screen of 188 variants, nine enzymes exhibited activity above a functional threshold (defined as having at least 5% total C-H amination activity relative to wild-type protoglobin from *Thermus amyloliquefaciens*, *Tam*Pgb wt) (Fig. 1D, S3B). Of those exceeding the threshold, none surpassed a 30:70 (PIP:PYR) regioisomeric ratio (r.r.), underscoring the challenge of favouring six-membered ring formation using existing natural scaffolds (Fig. 1D, S3B).

To achieve the desired high regioselectivity, we used APN to design novel scaffolds containing a histidine-coordinated heme. We designed three subsets, modelled with either the nitrene intermediate or one of the two divergent C–H amination products (Fig. S3C).

Using metrics from AF3 predictions, we selected 86 novel-scaffold designs for experimental validation. Each design subset successfully delivered at least one enzyme with activity exceeding the functional threshold (Fig. S3D). While designs targeted for nitrene formation largely favoured the pyrrolidine, the top-performing designs for each product successfully recapitulated the intended regioselectivity (Fig. S3D): GDM_NT_141, designed as a pyrrolidine synthase, yielded a 4:96 PIP:PYR ratio, while GDM_NT_121, designed as a piperidine synthase, achieved up to 94:6 PIP:PYR (Fig. 1C, S4A). In a second iteration of 90 novel scaffolds targeted specifically for piperidine synthesis, five designs exceeded the functional threshold, with all exhibiting high regioselectivity (Fig. 1D, S3F). The lead design, GDM_NT_270, delivered the piperidine product with 22 turnovers, 99:1 r.r., and 94% enantiomeric excess (e.e.). Compared to *Tam*Pgb wt, this represents an improvement of two orders of magnitude in total turnover for the piperidine product through complementary regioselectivity (Fig. S4A). Our success in designing piperidine-selective aminases provides orthogonal starting points to natural pyrrolidine-selective protoglobins. By accessing new catalytic properties such as novel regioselectivity, we anticipate APN will expand the chemical space accessible to biocatalysis.

### Design of serine esterases for DEHP hydrolysis

While piperidine synthesis involves new-to-nature chemistry, *de novo* design may also expand the substrate scope of known enzymatic mechanisms. To test this, we designed hydrolases for the partial degradation of DEHP, a ubiquitous plasticiser, pervasive environmental contaminant^30^, and endocrine disruptor linked to significant developmental and reproductive toxicity^36^ (Fig. 1E). A small number of DEHPases have been discovered in nature, many of which rely on a serine catalytic triad mechanism to hydrolyse one or both esters on the molecule^37–42^. However, despite the ubiquity of natural hydrolases, DEHPase activity is very rare due to the steric bulk of the 2-ethylhexyl side chains. Furthermore, DEHP is nearly insoluble in water (solubility ∼0.3 mg/L)^43^, and the organic cosolvents that improve its accessibility inactivate most natural esterases^44,45^.

We derived catalytic motifs containing Ser-His-Asp triads and oxyanion hole residues from natural hydrolase crystal structures (Table S9), and modelled DEHP conformers generated by RDKit^46^ into each motif as tetrahedral intermediates covalently linked to the nucleophilic serine (Fig. 1F). Following recently established design principles for serine esterases^17^, we used structure predictions of five catalytic states to select designs for testing (Supplementary Information). To assay DEHP degradation, we monitored formation of MEHP, the single-hydrolysis product of DEHP (Fig. 1E). In an initial round of 81 designs, four had activity three standard deviations above background and one design exhibited lysate activity of at least 2% of QHH21706.1 ("QHH"), a natural hydrolase known to degrade DEHP^41,47^ (Fig. S5A). Over two additional rounds we improved the novel-scaffold hit rate to 11% (activity exceeding 2% of QHH) and the recycled-scaffold hit rate to 39% (Fig. 1G, S5A). For comparison, a recent screen of 65 natural esterases (16 of which were predicted to have DEHPase activity) only found one active DEHPase^41^, highlighting the challenge of non-natural substrates even for enzymes whose core catalytic machinery is evolutionarily optimised.

Our hits represent seven new structural families able to perform this reaction that are distinct from each other (pairwise TM-score < 0.5) and from known natural esterases (see below).

To further characterise the top hits, we measured the activity of purified enzymes. At room temperature in aqueous buffer, our designs were less active than natural DEHP esterases, with the most-active design GDM_DEHP_378 reaching 6% of the MEHP production of QHH after 4 hours and 26% of Q7SIG1^41^, another natural DEHPase (or 13% and 34% respectively after 24 hours) (Fig. S5B). However, the designs possess stability properties rare in the natural enzyme repertoire. The most robust design, GDM_DEHP_376, is 14-fold more active at 90 °C than at room temperature (Fig. 1H, S5C) and 2-fold more active in 75% acetonitrile than at 0% (Fig. S5B); by contrast, both natural esterases, including thermophile-derived Q7SIG1, are completely inactive under either condition (Fig. 1H). While some natural enzymes tolerate moderate (<50%) or near-anhydrous organic solvent^48,49^, activity at high concentrations of polar, water-miscible solvents like acetonitrile is rarely reported^45,50^.

These conditions are uniquely denaturing because the solvent simultaneously interacts with the hydrophobic core and displaces stabilising water at the protein surface^48^.

The designs also have favourable production properties: GDM_DEHP_376, with 191 amino acids, is 60% and 40% smaller than QHH and Q7SIG1, respectively, and therefore has a higher mass density of active sites. Our designs also have higher expression in *E. coli* than the natural esterases (Fig. S5F), likely reducing the cost of enzyme production. Increased activity and full degradation of DEHP to phthalic acid^51^ are still needed before our designs can be used in real-world bioremediation, but their small size, high expression, and unusually high temperature and solvent stability already make them attractive starting points for optimisation. More broadly, their activity in 75% acetonitrile shows that *de novo* design can provide scaffold properties rare in nature, even for common reaction mechanisms.

### Designing enzymes for model reactions

During the development of APN we carried out design campaigns on three model reactions representing diverse design challenges (Fig. 2, Table 1):

● Kemp elimination, a single-step, unimolecular ring opening of the substrate 5-nitrobenzisoxazole (5-NBO), commonly used as a mechanistically simple enzyme-design benchmark^52–55^ (Fig. 2A).
● Serine ester hydrolysis of 4-methylumbelliferyl acetate (4MU-Ac), a multi-step, unimolecular reaction requiring precise molecular control in multiple conformational states^17,56^ (Fig. 2B). We developed our methods on 4MU-Ac alongside the work on DEHP, as 4MU-Ac hydrolysis can be measured easily for large numbers of designs. Below, “serine esterase” refers to enzymes designed and tested against 4MU-Ac.
● Styrene cyclopropanation via carbene transfer ^19,21,57^, which requires stereoselective coordination of a noncovalent heme cofactor, styrene, and ethyl diazoacetate (Fig. 2I).

**Figure 2.**
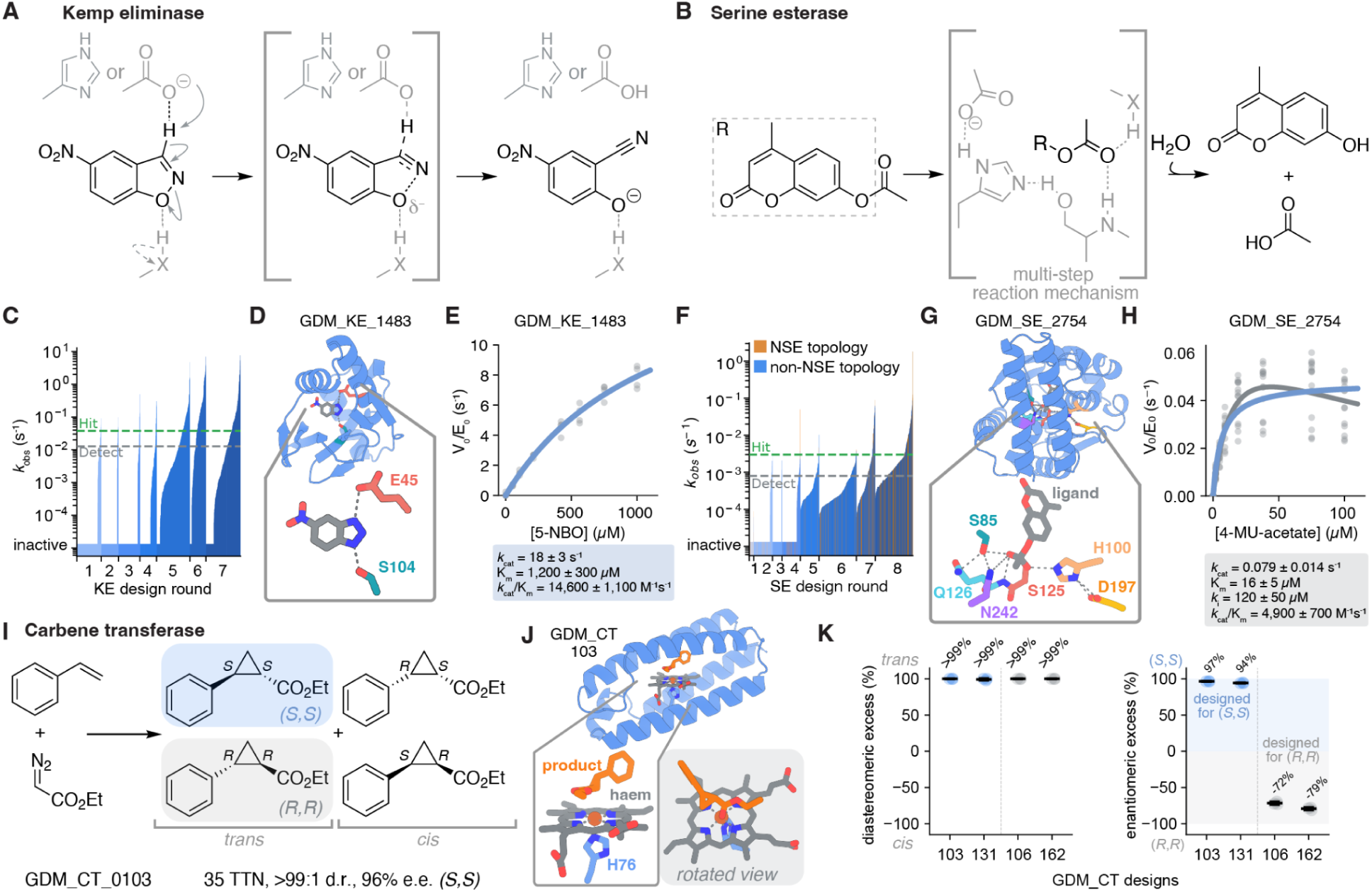
Designed enzymes for model reactions. Reaction schemes for A) Kemp elimination, B) serine-triad-based 4MU-Ac hydrolysis, and I) styrene cyclopropanation via carbene transfer with preparative scale results. Measured *k*_obs_ values across all tested C) Kemp eliminase and F) serine esterase designs, with hit-calling threshold and detection limit overlaid (0.038 s^-1^ and 0.0127 s^-1^ for Kemp eliminase and 0.003 s^-1^ and 0.0008 s^-1^ for serine esterase, respectively; see Supplemental Information). *k*_obs_ values were set to a floor of 10^-5^ for designs which are known to be inactive. D, G, J) AF3 predictions of selected active designs alongside (E, H) Michaelis-Menten kinetics for the Kemp eliminase and serine esterase, with errors representing standard deviation of four technical replicates and 14 technical replicates across two biological replicates, respectively. Blue fits are derived from classical Michaelis-Menten kinetics, whereas grey fits include substrate inhibition. K) LC-MS characterisation of styrene cyclopropanation activity in technical triplicate showing diastereoselectivity and enantioselectivity.

For Kemp elimination and serine ester hydrolysis, we iteratively improved filtering approaches over multiple rounds of design, with each round containing multiple “facets” (subsets) of designs from the same pipeline configuration (Fig. S7). The diffusion model *in silico* pass rates (frequency with which designs pass a fixed set of filters) also improved, but this did not measurably affect experimental outcomes when keeping other factors fixed (Fig. S8), so we focus here on the effect of filtering on design success.

We quantified catalytic activity by purifying enzymes from *E. coli* and measuring *k*_obs_, or the per-enzyme rate of product formation at a fixed substrate concentration (1 mM 5-NBO or 100 µM 4MU-Ac) (Fig. 2C, F, S6A, S7). For all reactions, >95% of designs were well-expressed as assessed by SDS-PAGE or SEC. Our *k*_obs_ measurements were highly reproducible (R^2^>0.84) between independent experiments (Fig. S6E). For the best designs, we also determined kinetic parameters *k*_cat_ and K_m_ (Fig. S6B). To compare across rounds and facets, we computed hit rates, defined as the fraction of designs exhibiting multiple-turnover kinetics and a *k*_obs_ exceeding 0.038 s^-1^ or 0.003 s^-1^ for Kemp eliminase and serine esterase, respectively (Fig. S6C, D, F). Note that these *k*_obs_ thresholds aim to identify strongly active designs and are well above the detection limit of our assays (Fig. 2C, F; Methods).

We tested a total of 1,536 novel-scaffold Kemp eliminases over seven rounds of design using previously published theozyme motifs^52^. Hit rates ranged from 3% in the first round to 34% in the best facet (Fig. S7A). The best design, GDM_KE_1483, had a *k*_cat_/K_m_ of 14,600 M^-1^s^-1^ and a *k*_cat_ of 18 s^-1^ at pH 7.5 (Fig. 2D-E, S9A, S14B, Table 1, S1). This is 70-fold higher than the best previously published novel-scaffold *k*_cat_/K_m_ values^53,55,58^, although 10- to 100-fold lower than that of experimentally optimised designs (Table S1). We also evaluated Kemp eliminases at elevated pH (9 and 10, instead of 7.5), observing increases in *k*_cat_ up to 30 s^-1^ and *k*_cat_/K_m_ up to 45,000 M^-1^s^-1^ (Fig. S10). In our final and best round, 5/90 novel-scaffold designs had a *k*_obs_ > 1.0 s^-1^ (Fig. S7A), achieving a fraction of 6% designs exceeding the best previously published novel-scaffold catalytic rates (Table S1). Across all design rounds, we measured 24 Kemp eliminase designs with *k*_cat_/K_m_ > 1,000 M^-1^s^-1^ and 11 with *k*_cat_/K_m_ > 5,000 M^-1^s^-1^. Table 1 summarises our model reaction results with further catalytic parameters of top designs and comparison to literature in Table S1.

We tested a total of 1,569 novel-scaffold serine esterases over eight rounds of design, with hit rates ranging from 0% in the first round to 44% in the best facet (Table 1, Fig. S7B). The best design GDM_SE_2937 had a *k*_cat_/K_m_ of 360,000 ± 70,000 M^-1^s^-1^ and a *k*_cat_ of 5.2 ± 1.3 s^-1^ (Table 1, Fig. S9B, S14A), exceeding even the most active previous *de novo* designs that have been iteratively optimised with experimental feedback (Table S1)^17^. However, this design has a similar scaffold and active-site placement to natural serine esterases in the Protein Data Bank (PDB)^59^ (Fig. S13; Supplementary methods). To investigate this further, we categorised a design as having natural serine esterase topology (NSE topology) if it had a TM-score > 0.45 and scaffold-aligned triad RMSD < 5 Å to any annotated serine hydrolase in the PDB. NSE topology designs constitute 16.4% of our novel-scaffold serine esterases overall (Fig. S11), and six of our top seven designs by *k*_cat_/K_m_ (Fig. S12). Our best non-NSE topology design had a *k*_cat_/K_m_ of 4,900 ± 700 M^-1^s^-1^ and a *k*_cat_ of 0.08 ± 0.014 s^-1^ (Fig. 2G-H, S9C, Table 1, S1). This *k*_cat_/K_m_ is still roughly 4-fold higher than the best previous novel-scaffold designs for 4MU-Ac or the related 4MU-PA (4MU phenylacetate) (Table S1)^17^. Our top DEHPase designs all had non-NSE topologies.

For styrene cyclopropanation via carbene transfer, we tested 91 novel-scaffold designs created from previously published theozyme motifs^21^ including a heme, axial histidine, and transition states for either of two *trans* (*R,R* and *S,S*) product enantiomers (Fig. 2J, S15A), which complements recent work targeting the *cis* products^23^. 77 (85%) of these designs bound heme, from which we selected 48 designs (24 designed for each enantiomer) for chiral analysis. Of these designs, the four most-selective designs for each enantiomer all had the enantioselectivity intended by design (Fig. S15B). Our best designs had near-perfect diastereoselectivity (diastereomeric ratio > 99%) and high enantioselectivity with 96% e.e. for the *S,S* product and 79% e.e. for the *R,R* product (Fig. S15C-D). This is comparable to the 72% and 94% e.e. (for *R,R* and *S,S* products, respectively) achieved in two rounds of design by previous work using the same input motifs^21^. Another study^19^ targeted the *S,S* enantiomer using a synthetic porphyrin cofactor and achieved 82% e.e. by redesigning a scaffold known to bind porphyrins. Unlike these previous studies, we achieved cofactor-binding, target activity, and stereoselectivity in a single round of design with no intermediate experimental feedback.

### Generalised structure-sequence ensemble scoring increases design success

A key goal of our model reaction work was to discover new ways to evaluate designs *in silico* to obtain higher hit rates and activities. Since our design process is driven by a structural hypothesis in the form of a catalytic motif and ligand pose, we evaluated our designs based on AF3-predicted structures of apo and ligand-bound states across the catalytic cycle ("multi-state"). To assess pre-organisation, or the robustness with which an enzyme forms key catalytic interactions, we generated ensembles of AF3 predictions with different random seeds ("multi-seed")^27,60,61^. We also introduced a generalisation of multi-seed scoring which evaluates a designed backbone via an ensemble of LigandMPNN-derived sequences ("multi-resequence"). Multi-resequence scoring greatly amplifies the predictive power of multi-state, multi-seed filters and was the most important contributor to our high success rates and activities in later rounds.

All of our filtering strategies make use of activity-predicting metrics that we developed based on biochemical principles. In Kemp elimination, preorganisation of the catalytic base for proton abstraction from the substrate is known to be crucial for catalysis^62^. Consistent with this, we observed that by retrospectively filtering designs on the AF3-predicted geometry between the transition-state analog 6-nitrobenzotriazole (6-NBT) and the catalytic base or oxyanion hole, we could increase hit rate by 2.7-fold at 50% recall (Fig. 3A-B, S19A). Prospectively filtering on these geometries in a subsequent round of design increased hit rate by 3-fold and best k_obs_ from 0.49 s^-1^ to 1.38 s^-1^ (Fig. 3B). For DEHPases and 4MU-Ac serine esterases, the consistency of the ligand pose between enzyme-substrate (ES) and tetrahedral intermediate (TI1) predictions correlates to *k*_obs_ (Spearman *ρ*=-0.19; Fig. 3C-D, S24) in retrospective data. This could indicate pre-organisation of the substrate ester for nucleophilic attack by the catalytic serine^61^, or more general competence for substrate binding. Additional predictive metrics include hydrophobic burial of the Kemp eliminase catalytic base, polar interactions in the serine esterase triad, and near-attack positions of the bond-forming carbon and nitrogen atoms in piperidine synthases (Fig. S22-23, S29; Supplementary Notes). The discriminative signal of these metrics across hundreds of novel-scaffold designs spanning 4 reactions indicates that AF3 predictions offer a direct and general way to relate the atom-level geometry of designs to real-world catalytic outcomes.

**Figure 3.**
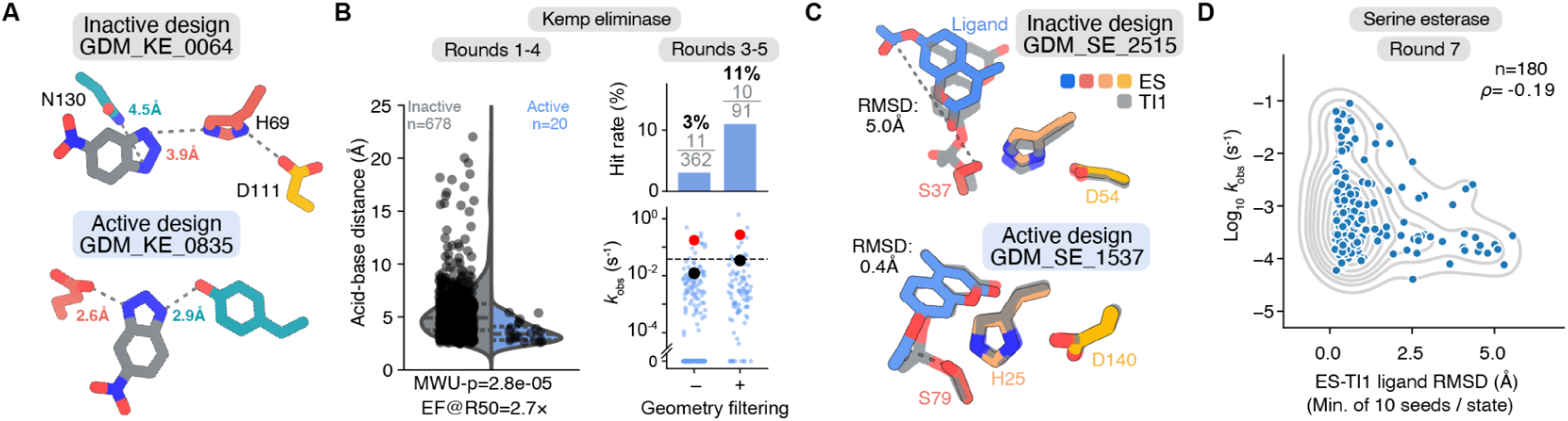
Filters based on AlphaFold structure prediction. A) Example Kemp eliminase design active sites, showing positions of catalytic base (red), hydrogen bond donor (teal), and transition state analog (grey). B) Left: Distribution of mean distance between catalytic base and corresponding N3 of 6-NBT across seven structure prediction seeds, split by hit status, for designs which were not filtered on this metric. Mann-Whitney U statistic=10503; “EF@R50": precision enrichment factor at 50% recall. Right: Hit rates and reaction rates *k*_obs_ for designs prospectively filtered on catalytic base and 6-NBT geometry in design round 5 (+), compared to baseline designs not filtered on this metric but matched in other design variables from rounds 3-4 (-). Black dots: mean *k*_obs_ of all designs. Red dots: mean *k*_obs_ of only hits. C) Example serine esterase design active sites with ES (colours) and TI1 (grey) predictions aligned on catalytic residues. When ES-TI1 RMSD is low, there is also a shorter distance between the serine O*_γ_* nucleophile and the substrate carbon electrophile (dotted line). D) Log_10_ k_obs_ versus minimum ligand RMSD between an ES and TI1 prediction across 10 seeds of each state. Dataset size n and Spearman *ρ* are shown. Only data from design round seven is shown because earlier rounds yielded few high k_obs_ values and later rounds were filtered on this metric. See additional examples of biochemically inspired filters in Supplementary Notes and details of design rounds in Fig. S7 and Supplementary Methods.

Given the effectiveness of multi-seed scoring (Fig. S25)^17,27^, we asked whether discrimination could be improved further by considering not only ensembles of predicted structures but also of sequences derived from the same backbone. In multi-resequence (MR) scoring, we evaluated multiple LigandMPNN-designed sequences ("resequences") for each backbone and required each resequence to pass the same set of filtering metrics (Fig. 4A, Methods). In effect, MR scoring evaluates whether key catalytic interactions are preorganized (and AF3 can make confident predictions) based on the backbone alone, not just in the context of a particular sequence.

**Figure 4.**
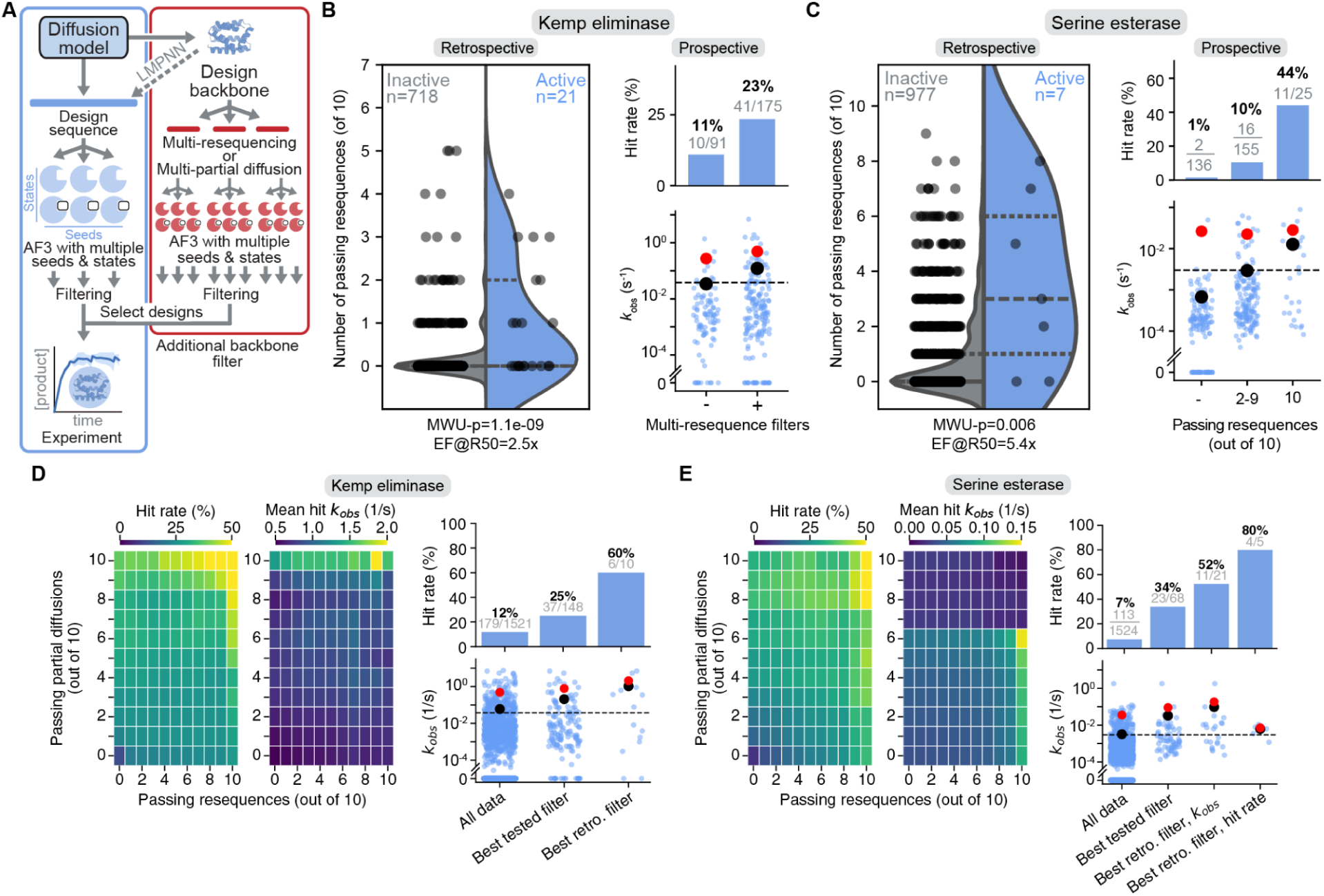
Sequence ensemble-based scoring. A) Schematic of multi-resequence and multi-partial-diffusion scoring. B) Left: Retrospective number of resequences passing filters versus hit status for Kemp eliminases (rounds 1-4, before MR filters were added). Mann-Whitney U (MWU) statistic= 4514.5. “EF@R50": precision enrichment factor at 50% recall. Right: Prospective hit rates and k_obs_ values versus multi-resequence filtering. “-": MR filters were not used (round 5, it is unlikely, but still possible, for designs to have some passing resequences); “+": designs from round 5 and 6 selected for at least 1 of 7 resequences passing, and usually 7/7. Gray fractions indicate the number of hits and total designs in each condition. Red dots indicate the mean *k*_obs_ of all hits, whereas black dots indicate the mean *k*_obs_ of all samples. C) Same as (B) but for Serine esterases. Retrospective data from rounds 2-6 (left) and prospective data from round 7 (right). Mann-Whitney U (MWU) statistic= 1652.5. More details on design rounds in Fig. S7 and Supplementary methods. D) Left: Retrospective hit rates and mean hit k_obs_ across all Kemp eliminase designs, versus the number (or more) of their resequences or partial diffusions passing filters out of 10 trials. Right: Hit rates (with number of hits and total designs in gray) and k_obs_ distributions (with overall mean as black dots and mean of hits as red dots) for selected filtering conditions. “Best tested filter” is the best filter that was tested prospectively as shown in (B) but applied retrospectively here to all data: 7 passing resequences out of 7 trials (denoted as N_MR_ = 7/7), no scoring of multi-partial diffusion (N_MPD_ ≥ 0/10). “Best retro. filter” is the best filtering condition discovered in this analysis: N_MR_ = 10/10, N_MPD_ = 10/10. E) Same as (D) but for serine esterases. The best retrospective filters differ for hit rate and mean k_obs_ and are shown separately. Best tested filter is N_MR_ = 10/10, N_MPD_ ≥ 0/10; best retrospective hit rate filter is N_MR_ = 10/10, N_MPD_ = 10/10; best retrospective k_obs_ filter is N_MR_ = 10/10, N_MPD_ ≥ 6/10. Note that (B) and (C) analyse exactly the design rounds before and after introducing multi-resequencing filters, while (D) and (E) retrospectively analyse all designs, hence hit rates and design numbers differ slightly for “best tested filter”. See additional analyses in Figure S28 and Supplementary Notes.

When we retrospectively thresholded at 50% recall on the number of resequences passing filters, we found that Kemp elimination and serine esterase hit rates increased by 2.5-fold and 5.4-fold, respectively (Fig. 4B, C). In subsequent prospective design rounds, Kemp eliminase designs filtered by MR had a hit rate of 23% or 2-fold higher than designs that were not MR-filtered (Fig. 4B). Serine esterases filtered by MR had hit rates of up to 44%, or 30-fold higher than without MR filtering (Fig. 4C). The benefits of multi-seed and MR scoring can be combined; increasing the number of required passes for both seeds and resequences leads to retrospective hit rates for serine esterases as high as 56% (5/9 designs) (Fig. S25B). Finally, MR filtering leads to better starting points or “parents” for further optimization. When we tested multiple LigandMPNN resequences of known active parent designs, we found that resequences of designs passing MR filters were more likely to improve on their parent than resequences of parents not passing MR filters (Fig. S26).

Our filter improvement efforts revealed a number of limitations of our pipeline and of *de novo* design more generally. Specifying motifs and filters for new target reactions requires reaction-specific mechanistic understanding and trial-and-error. The input motif in particular has a strong influence on design outcome that is not captured by our filters (Fig. S7B, Table S2). Similarly, although MR scoring is a promising general strategy that proved effective on multiple reactions we tested, it depends on having already established reaction-specific metrics. Encouragingly, we found that general AF3 confidence and consistency metrics can enrich for hits, providing a reaction-agnostic complement to mechanism-based filters (Fig. S29C). Finally, our success on the first rounds of piperidine synthase and DEHPase design was partly due to insights from model reaction work, suggesting that filters are transferrable between reactions.

### Sequence-structure co-generation for design and filtering

The results above are all for designs that have undergone a LigandMPNN resequencing step after structure generation by the AP Novo diffusion model. However, this step is not essential because the diffusion model co-generates sequence alongside structure, similar to other recent models^13,15,63^. It achieves this by concurrently running a Gaussian diffusion process for structure coordinates and masked discrete diffusion for the amino acid sequence (Supplementary Methods). The diffusion-model-generated sequences ("AP Novo sequences") have similar *in silico* pass rates to LigandMPNN sequences (Fig. S2G), as well as comparable experimental activities for Kemp eliminase and serine esterase designs (Fig. S28B, Supplementary Notes).

Despite having similar *in silico* and experimental performance to LigandMPNN, AP Novo sequences exhibit a distinct amino acid composition (Fig. S27A). This led us to hypothesise that AP Novo sequences could increase the discriminating power of sequence-ensemble-based filtering. Indeed, across the LigandMPNN sequences tested above, requiring their corresponding AP Novo sequence (which was not experimentally tested) to also pass filters increased retrospective hit rate from 29% to 45% for Kemp eliminases and from 5% to 15% for serine esterases (Fig. S28A). We also confirmed this effect prospectively: in a new round of MR-filtered LigandMPNN sequences, having a non-passing versus passing AP Novo sequence led to a hit rate increase of 21% to 37% for Kemp eliminases and 20% to 43% for serine esterases (Fig. S28A).

These results suggest that the AP Novo diffusion model can itself be used for MR-style filtering. To test this, we used the model to run “partial diffusion” - adding a small amount of noise to an existing design and re-running the denoising process to completion (Supplementary Methods). This yields a modified sequence with an almost identical backbone, an outcome similar to LigandMPNN resequencing. For a given design, we generated multi-partial-diffusion sequences and applied filters to each ("MPD” filtering). We found that MPD-filtered AP Novo sequences have similar hit rates and k_obs_ activity distributions as MR-filtered LigandMPNN sequences (Fig. S28B, Supplementary Notes).

More strikingly, MR and MPD filters can be combined synergistically. Across all of our Kemp eliminase and serine esterase designs, retrospectively filtering to those passing both 10/10 MPD filters (i.e. 10 partial diffusion sequences are assessed and all pass filters) and 10/10 MR filters leads to hit rates of 60% and 80% respectively, the highest in all of our analyses (Fig. 4D-E). Even higher precision, up to 100%, can be achieved if filters are tightened until fewer than five designs pass (Fig. S30). Combined MR and MPD filtering also selects for higher-k_obs_ hits, although hit rate and activity are maximised under slightly different filter settings (Fig. 4D-E, S30). Overall, these results show that in addition to scoring sequence ensembles, it is also advantageous to ensemble over different sequence-design models.

Our MR and MPD results may provide a unified explanation for the success of recent design pipelines based on iterative redesign.^17,18,24,25,64,65^ These approaches sample an initial set of designs and then apply rounds of resequencing or partial diffusion followed by further filtering. The goal is to obtain designs passing stringent filters in a more computationally efficient manner than non-iterative sampling. However, a separate, potentially more important effect is that designs implicitly undergo a process akin to MR or MPD scoring by having their backbones (with small variations) repeatedly pass filtering stages.

Here we used non-iterative sampling and directly compared novel-scaffold designs whose backbones pass or fail MR/MPD filters, while the design sequences all pass the same sequence-level filters. Unlike iterative pipelines, which conflate selection for backbone quality with improvements in sequence quality across rounds, this experimental design isolates the contribution of backbone quality and shows that it plays an outsized role in experimental success. Therefore, our results argue for evaluating methods not only by individual designed sequences, but also the quality of their backbones via ensemble scoring.

The “backbone quality” measured by the MR pass rate can be interpreted as the marginal likelihood under the resequencer model that a given backbone satisfies filters, or intuitively, the resequencer-accessible volume of functional sequence space for that backbone (Supplementary Methods). MPD scoring extends this principle by also exploring the local structural neighborhood, selecting for backbones having broadly accessible functional basins in joint sequence-structure space. Seen in this light, iterative pipelines implicitly select for backbones with larger functional basins. A similar effect could also be achieved by sampling methods such as beam search and Feynman-Kac correctors, which likewise perform selection over an ensemble of design trajectories^15,63,66^.

### Biochemical and bioinformatic characterisation of designs

To verify that the designs have activity via the intended catalytic interactions, we mutated active-site residues in selected Kemp eliminases and serine esterases (Fig. 5A-B, S16). Five of 10 mutants across four Kemp eliminase designs had abolished activity and the remaining mutants exhibited reduced activity (Fig. 5A, S16A). Additionally, across four serine esterase designs, activity was reduced in all mutants, and abolished in 14 of 19 mutants, including in 100% of mutations targeting the catalytic triad (Fig. 5B, S16B).

**Figure 5.**
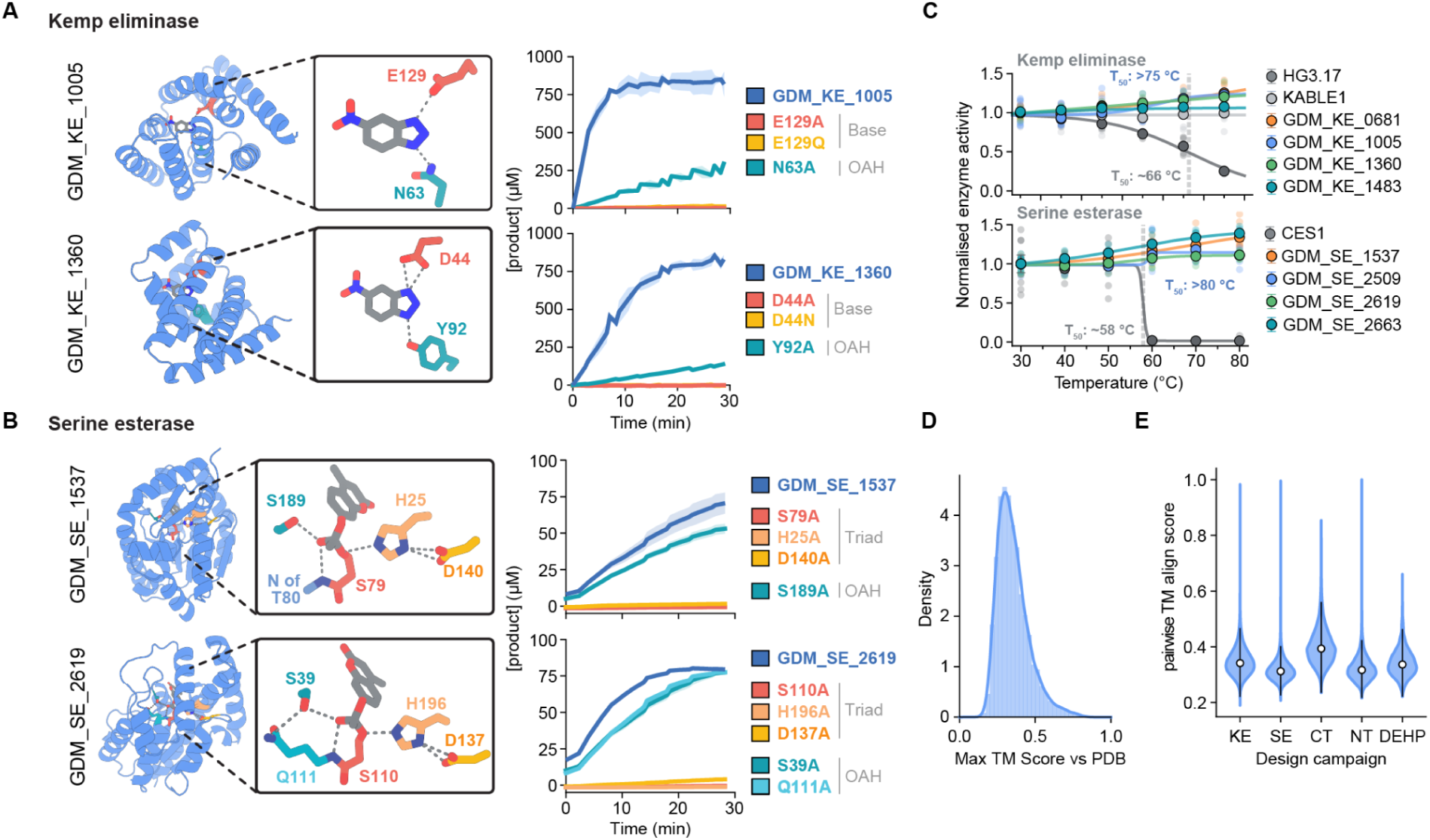
Active site mutagenesis, thermostability, and scaffold novelty and diversity. A) Catalytic residue knockouts of Kemp eliminases. “Base": catalytic base. “OAH": oxyanion hole (hydrogen bond donor). B) Catalytic residue knockouts of serine esterases. “Triad": serine catalytic triad. C) Thermostability of designs versus previously published proteins. Residual enzyme activity was assayed after 10 minutes at indicated temperatures and cooling to standard conditions. T_50_ values, the temperature at which 50% activity is lost, exceeded 80 °C for all designs. D) Max TM-score of all designs when compared to the PDB. E) Pairwise inter-design TM-scores, by campaign.

To assess thermostability, an important property for practical applications and a determinant for evolvability^67^, we assayed the activity of Kemp eliminases and serine esterases after incubation for 10 minutes at temperatures up to 80 °C, followed by 10 minutes at room temperature. All designs tested either maintained or slightly increased activity after heat treatment, while the natural esterase human carboxylesterase 1 (CES1) lost all activity after heating above 60 °C and the previously reported Kemp eliminase, HG3.17^62^, lost most of its activity by 75 °C (Fig. 5C).

With the exception of the NSE topology serine esterases discussed above, our designs are structurally novel (Fig. S17), with a median template-modeling-score (TM-score) of 0.33 to the best match in the PDB across all novel-scaffold designs (Fig. 5D). They are also structurally diverse (Fig. S18), with a mean of campaign-wise median inter-design TM-score of 0.34 (Fig. 5E), and a large fraction of designs falling into distinct structural clusters with TM-scores < 0.5 relative to each other (Table 1). Among our most active designs, the maximum sequence identity to a natural sequence across the Uniref, MGnify, and Small BFD databases is 29% (Table S3). Notably, even the NSE topology serine esterases generally have extremely low or undetectable sequence homology (<20%) with natural sequences and can be predicted with high confidence by AF3 without a multiple sequence alignment. This suggests that these designs occupy a middle ground between traditional *de novo* designs and “natural-like” proteins from sequence-based generative models^68–70^.

### Conclusions

These results show that *de novo* enzyme design has matured beyond proof-of-concept into a source of biocatalysts for real-world problems where natural enzyme mining has had limited success. Our nitrene transferases have unprecedented regioselectivity in piperidine synthesis, making them preferable to natural sequences as starting points for engineering. Additionally, our DEHPase designs expand the pool of candidate enzymes that can be developed for DEHP degradation. These examples underscore a key advantage of *de novo* design: the ability to tailor active sites to effect specific, challenging chemical outcomes, particularly when interacting with molecules that natural enzymes struggle to accommodate.

We applied multiple design-test-learn cycles on model reactions to uncover general design principles, in the process obtaining state-of-the art catalytic parameters. This effort revealed that applying filters to sequence ensembles derived from the same backbone greatly amplifies their discriminating power. This helps to explain the previously observed effectiveness of iterative design pipelines and highlights the role played by the protein backbone in catalytic pre-organisation. We also show that the combination of design and structure prediction has sufficient accuracy to test detailed mechanistic hypotheses and thereby increase design success rates. We envision that this approach of “understanding by creating”, a long-held tenet of computational design^4^, can now be applied at scale for a wide range of reactions and fundamental questions. Finally, we showed that APN can directly co-generate structures and sequences of active enzymes without the need for a separate sequence-design model, removing a layer of complexity from the design process and opening up the possibility of applying post-training techniques to optimise final design sequences directly.

While this is exciting progress, limitations remain. The initial stages of design require significant expertise to construct motifs, which influence design outcomes in a way that is not captured by current filters. Many samples must be generated for a given motif in order to find a design that passes filters, which can be practically prohibitive. While evaluating sequence ensembles is a significant step in assessing backbone quality, the problem of choosing an optimal backbone, and furthermore an optimal sequence for a given backbone, remains far from solved. Most importantly, the catalytic activities of *de novo* designs are still orders of magnitude lower than natural enzymes^26,71^ or those optimised by directed evolution20,62,72,73.

All of these limitations indicate that our models do not meaningfully model the physics of catalysis. Addressing this will require progress in incorporating physical understanding into design pipelines, or machine-learning approaches to better extract the determinants of function from natural protein data. Coupled with increases in gene synthesis and assay throughput to unlock further data-driven insights, we are optimistic that *de novo* enzyme design will rapidly become a central tool for tackling chemistry’s greatest challenges.

## Methods

Full details, including experimental methods, are available in the Supplementary Methods.

### AlphaProtein Novo design pipeline

The AP Novo pipeline designs enzymes by motif scaffolding followed by filtering of candidate designs (Fig. 1A, Fig. S1)^7,8^. For each reaction, we constructed catalytic motifs comprising sidechain functional groups and small molecule ligands based on a hypothesised mechanism. We input each motif into a diffusion model to generate a structure holding the motif in the specified conformation ("scaffold” the motif), and input the structure into LigandMPNN^74^ to obtain a final designed sequence ("resequencing"). A subset of experiments also tested sequences generated directly by the diffusion model, without resequencing. To select designs for experimental testing, we filtered on metrics derived from structure predictions from AlphaFold 3 (AF3)^5^ or a fine-tuned variant (see Supplementary Information for details of where each model was used). Full details of the AlphaProtein Novo design pipeline, including *in silico* benchmark performance and details of model architecture and training, are available in the Supplementary Methods.

### Sequence Ensemble Scoring

For Kemp eliminases, we consider a MR pass as a backbone that passes at least 6 out of 7 resequences. For serine esterases, we consider a success as 10 out of 10 resequences passing. For MPD, for both Kemp eliminases and serine esterases, partial diffusion was constrained such that the average sequence identity to the parent was roughly 88% and the RMSD to the parent was roughly 0.5 Å (Fig. S27C), and we consider a success as 9 out of 10 MPD sequences passing.

### Structural Novelty and Diversity

To analyse the structural novelty of our novel-scaffold designs (Fig. 5), we used Foldseek^75^ to compute the maximum TM-score for each design against any structure in the PDB^59^. We used the default alignment mode (3Di+AA Gotoh-Smith-Waterman local alignment), an e-value threshold of 10^10^, and standard defaults for all other parameters. For each Foldseek hit, we extracted the `qtmscore` (query-length-normalised) and `ttmscore` (target-length-normalised) values and averaged them to obtain the TM-score. We were able to obtain at least one Foldseek hit for each novel-scaffold design, and report the maximum TM-score across all Foldseek hits. Recycled-scaffold designs, as well as the first round of serine esterase designs (which did not result in any active hits), were excluded from this analysis. To analyse diversity, we used TM-align^76^ to compute the TM-score between each pair of novel-scaffold designs for each reaction. We found a median pairwise TM-score of 0.34 and a 99th percentile of 0.48. In rare cases we observed pairs of designs with TM-score close to 1, possibly due to coincidental repetition of random seeds for the diffusion model. In later design rounds we filtered designs to avoid repetition of highly similar structures.

### Natural Serine Esterase Topology

We defined natural serine hydrolases as PDB entries annotated with an Enzyme Commission^77^ (EC) prefix of 3.1.1 or 3.4.21. To calculate the TM-score, we used TM-align^76^ and reported the TM-score normalised to the length of the query design. To calculate the triad RMSD, we computed the RMSD over all heavy atoms (both backbone and sidechain) of the catalytic triad. To locate the corresponding triad residues within the PDB structures, we employed a geometric motif-matching approach as follows. Because the target structures lack pre-assigned motif residue indices, we matched residues from the reference motif to the target structure by constraining the search to corresponding sequence tokens and atom names. We then applied the Hungarian algorithm to determine the optimal residue assignment by minimising the distances between matched atoms, similar to the approach used in RFdiffusion2^7^ to identify unindexed motif residues.

### Model availability

The code and weights for the AlphaProtein Novo diffusion model and AF3 variant are available freely for non-commercial use at https://github.com/google-deepmind/alphaprotein-novo.

### Data availability

All designed sequences from this study, along with predicted structures and metadata, will be made publicly available upon publication.

All scientific datasets used to create training and evaluation inputs are freely available from public sources. Structures from the PDB were used for training and evaluation of the diffusion model and AF3 variant.

Furthermore, https://files.wwpdb.org/pub/pdb/data/assemblies/mmCIF/, and a structure distillation set created from UniRef90 v.2021_04 (https://ftp.ebi.ac.uk/pub/databases/uniprot/previous_releases/release-2021_04/uniref/), was also used for training; for sequence clusters see https://cdn.rcsb.org/resources/sequence/clusters/clusters-by-entity-40.txt; for sequence data see https://files.wwpdb.org/pub/pdb/derived_data/) as well as the Chemical Components Dictionary (https://www.wwpdb.org/data/ccd).

We also used the following publicly available databases for training or evaluation of the AF3 variant, or in the creation of the structure distillation set.

UniRef90 v.2020_01 (https://ftp.ebi.ac.uk/pub/databases/uniprot/previous_releases/release-2020_01/uniref/), UniRef90 v.2020_03 (https://ftp.ebi.ac.uk/pub/databases/uniprot/previous_releases/release-2020_03/uniref/), UniRef90 v.2021_04 (https://ftp.ebi.ac.uk/pub/databases/uniprot/previous_releases/release-2021_04/uniref/), UniRef90 v.2022_05 https://ftp.ebi.ac.uk/pub/databases/uniprot/previous_releases/release-2022_05/uniref/), Uniclust30 v.2018_08 (https://wwwuser.gwdg.de/∼compbiol/uniclust/2018_08/), Uniclust30 v.2021_03 (https://wwwuser.gwdg.de/∼compbiol/uniclust/2021_03/), MGnify clusters v.2018_12 (https://ftp.ebi.ac.uk/pub/databases/metagenomics/peptide_database/2018_12/), MGnify clusters v.2022_05 (https://ftp.ebi.ac.uk/pub/databases/metagenomics/peptide_database/2022_05/), UniProt v.2021_04 (https://ftp.ebi.ac.uk/pub/databases/uniprot/previous_releases/release-2021_04/knowledgebase/), BFD (https://bfd.mmseqs.com), RFam v.14.9 (https://ftp.ebi.ac.uk/pub/databases/Rfam/14.9/), RNAcentral v.21.0 (https://ftp.ebi.ac.uk/pub/databases/RNAcentral/releases/21.0/),

## Acknowledgements

We thank C. Beattie, R. Broomfield-Tagg, A. Ceccarelli, A. Cowie, R. Dooley, A. Doucet, J. Dunger, N. Elmashky, L. Hall, N. Jean, J. Jenkins, J. Jumper, O. Kovalevskiy, O. Lange, C. Langstaff, M. Levine, V. Martin, N. Monck, I. Moraes, L. Petruska, A. Pita, K. Rajasekar, Y. Righetti, D. Saxton, N. Sultana, F. Tian, Z. Wang, R. Ward, and the Structural Biology Science Technology Platform (STP) at the Francis Crick Institute for valuable discussions and technical support.

## Author contributions

J.A., J.W. and Z.W. led the research. J.A., P.K., J.W. and Z.W. developed the computational research strategy. F.H.A., J.W. and Z.W. developed the experimental research strategy. S.S. coordinated and managed the research project. H.P., G.M., and L.C. coordinated and managed the GDM wet lab. J.A., A.E.C., J.K., J.S., A.T., L.K.W., Y.W., and Z.W. developed the neural network architecture and training procedure. J.A., T.F., J.K., E.P., R.G.S., J.W., V.F.Z., A. Žemgulytė and A. Žídek developed the data, inference and evaluation setup. J.A., A.E.D., T.F., J.K., D.L., P.L., L.S., J.W., Z.W., R.K.Z. and Y.Z. created designs for experimental testing. A.E.D., M.D., T.O.C.K., Z.-Q.L., A.N.M., H.P., J.C.R., L.S., R.K.Z., N.A., K.S., and C.E. performed the wet lab experimentation. J.A., A.E.C., A.E.D., M.D., T.F., J.K., P.K., T.O.C.K., D.L., Z.-Q.L., L.S., S.S., A.T., J.W., L.K.W., Z.W. and R.K.Z. contributed to the writing of the paper. R.F. and D.H. advised on the project.

