## Supplementary Information for "Designing enzymes for new-to-nature chemistry and non-natural substrates with AlphaProtein Novo"

### 1 Supplemental Figures

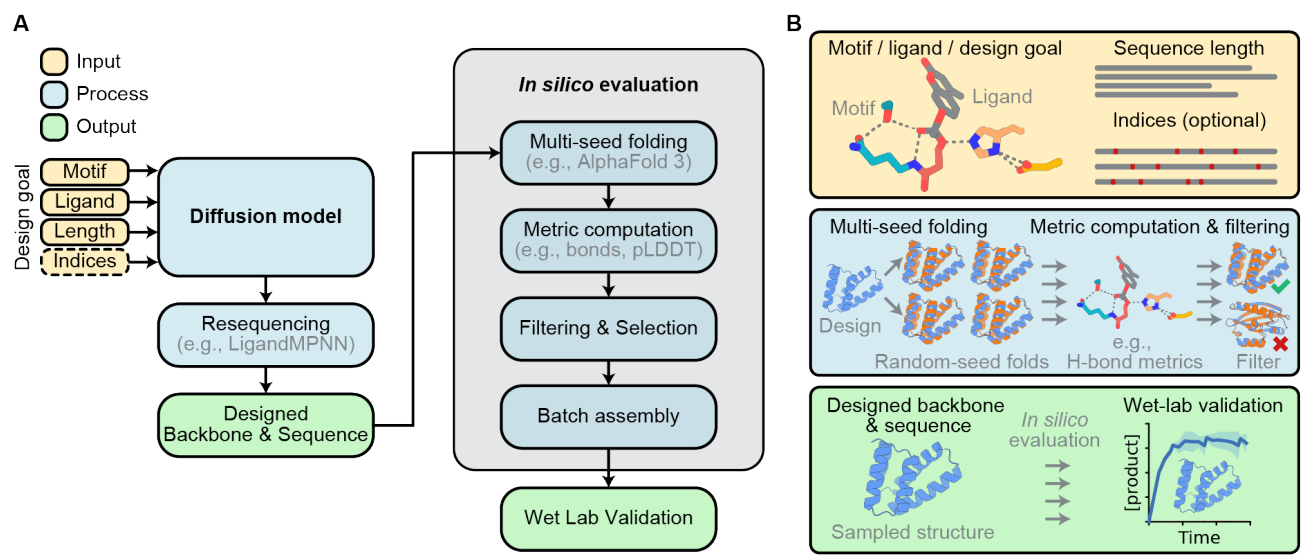

**Figure S1. End-to-End Pipeline.** A) High-level overview. A design goal (e.g., target reaction) is specified along with conditioning inputs (motif, ligand, length, and optional residue indices). The diffusion model generates candidate backbones and sequences via conditional diffusion, optionally with a subsequence resequencing (e.g. LigandMPNN) step. Candidates then undergo *in silico* multi-seed AlphaFold 3 structure prediction, metrics computation (e.g., hydrogen bonds, pLDDT), and filtering and selection. Passing designs are assembled into batches for wet-lab validation. B) Molecular-level illustrations of each stage, depicting motif/ligand inputs, multi-seed fold predictions, metric-based filtering, and the progression from sampled structures through *in silico* evaluation to experimental characterisation.

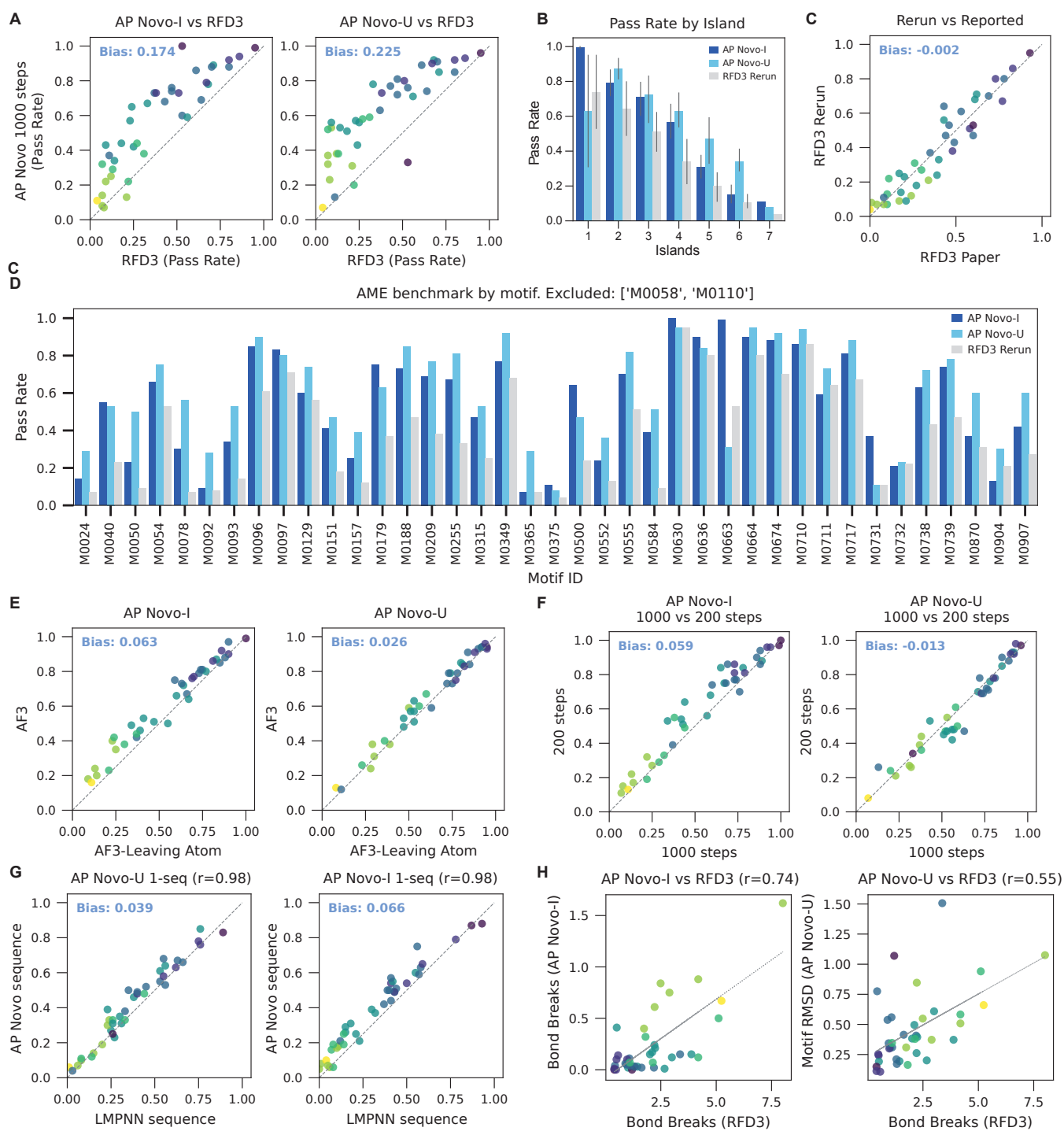

**Figure S2. AME Benchmark Comparisons.** A) Comparison by spec for APN indexed (APN-I) and APN unindexed (APN-U) against RFD3. B) Comparison by number of residue islands. C) Our simplified rerun of RFD3 correlates well with reported values in their paper. D) Alternate view of A, comparing pass rates by spec. E) Benchmark performance is similar when scoring with AF3 versus AF3-LA. F) APN performance with a lower number of sampling steps. G) The performance of APN sequences (No Reseq) is similar to LigandMPNN (LMPNN), as evaluated on single sequences. H) APN-I motif bond breaks and APN-U's reference-sample motif RMSD correlate with RFD3 bond breaks.

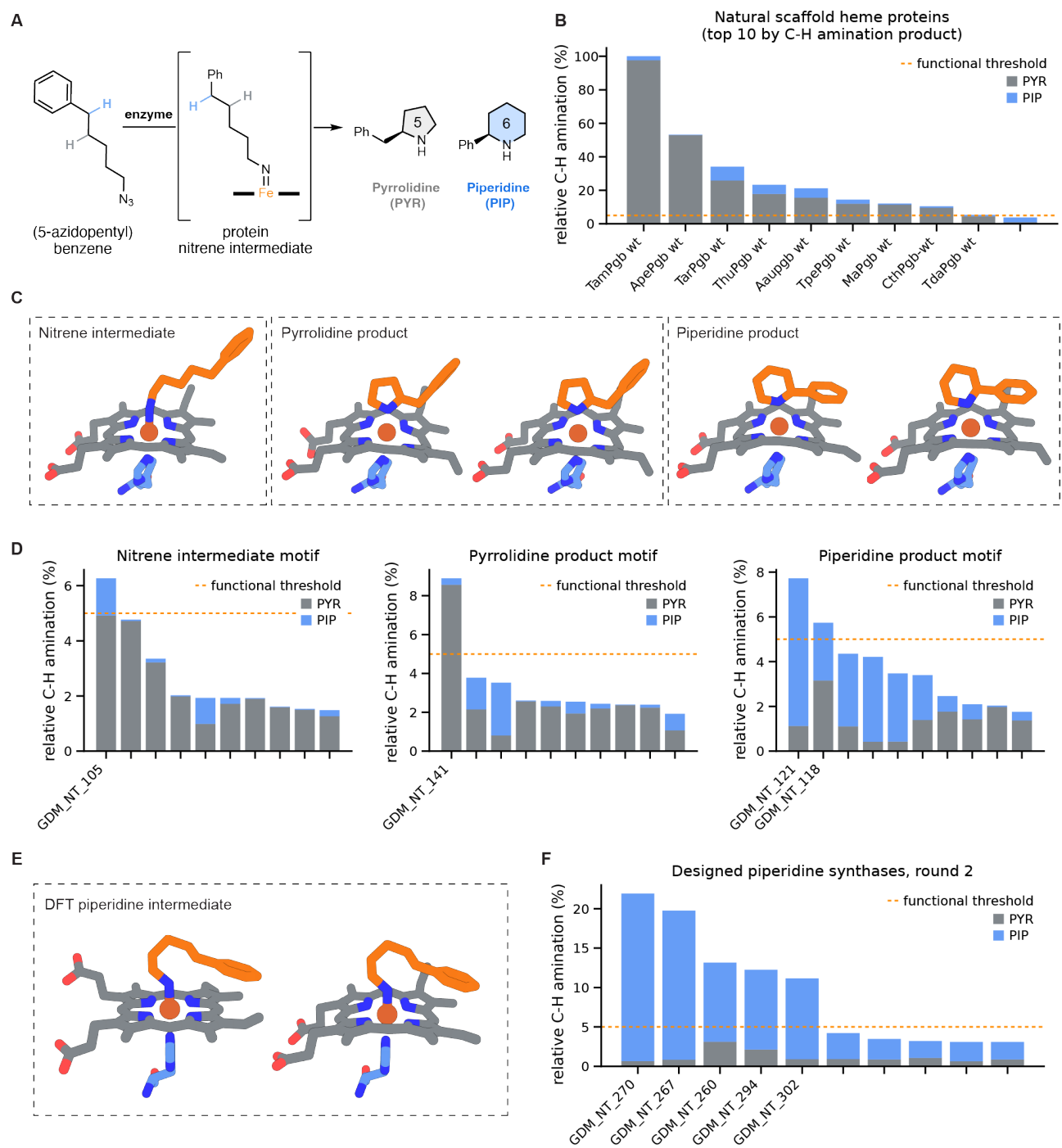

**Figure S3. Application of APN to design novel protein scaffolds for regioselective C–H amination.** A) Intramolecular C–H amination of (5-azidopentyl)benzene yields piperidine or pyrrolidine products. B) Screen of natural and previously engineered natural heme proteins on the reaction shown in A ($n=188$). Product formation and distribution by the top 10 variants are shown. C) Representative motifs for round 1 APN scaffold generation. D) Experimental activity of round 1 designs categorised by design motif; top 10 designs shown. E) Representative DFT-derived motifs for round 2 APN generation. Piperidine product motifs shown in (C) were also included in round 2 generation. F) Experimental performance of round 2 piperidine synthases targeted specifically for piperidine synthesis; top 10 designs shown.

Unless otherwise noted, whole-cell transformations were performed with 5 mM substrate at room temperature, as described in the experimental methods. Variants were screened without individual confirmation of protein expression. Functional threshold (dashed line) represents 5% of *TamPgb* wild type (wt) C–H amination activity. Activities are normalised to *TamPgb* wt within the same experiment; enzymes exceeding the threshold are labeled. Natural scaffold abbreviations: Tam, *Thermus amyloliquefaciens*; Ape, *Aeropyrum pernix*; Tar, *Thermus arciformis*; Thu, *Thermoflexus hugenholtzii*; Aau, *Acetithermum autotrophicum*; Tpe, *Thermorudis peleae*; Ma, *Methanosarcina acetivorans*; Cth, *Crenotalea thermophila*; Tda, *Thermanaerothrix daxensis*; Pgb, protoglobin

**A** Biocatalytic performance of natural and APN nitrene transfer enzymes

| Name | Description | [Enz]<sub>active</sub><br>(μM) | TTN<br>to<br>PIP | PIP:PYR | %e.e. |
| --- | --- | --- | --- | --- | --- |
| *TamPgb*<br>wt | Natural scaffold | 6.5 | 0.1 | 3:97 | n.m. |
| GDM_NT<br>121 | Novel scaffold,<br>round 1 | 2.6 | 1.7 | 94:6 | -68% |
| GDM_NT<br>270 | Novel scaffold,<br>round 2 | 1.2 | 22 | 99:1 | 94% |

**B**

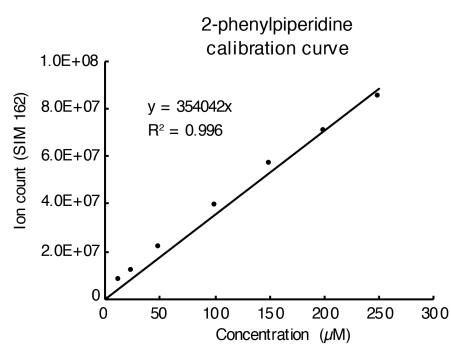

**C**

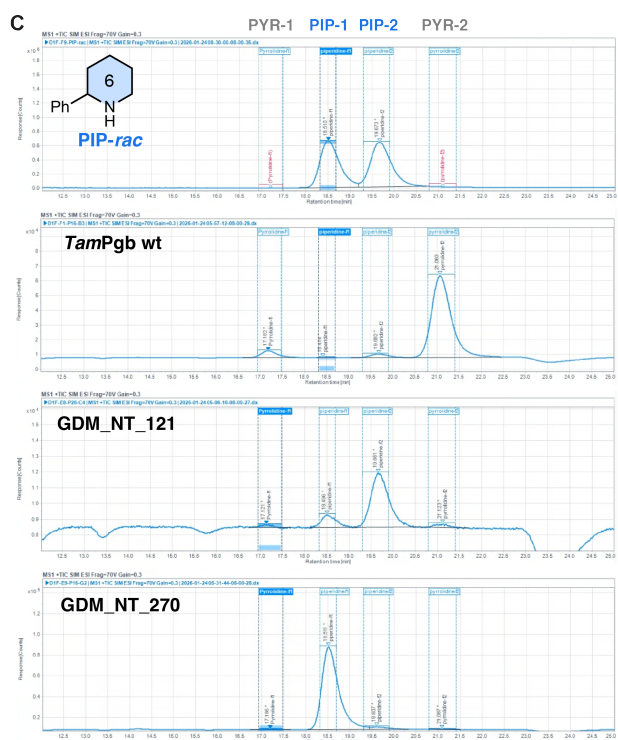

**Figure S4. Summary of top performing APN nitrene transferases in each design round compared to natural enzyme *TamPgb* wt.** A) Biocatalytic performance of APN nitrene transfer enzymes and *TamPgb* wt for the reaction shown in Figure S3A. Reactions were performed under standard conditions (5 mM substrate, room temperature) using OD<sub>600</sub> normalized clarified lysate. Reactions were performed in triplicate; standard deviation for TTN to PIP is within 20% of the mean, which is listed in the table. B) Calibration curve used to quantify 2-phenylpiperidine formation from biocatalytic reactions via reverse-phase HPLC-MS. Calibration is acquired on the same instrument and in the same sequence as the experimental samples. C) Representative chiral HPLC-MS for determination of enantiomeric excess. Abbreviations: PIP, 2-phenylpiperidine; PYR, pyrrolidine shown in Figure S3A; TTN, total turnover number; %e.e., enantiomeric excess; [Enz]<sub>active</sub>, active enzyme concentration in the reaction setup as determined by hemochrome assay; n.m., not measured; wt, wild type.

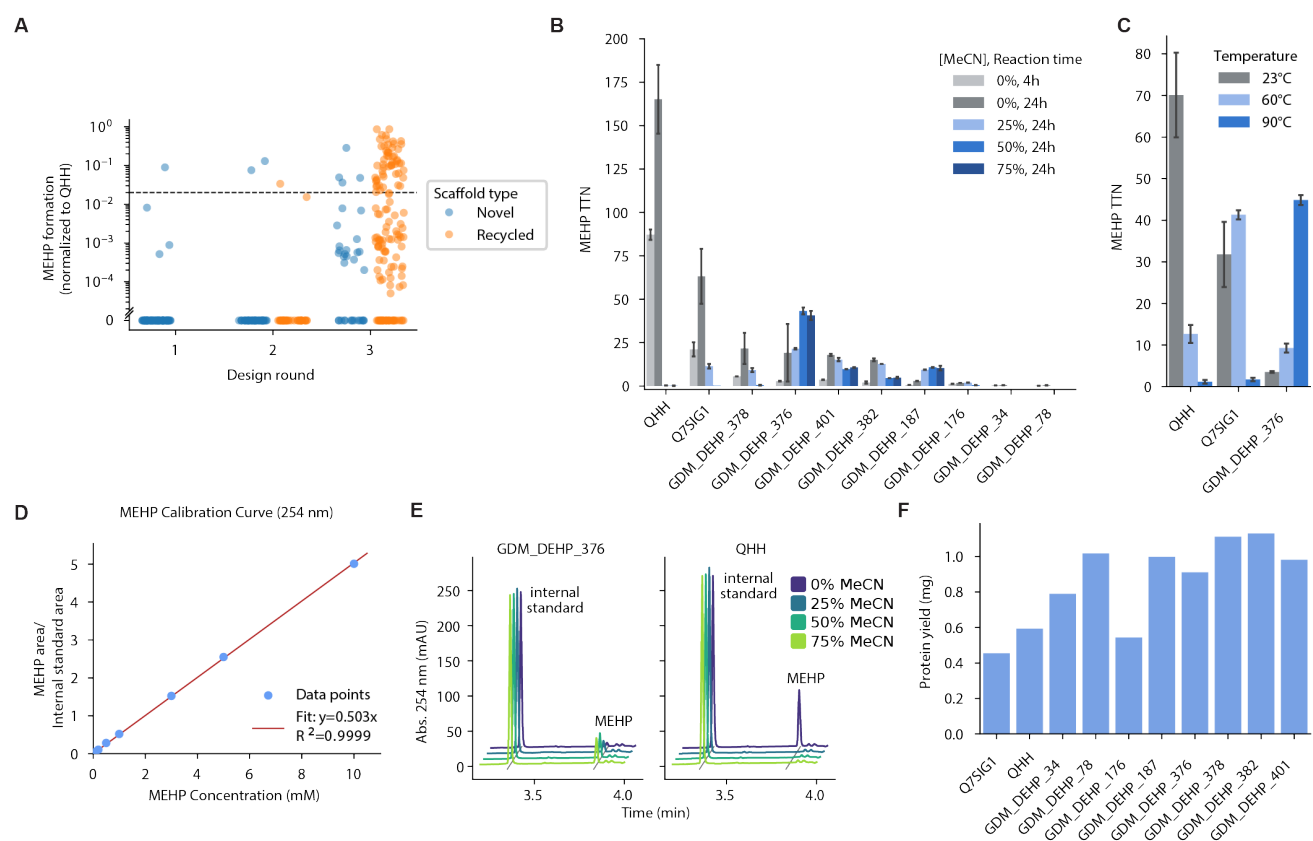

**Figure S5. DEHPase supporting characterisation** A) Lysate activity of DEHPase designs (amount of mono(2-ethylhexyl) phthalate (MEHP) formation normalised to natural esterase QHH) from screening assay, versus design round and scaffold type (novel- versus recycled-scaffold). Note that these measurements convolve the effect of protein expression and specific activity. Dotted line: threshold for calling hits (2% of QHH activity). B) Activity of purified DEHPases (total turnovers from DEHP to MEHP) at different acetonitrile (MeCN) concentrations for top hits from screening. Error bars represent standard deviation of 2-4 biological replicates. TTN, total turnover number measured at the indicated condition. C) Activity of purified DEHPases after 4-hour reactions at different temperatures. D) Calibration curve used to quantify MEHP. A calibration curve for phthalic acid (PA) was also generated; however PA was only observed in QHH samples and amounted to < 10% of MEHP formed. E) Overlaid representative HPLC chromatograms of DEHP hydrolysis catalyzed by GDM\_DEHP\_376 and QHH after 24 hours at room temperature. The traces highlight the internal standard and MEHP peaks under varying MeCN concentrations (0–75%) in the reaction buffer, illustrating that GDM\_DEHP\_376 exhibits higher activity in the presence of MeCN. F) Protein yield (mg) after expression in 6 mL cultures of *E. coli*.

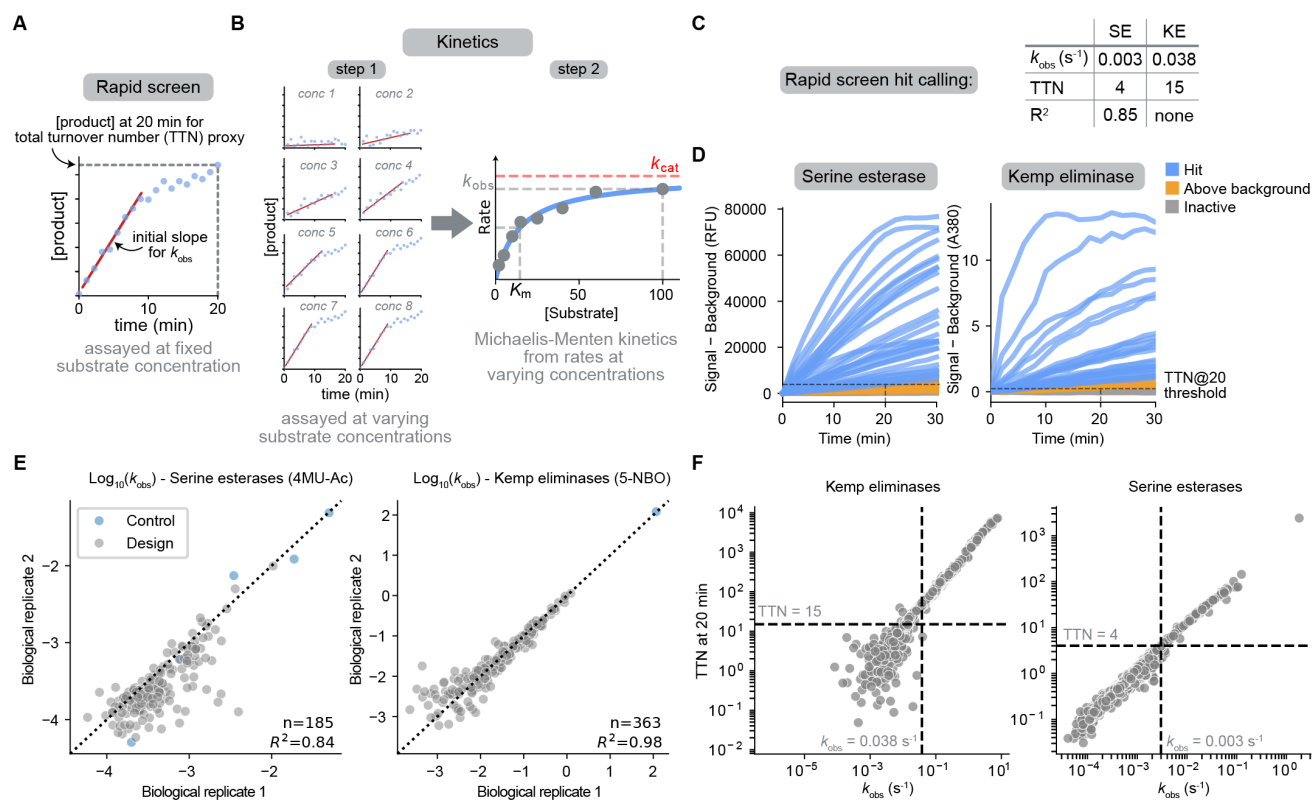

**Figure S6. Principles of $k_{obs}$ measurement, hit calling, and Michaelis-Menten kinetic measurements.** A) Schematic representation of enzyme screening assays to measure the initial rate $k_{obs}$. Assays are performed at fixed substrate concentrations in technical and biological replicates and used to determine initial rates and a total turnover number proxies at 20 minutes. B) Schematic representation of Michaelis-Menten kinetic data acquisition. Assays are performed at varying substrate concentrations to determine initial rates. Resulting data is used to fit Michaelis-Menten curves and determine kinetic parameters. C) Thresholds used for hit calling in rapid screens for SE (serine esterase) and KE (Kemp eliminase) model reactions. D) Raw traces for a design round of serine esterases (left) and Kemp eliminases (right). Progress curves for variants that show activity above background but do not satisfy the hit-calling requirements are highlighted in orange, progress curves of hits are blue, and those of inactive variants are grey. E) Reproducibility of $k_{obs}$ measurements across biological replicates. F) Correlation of $k_{obs}$ and total turnover number proxy at 20 minutes. Dashed lines indicate applied thresholds for hit identification.

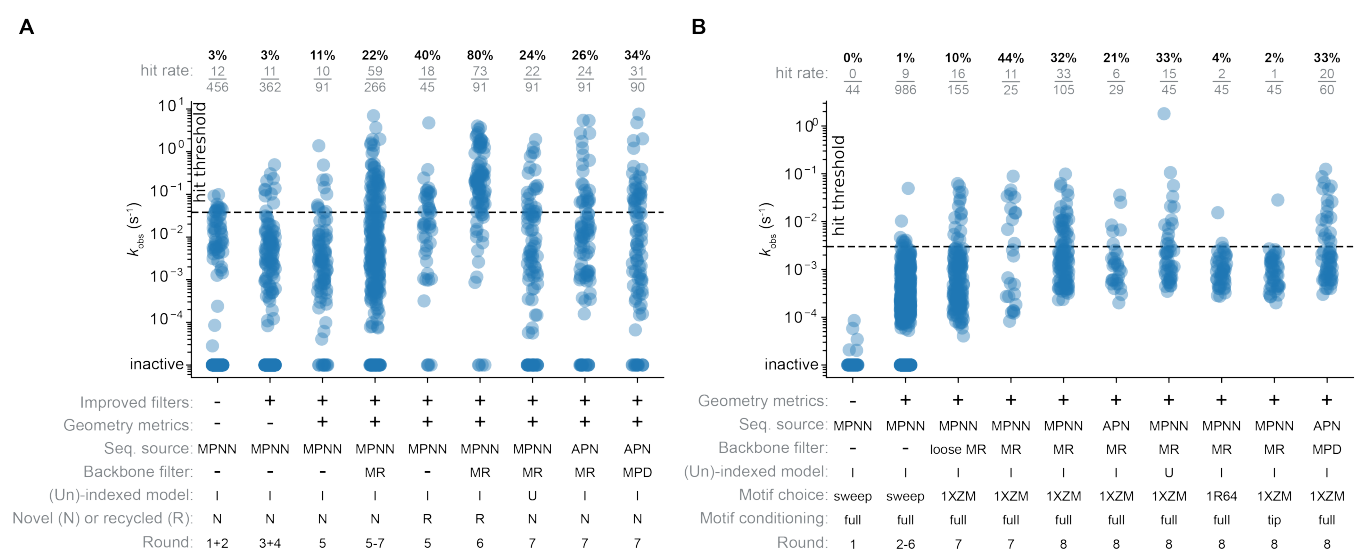

**Figure S7. Model reaction rates grouped by facet.** $k_{obs}$ values for (A) Kemp eliminases (B) and Serine esterases. Facets are grouped by design setup, which comprises whether geometry metrics were applied, the sequence source as LigandMPNN or APN, the backbone ensemble filtering strategy (multiple-resequencing (MR), multiple-partial-diffusion (MPD)), whether a motif with indexed residues (I) or an unindexed motif (U) was used, whether the designs were novel (N) or recycled (R), whether the full motif was used (full) or only tip-atoms were specified (tip), and which design rounds the facet is sourced from.

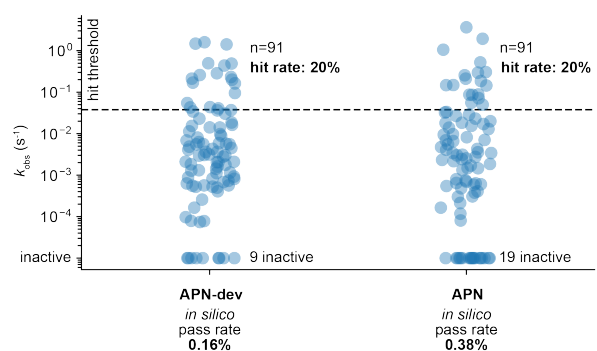

**Figure S8. *In silico* and wet-lab performance during the development of the APN diffusion model.** Comparison of $k_{obs}$ distributions, hit rates, and *in silico* pass rates between the APN diffusion model and an earlier development version of the model (APN-dev) for Kemp eliminase design. Although the APN model has more than 2x higher *in silico* pass rates than APN-dev, when the designs are selected using the same filters (Table S7, S8), their measured activity distributions are similar. All designs are resequenced with LigandMPNN before computing *in silico* pass rates or selection for experimental testing.

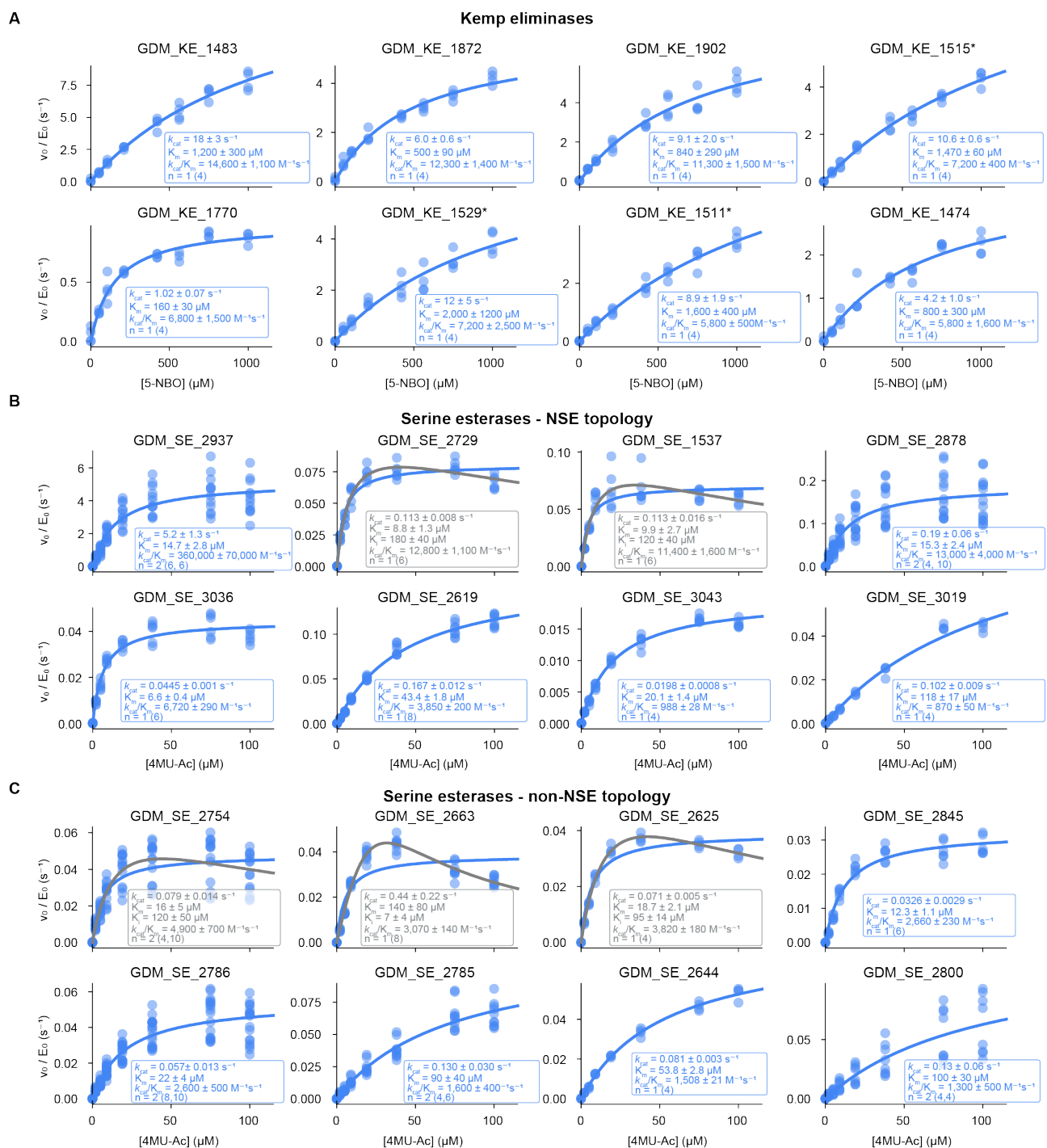

**Figure S9. Michaelis-Menten kinetics for selected Kemp eliminases and serine esterases.** A) Michaelis-Menten kinetics for Kemp eliminases (\*denotes recycled scaffolds). B) and C) Michaelis-Menten kinetics for serine esterases, subdivided into NSE topology-variants (B) and non-NSE topology variants (C). Biological replicates are listed as $n=x$, with the technical replicates per biological replicate listed in parentheses. Errors represent standard deviation

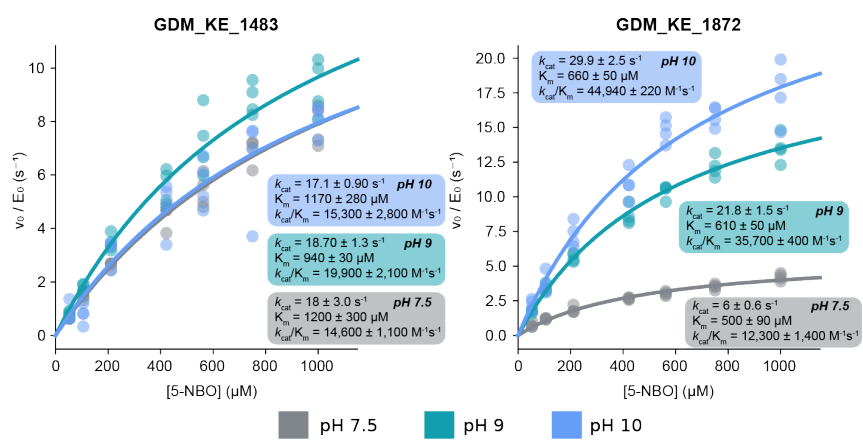

**Figure S10. Characterisation of designed Kemp eliminases at varying pH values.** GDM\_KE\_1483 and GDM\_KE\_1872 were characterised at pH 7.5, 9, and 10. Inferred values are listed in the figure (shaded backgrounds). Errors represent standard deviation of four technical replicates.

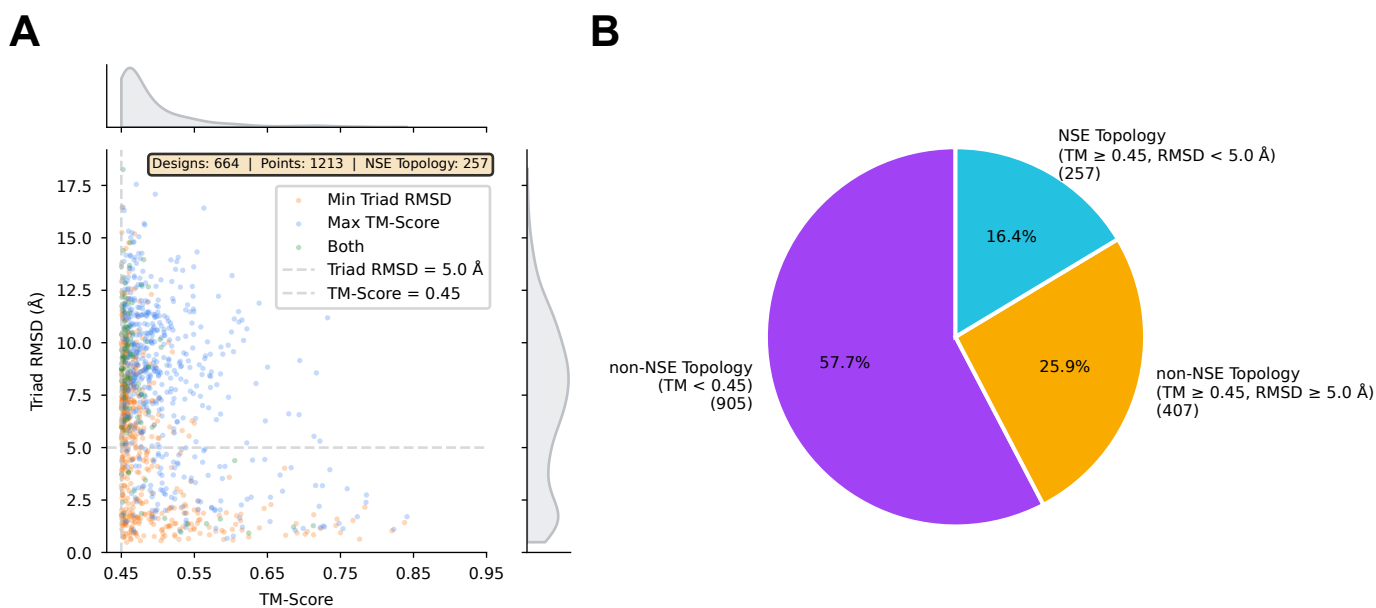

**Figure S11. Structural novelty of designed serine esterases.** A) Scatter plot of TM-score versus triad RMSD for each design. Triad RMSD was computed only for PDB hits with TM-score > 0.45; for each design, the search was terminated once a hit with triad RMSD < 2.0 Å was found. Thus, points below the 2.0 Å threshold represent upper bounds on the triad RMSD. Each design contributes up to two points: the hit with the highest TM-score (blue), the hit with the lowest triad RMSD (orange), or a single point when both criteria select the same hit (green). Dashed lines indicate classification thresholds (TM-score = 0.45, triad RMSD = 5.0 Å) for defining natural serine esterase (NSE) topology. B) Proportion of designs classified by structural similarity. Designs meeting both thresholds (TM-score ≥ 0.45 and triad RMSD < 5.0 Å; 16.4%) are classified as adopting a NSE topology; all remaining designs are classified as non-NSE topology, comprising those with a similar fold but divergent active-site geometry (TM-score ≥ 0.45 and triad RMSD ≥ 5.0 Å; 25.9%) and those with no close structural match (TM-score < 0.45; 57.7%). Counts shown in parentheses.

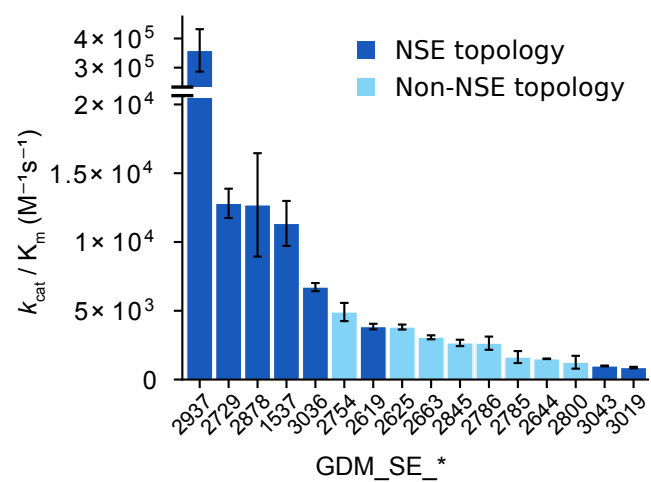

**Figure S12. Catalytic efficiency ($k_{cat}/K_m$) of designed serine esterases, sorted in descending order.** Designs adopt either a natural serine esterase (NSE) topology (dark blue) or a non-NSE topology (light blue). Error bars denote the standard deviation of the $k_{cat}/K_m$ ratio from independently fitted $k_{cat}$ and $K_m$ values across at least four technical replicates.

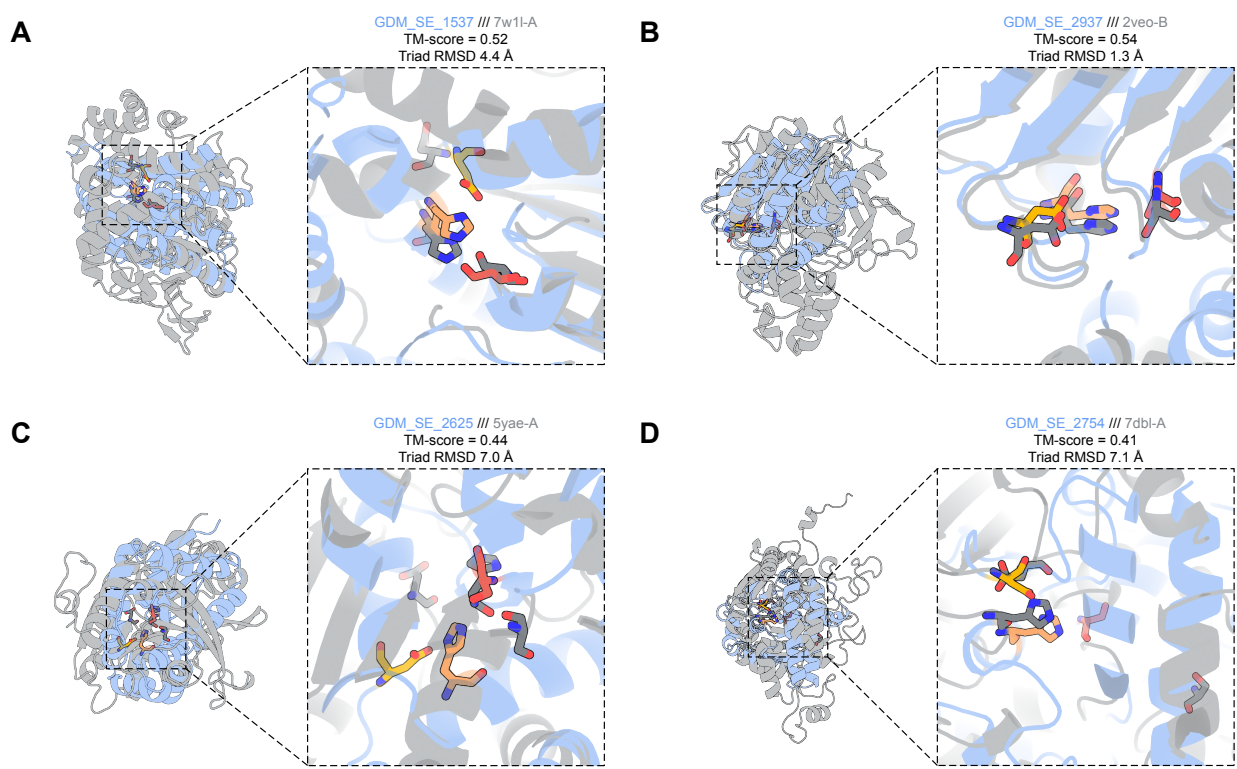

**Figure S13. Structural comparison of designed serine esterases with their closest EC-annotated PDB hits.** Each panel shows a design (blue) superimposed on its highest TM-scoring EC-annotated PDB chain (grey). PDB hits were identified by running TM-align against all PDB chains annotated with EC 3.1.1 or EC 3.4.21. For each hit, the corresponding catalytic triad residues in the PDB chain were identified by geometric motif matching, and the triad RMSD was computed as the all-atom RMSD over the backbone and side-chain heavy atoms of the three matched triad residues. Hits were filtered for TM-score > 0.45 and triad RMSD < 5.0 Å, and the hit with the highest TM-score was selected. Designs were structurally aligned to their best-matching PDB chain using TM-align, and the TM-score (query-normalised) is reported for each pair. The catalytic triad residues (Ser, His, Asp) are shown as sticks and highlighted in both the design and PDB hit. Insets show a zoomed-in view of the active site. (A, B) Designs with natural serine esterase (NSE) topology. (C, D) Designs with non-NSE topology.

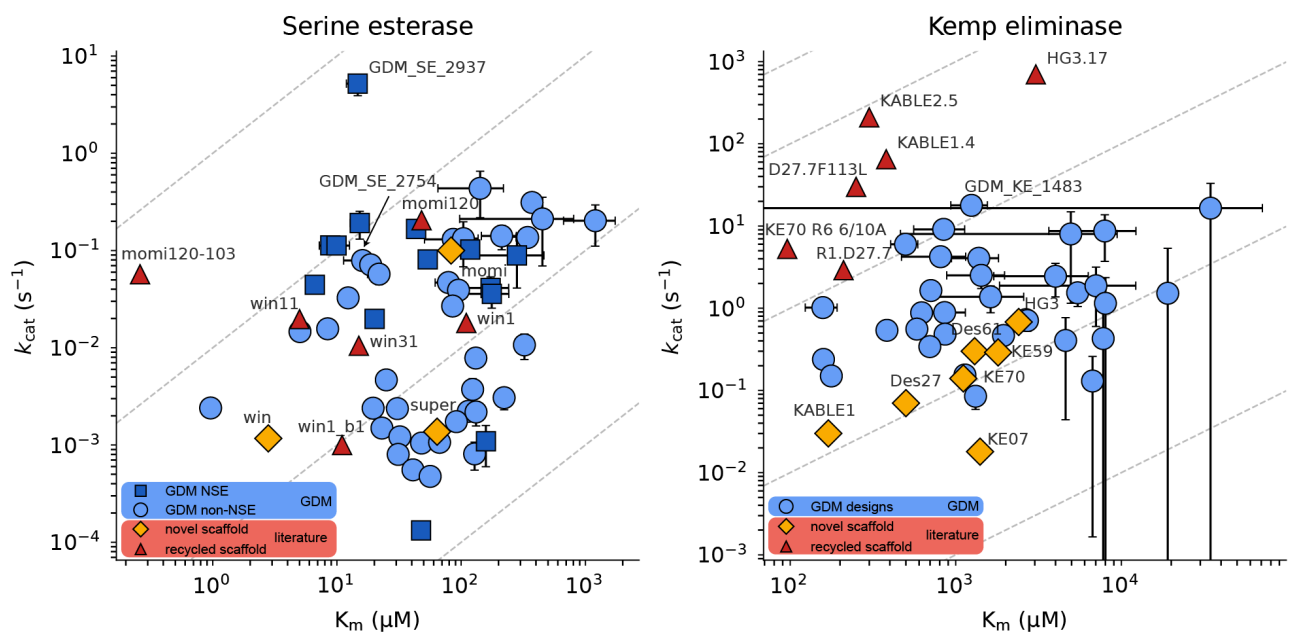

**Figure S14. Scatter-plot of catalytic rate and Michaelis constant for literature and in-house designs.** Catalytic rate ($k_{\text{cat}}$, $\text{s}^{-1}$) versus Michaelis constant ($K_{\text{m}}$, $\mu\text{M}$) for serine esterase (left) and Kemp eliminase (right) designs on log-log axes. Diagonal dashed lines indicate iso-efficiency contours of constant $k_{\text{cat}}/K_{\text{m}}$; catalytic efficiency increases towards the upper left. All GDM designs plotted (blue) are novel-scaffold and not derived from iterative optimisation of individual designs; for serine esterases, designs are further split into those with a natural serine esterase (NSE) topology or non-NSE topology. Literature variants are categorised as novel scaffold or evolved/mutated, where the latter encompasses variants derived from directed evolution, mutagenesis, partial diffusion, and other iterative optimisation methodologies. Error bars denote standard deviations across at least two technical replicates.

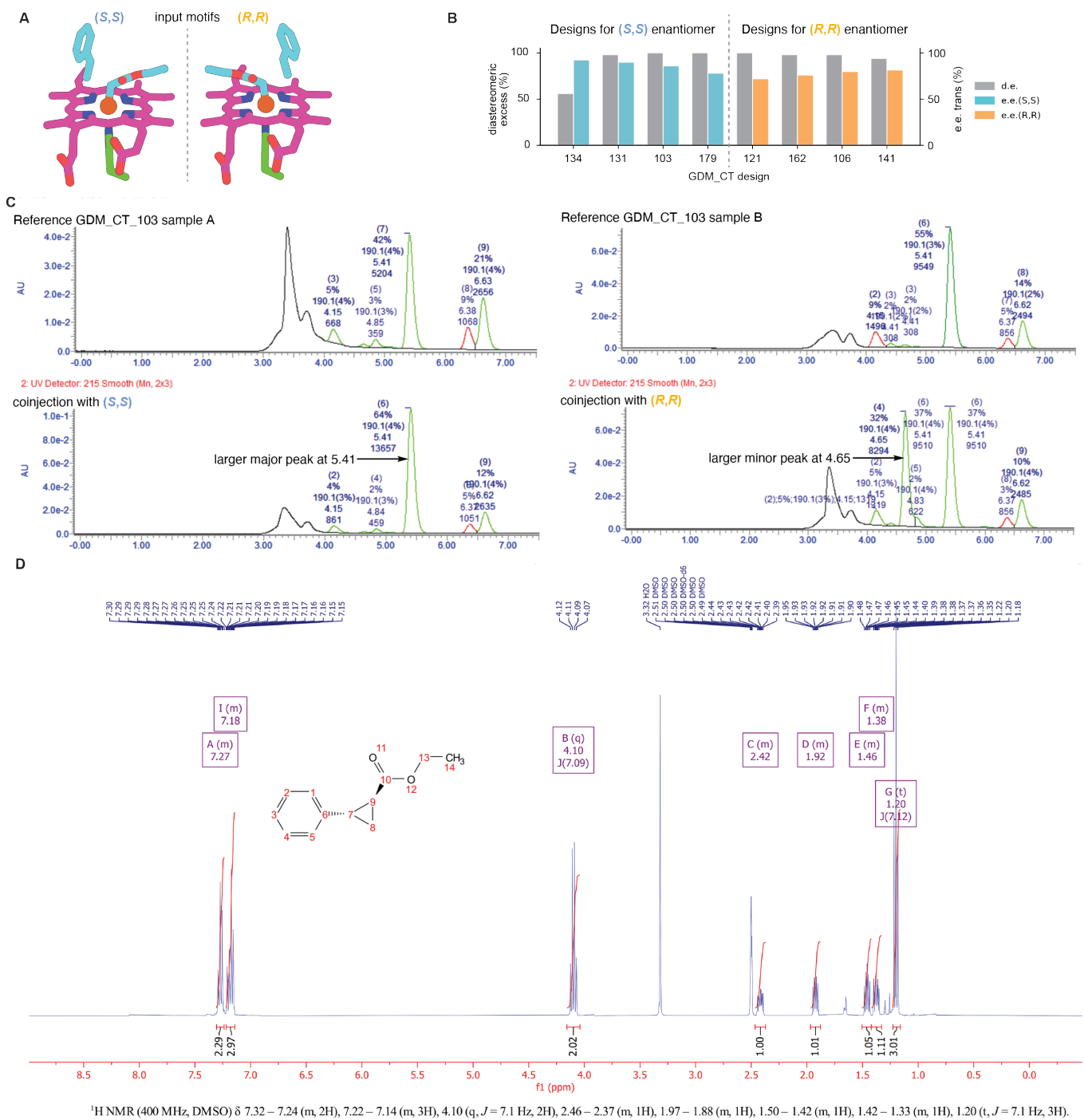

**Figure S15. Application of APN to novel-scaffold design for styrene cyclopropanation.** A) Example input motifs used for inference, as obtained from [1]. We searched for inverse rotamers of histidines that ligate heme through the PDB to sample backbone coordinates for the residue (not shown). B) Stereoselectivity screening results for the most enantioselective designs for each *trans* product. C) Co-injection experiment to validate the identity of the products against authentic product standards. The expected retention times are 4.65 min for the *R,R* and 5.41 min for the *S,S* product. For GDM\_CT\_103, which was designed for *S,S*, co-injection validates the expected product. D) The identity of the enzymatic product is further validated via NMR.

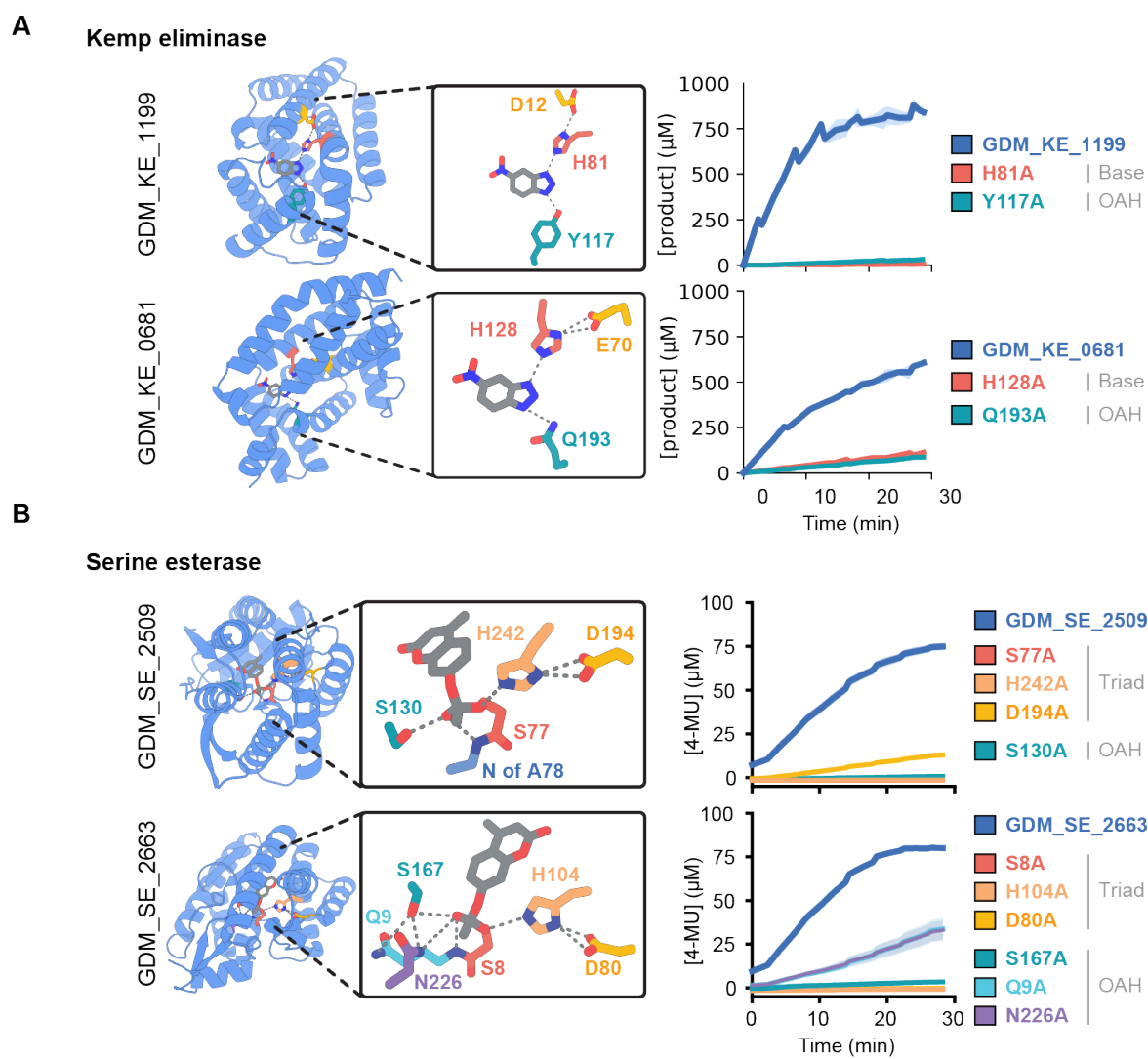

**Figure S16. Structural overview and experimental characterisation of additional knock-out mutants for A) Kemp eliminase and B) serine esterase.**

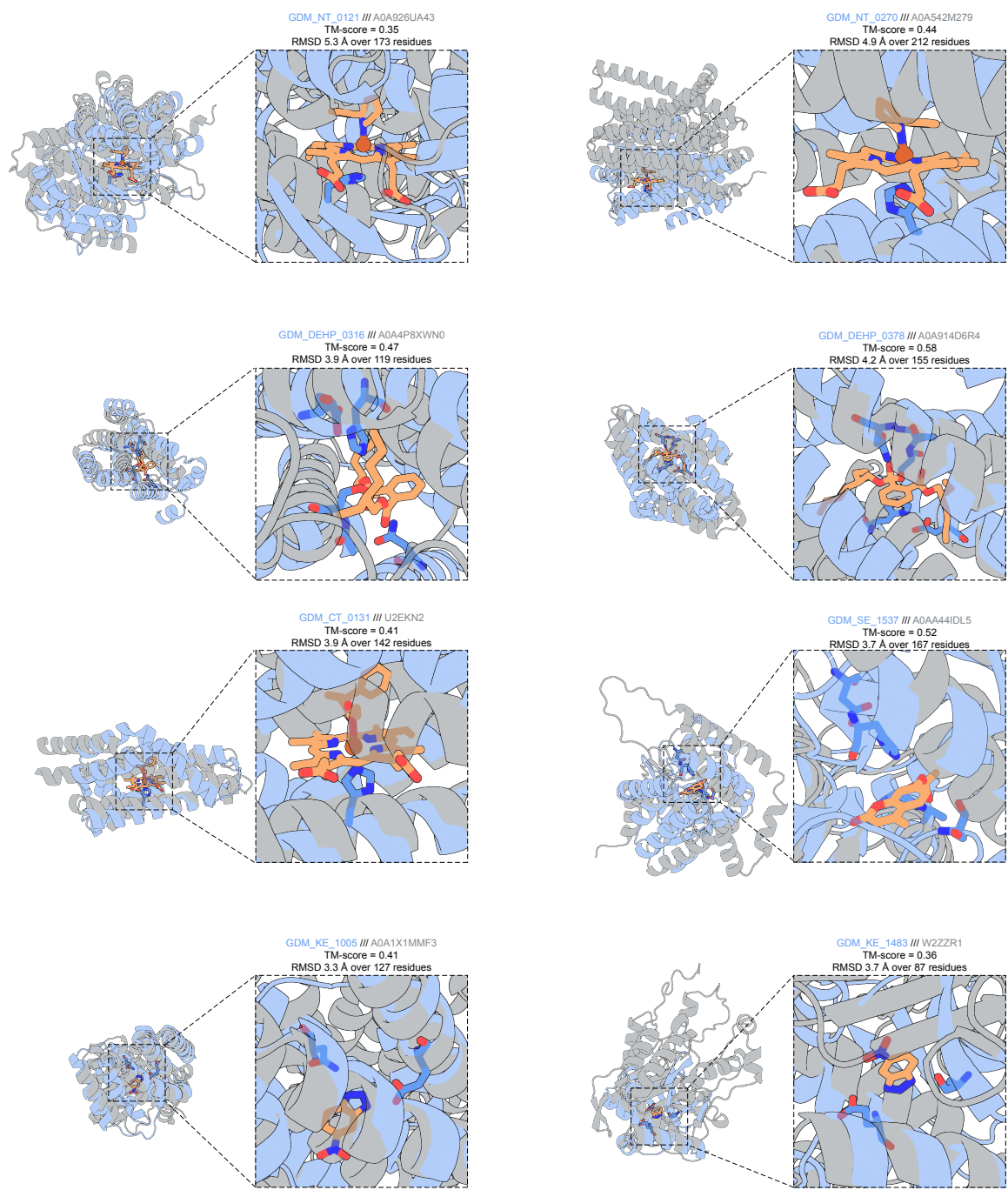

**Figure S17. Structural novelty of designs.** For each design, we use the predicted structure (using AF3 or AF3-LA) in complex with the ligand used to generate the design. We use Foldseek to find the closest structural match for each design to any structure in AFDB50v6 ([2]). We use the TM-align alignment mode for Foldseek, and otherwise the default parameters. For each design, we take the Foldseek hit with the highest hit probability. We then use TM-align to compute an exact TM-score (by default, Foldseek computes a fast approximation of the TM-score). We visualise the structures (aligned using TM-align) of each design and its top Foldseek hit in AFDBv6.

**Kemp eliminase**

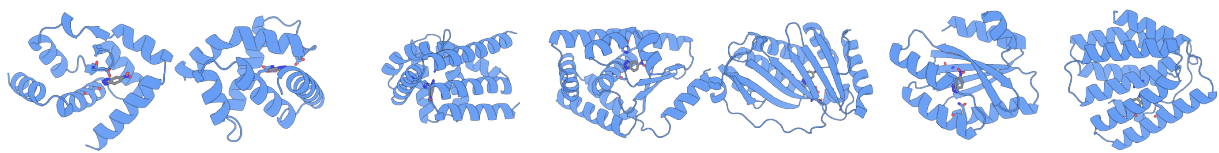

**Carbene transferase**

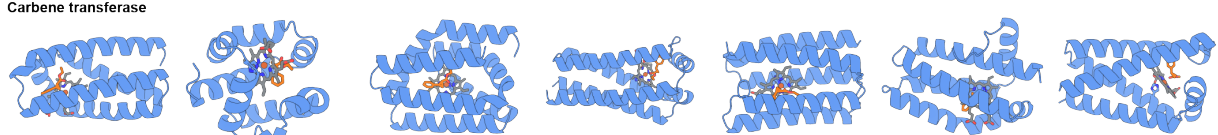

**DEHPase**

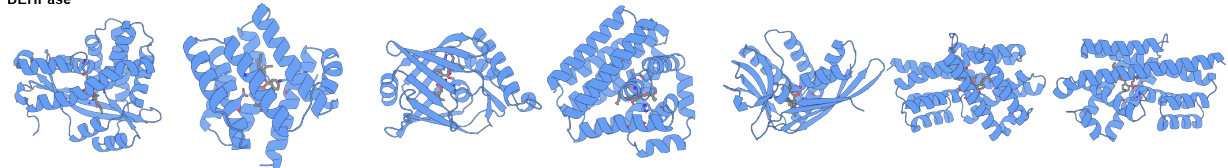

**Serine esterase**

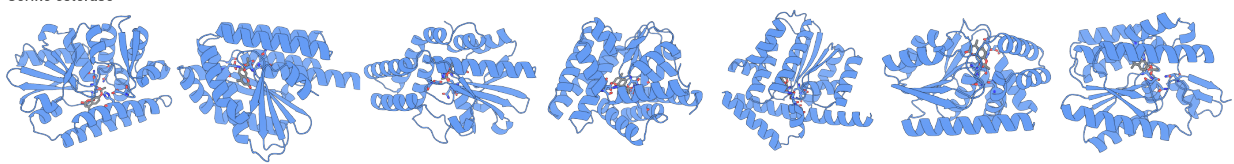

**Nitrene transferase**

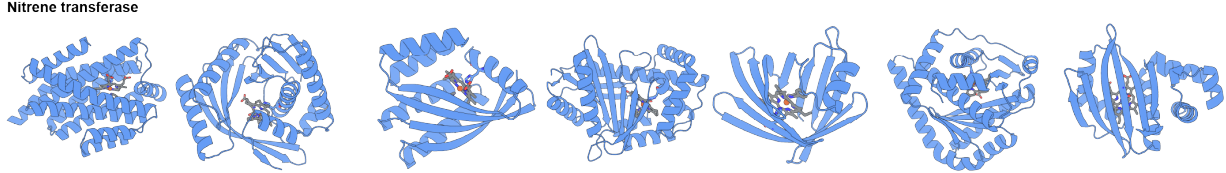

**Figure S18. Structural gallery of protein designs.** Representative structures across five target classes: Kemp eliminases, carbene transferases, DEHPases, serine esterases, and nitrene transferases. For each class, seven representative designs are shown, folded using AF3 or AF3-LA in complex with their respective ligands.

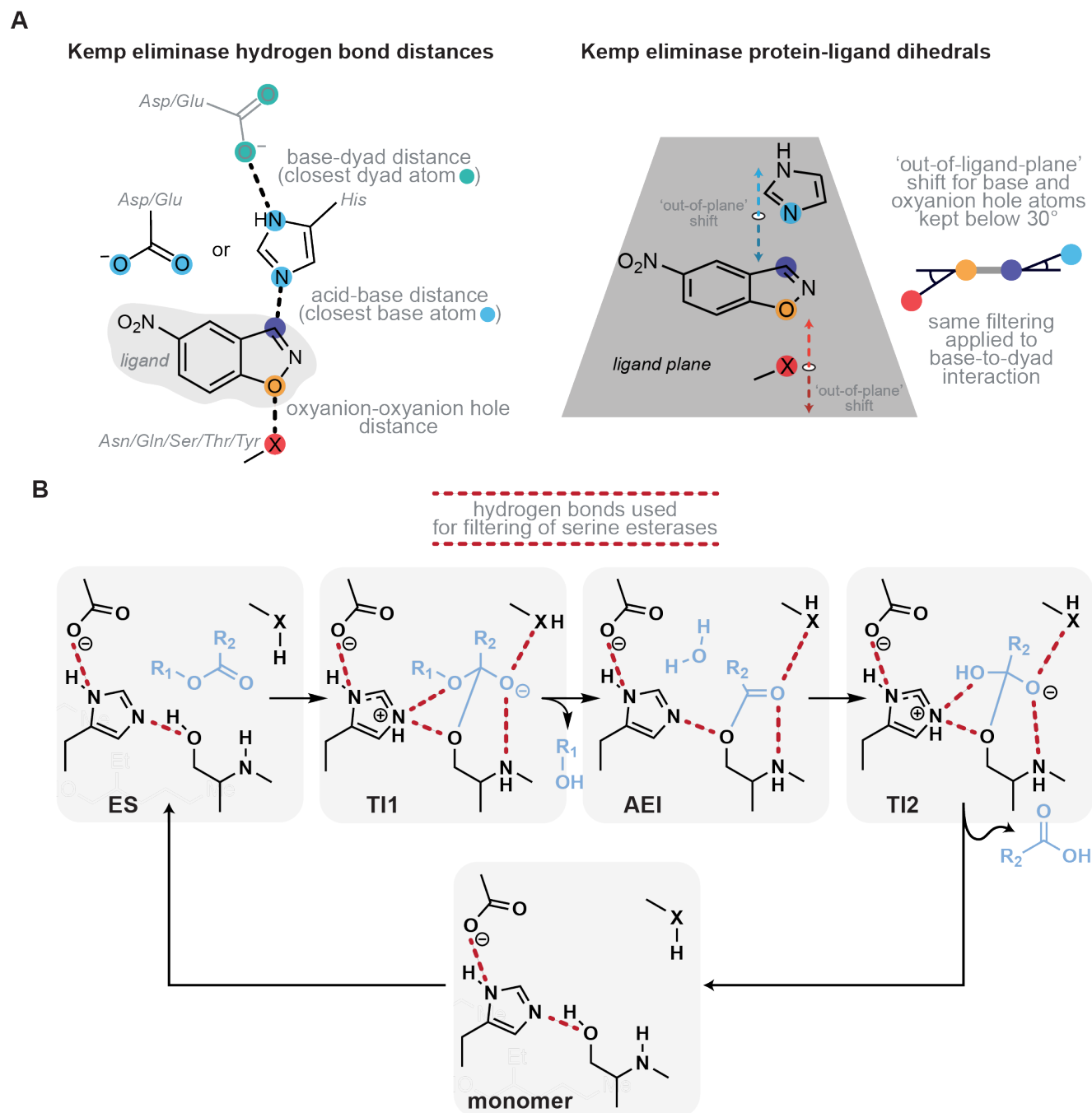

**Figure S19. Mechanistic filtering metrics.** A) Distance and geometry metrics that were used to filter Kemp eliminase designs. B) The catalytic cycle of serine esterases. 'Hydrogen-bonding' interactions that were used for filtering highlighted in dashed red lines.

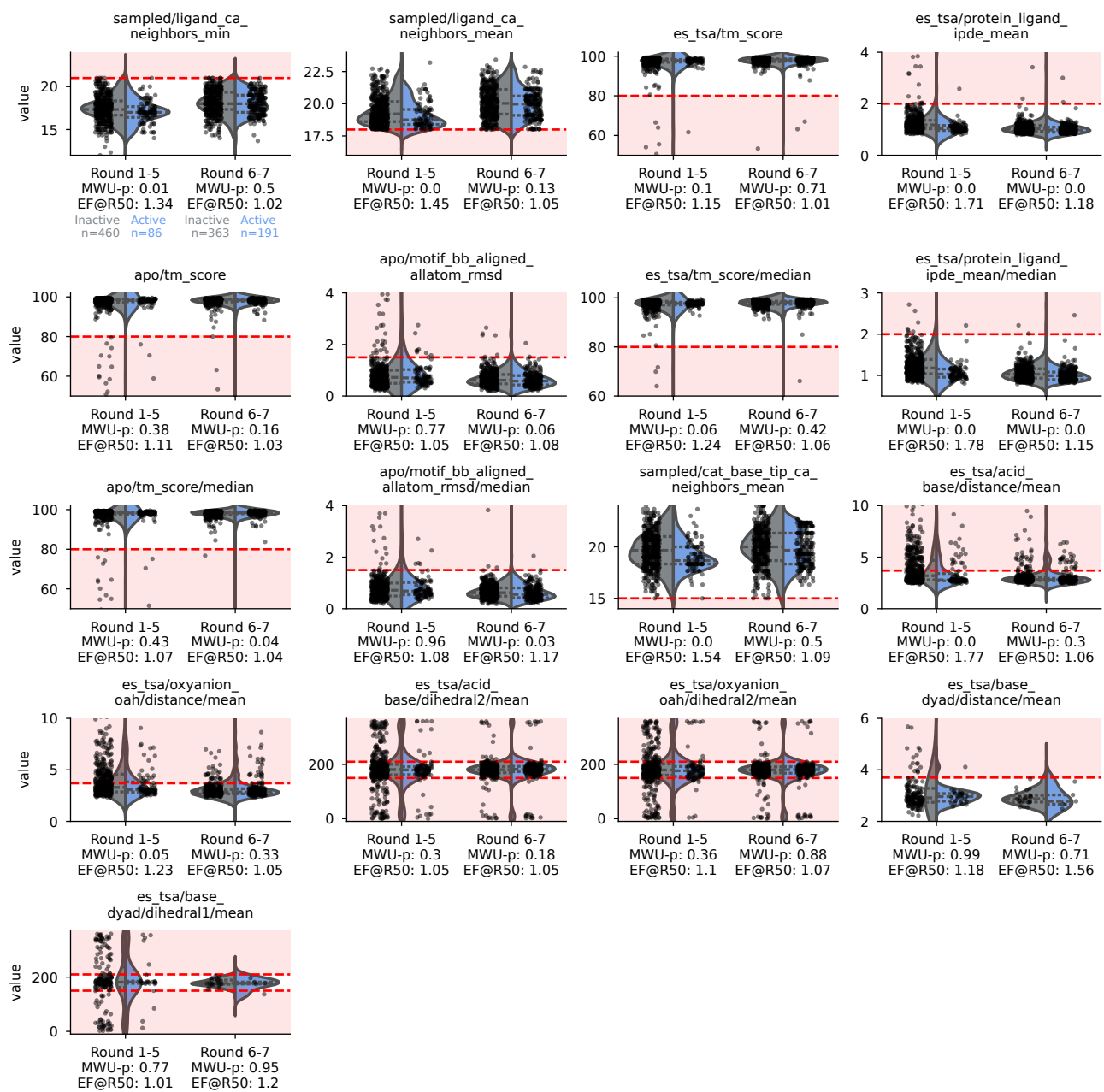

**Figure S20. Base filters for Kemp eliminase design.** Distributions of structure metrics, by hit status, for round 1-5 (some or no MR filtering) and round 6-7 designs (mostly MR filtered). Hits are indicated in blue on the right, and non-hits in grey on the left. Red line: threshold used for filtering (see Supplementary Methods). Some designs appear to fail the filters because thresholds were adjusted over time and structure predictions were recomputed for this analysis. "MWU-p": two-sided Mann-Whitney U test p-value. "EF@R50": enrichment over baseline precision at 50% recall.

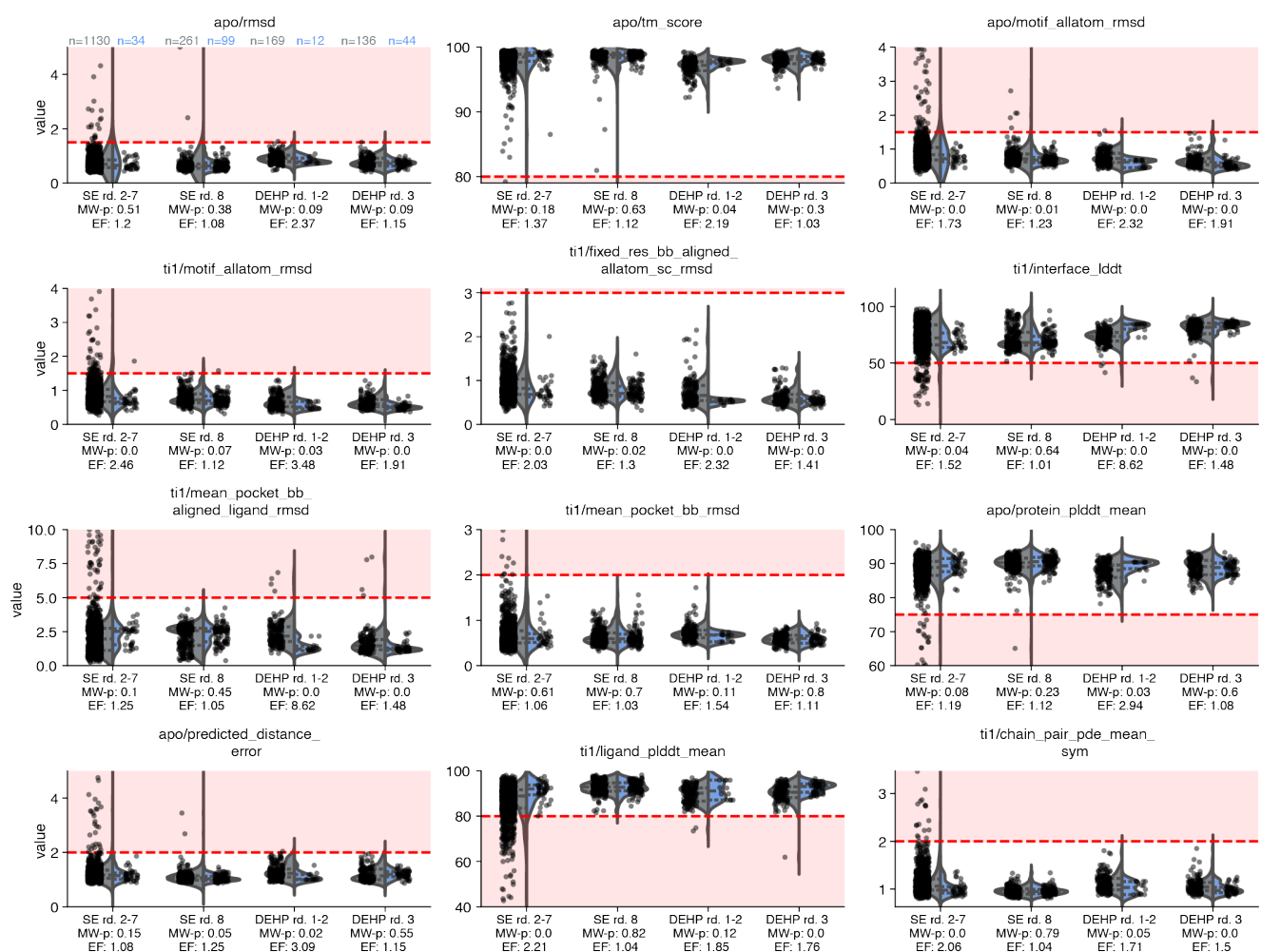

**Figure S21. Base filters for 4MU-Ac serine esterase and DEHPase design.** Distributions of structure metrics used for filtering designed serine esterases (for 4MU-Ac and DEHP). Hits are indicated in blue on the right, and non-hits in grey on the left. Red line: threshold used for filtering (see Supplementary Methods). The last design round is shown separately as the discriminating signal decreased in this round, likely due to correlations between these metrics and multi-resequence filtering. Some designs appear to fail the filters because thresholds were adjusted over time and structure predictions were recomputed for this analysis. "MW-p": two-sided Mann-Whitney U test p-value. "EF": enrichment over baseline precision at 50% recall.

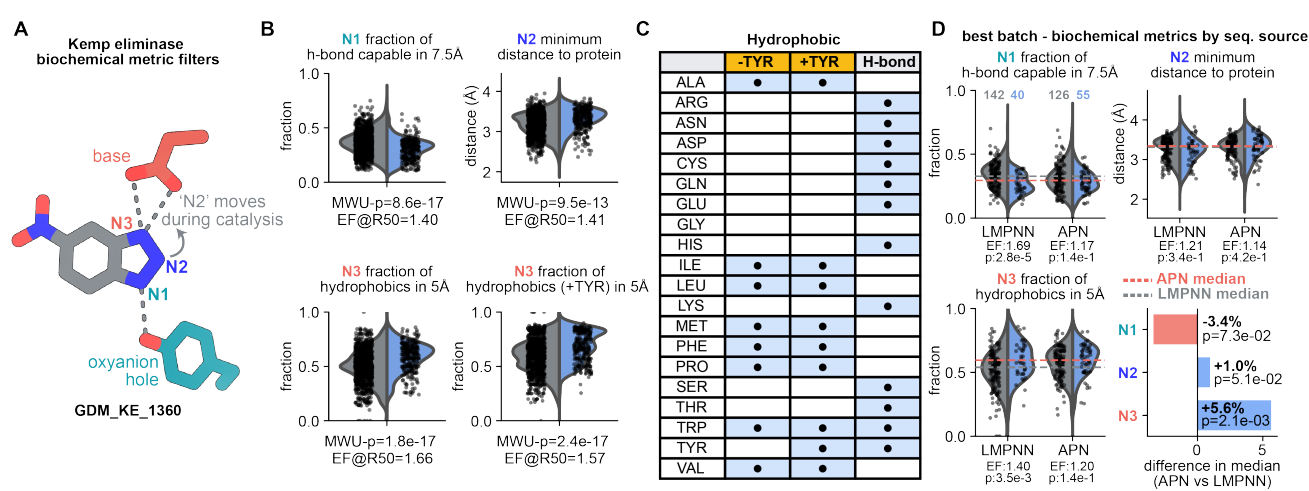

**Figure S22. Biochemical metrics help discriminate hits in Kemp eliminases.** A) Schematic of a Kemp eliminase active site (here: GDM\_KE\_1360) with individual nitrogen atoms of the inhibitor labeled as N1, N2, and N3. B) Biochemical metrics describing the chemical environment of positions that are involved in catalysis can be used to enrich hit rates. Hits are indicated in blue on the right, and non-hits in grey on the left. "MWU-p": two-sided Mann-Whitney U test p-value. "EF@R50": enrichment over baseline precision at 50% recall. C) Amino acids that were considered hydrophobic or H-bond capable for the analyses presented in this figure. D) Distribution of biochemical metrics by sequence source (APN or LMPNN) for the latest design round. The median of all plotted data is shown in dashed red lines for APN-derived sequences and in grey for LMPNN-derived sequences. To aid in highlighting the shift, both medians are highlighted for each respective violin plot. The bottom right panel highlights the absolute change (for N1 and N3) or relative change (for N2) in median between APN and LMPNN as the sequence source. EF: enrichment over baseline precision at 50% recall.

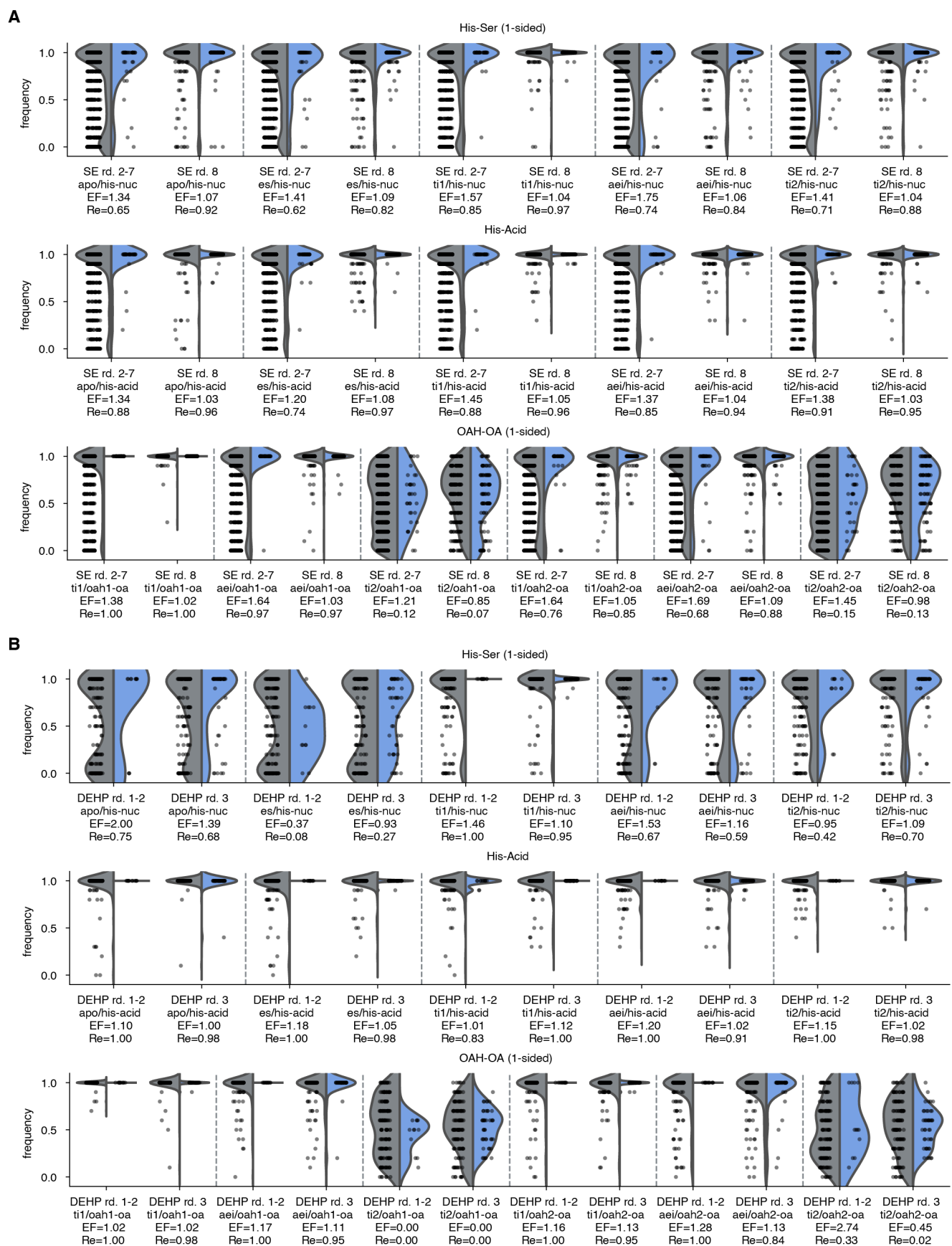

**Figure S23. H-bond filters for 4MU-Ac serine esterase and DEHPase design.** Distributions of H-bond metrics (frequency of H-bond in 10 prediction seeds) for (A) 4MU-Ac serine esterase and (B) DEHPase design. Hits are indicated in blue on the right, and non-hits in grey on the left. All H-bond metrics shown here were used for filtering except TI2 oxyanion hole (see Supplemental methods). Threshold for all H-bond metrics is $>0$ (at least 1 in 10 seeds). "EF": enrichment factor when requiring frequency=1.0; "Re": Recall at frequency=1.0.

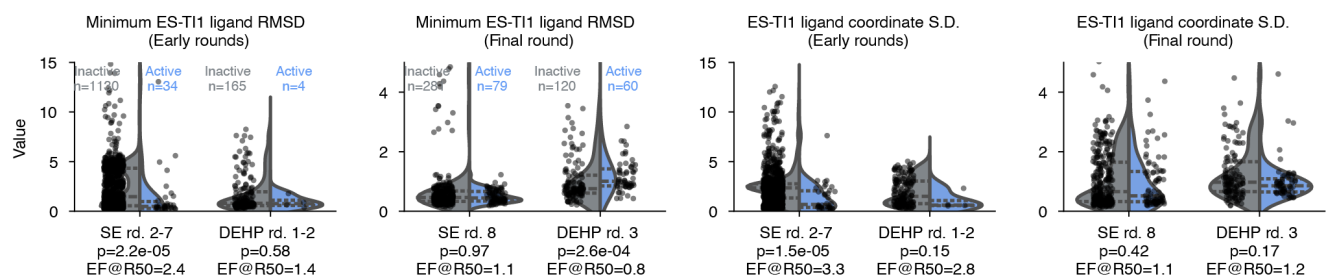

**Figure S24. ES-TI1 consistency filters for 4MU-Ac serine esterase and DEHPase design.** Distributions of metrics measuring ligand pose similarity between ES and TI1 states (see metrics definitions in Supplementary Methods). Hits are indicated in blue on the right, and non-hits in grey on the left. The final round of design is shown separately because it was filtered on these metrics. P-values from two-sided Mann-Whitney U test for differences in hit and non-hit metric distributions are indicated. "EF@R50": enrichment factor at 50% recall.

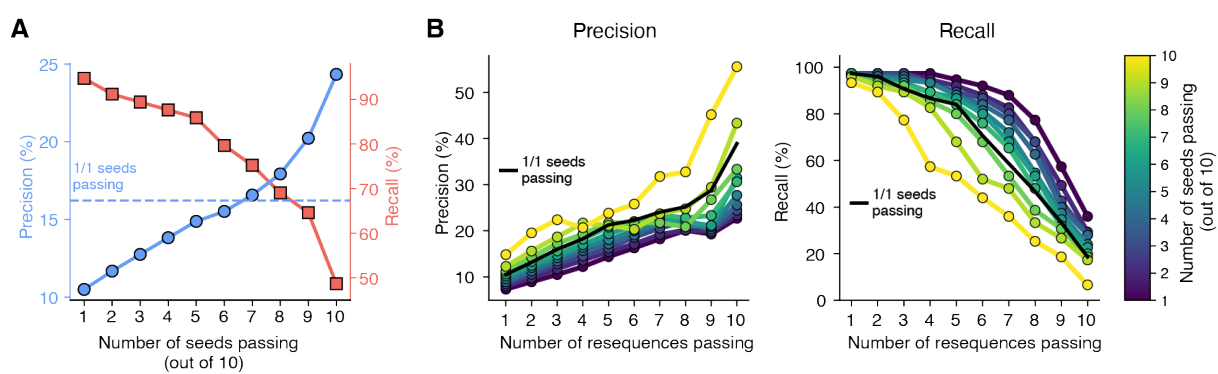

**Figure S25. Retrospective precision of multi-seed and multi-resequence filtering on serine esterase designs.** A) Precision (blue circles) and recall (red squares) versus number of structure prediction seeds passing H-bond filters over all serine esterase data ($n=1,524$). The dashed line shows the precision of filtering using only a single prediction seed. Designs have previously been filtered on base filters and 1/10 multi-seed H-bond filters. B) Precision and recall versus number of resequences and seeds passing, for designs which had not been filtered at 10/10 LMPNN-MR ($n=1,344$). Each design was resequenced 10 times and each resequence was predicted with 10 seeds. Black line shows the effect of only using one prediction seed for filtering each resequence.

**Figure S26. Backbone-conditioned sequence ensemble activity.** Activity of 9-11 LigandMPNN sequences with a common backbone not passing (blue) and passing (orange) the multiple-resequencing (MR) filter. Sequences for which the backbone had passed an MR filter have a higher probability of yielding more active sequences.

**Figure S27. Comparison of sequence characteristics and experimental performance between AP Novo and LigandMPNN designs.** A) Amino acid frequencies of LigandMPNN and AP Novo designed sequences. Frequencies are shown for both the full protein ("global") and for residues within 4 Å of a ligand ("pocket"). Sequences are taken from Kemp eliminase and serine esterase designs in the latest batches which have undergone filtering at or near our most effective procedures. For LigandMPNN, 357 Kemp eliminases and 240 serine esterases are shown; for AP Novo, 181 Kemp eliminases and 120 serine esterases are shown. B) $\log_{10} k_{obs}$ of LigandMPNN ("LMPNN") sequences versus AP Novo sequences from the same backbones. Dotted line: hit threshold. Data are 68 (34 for each sequence model) Kemp eliminases and 60 serine esterases (30 per model). All designs shown have been MR-filtered (see also Figure S28B) C) Deviation of partial diffusion designs from parent design. Sequence identity (left) and RMSD (right) to the parent design, for serine esterase re-designs generated for multi-partial-diffusion filtering.

**Figure S28. Using AP Novo to score and filter designed sequences.** A) Hit rate and $k_{obs}$ distributions for designs whose AP Novo sequence pass or fail filters (the tested sequence for the design comes from LigandMPNN). "Retrospective": For Kemp eliminase, round 4-6 designs pre-filtered on geometry metrics to isolate the effect of AP Novo scoring. For serine esterase, round 2-8 designs. "Prospective": Round 6 (Kemp eliminase) or round 8 (serine esterase) designs curated to test the effect of AP Novo filtering. All in this set are MR-filtered, unlike the retrospective set. Black dots: mean $k_{obs}$; red dots: mean $k_{obs}$ of hits. B) $k_{obs}$ distributions of design sequences generated using LigandMPNN and MR-filtered, compared to those generated by AP Novo and MPD-filtered. Data are facets from Kemp eliminase round 6 (LMPNN MR) and round 7 (AP Novo MPD) and serine esterase round 8 (both LMPNN MR and AP Novo MPD). Additional designs generated by AP Novo and MR-filtered are shown in Figure S27B

**Figure S29. Extended filters based on AlphaFold structure predictions and metrics.** A) Example structure of nitrene intermediate illustrating the C5-N-versus-C4-N distance difference metric, which has negative values when the intermediate is more piperidine-like and positive values otherwise. B) Distribution of C5-N-versus-C4-N values (minimum of seven prediction seeds) for round one nitrene transferase designs, split by activity. (left). 'Active' denotes designs yielding >5% of the piperidine peak area of TamPgb wt (rather than total activity, as in Fig. 1D). Piperidine formation in nitrene transferase round two, split by whether designs passed the C-N distance filter (middle). Hit rates and piperidine formation of designs in nitrene transferase round two, as a function of how many resequences a given design passed (right). C) Enrichment factor (precision normalised to baseline) at 80% recall for each of the nine AF3 metrics representing general, reaction-agnostic features (see Supplemental Information for exact metrics used). Labels in parentheses indicate number of hits and total number of designs tested. "SE": (4MU-Ac) serine esterase; "KE": Kemp eliminase; "NT": nitrene transferase. Bottom graph: Cumulative enrichment (solid lines) and recall (dotted lines) from jointly filtering on the metrics in top graph from left to right. All p-values are for a two-sided Mann-Whitney U-test (MWU-p). EF@R50 denotes enrichment factor at 50% recall.

**Figure S30. Combined effects of multi-seed, multi-resequence, and multi-partial-diffusion scoring.** A) Precision (equivalent to retrospective hit rate), recall, and mean $k_{obs}$ of hits versus number of resequences and partial diffusions passing filters for Kemp eliminases (exact filters in Table S7). Different columns show results under different multi-seed filtering approaches. "N/10" indicates a certain number of seeds (or resequences or partial diffusions) pass filters out of 10 attempts. "N/N" indicates a certain number of attempts are evaluated and all pass. "Seed-median" means that a subset of metrics were median-aggregated across seeds before filtering—this is the setting used in design campaigns. The data in column 3 are shown in Fig. 4D as a heatmap. The total number of designs and hits in this analysis are 1,521 and 179, respectively. B) Same as (A) but for serine esterases. In design campaigns, resequences and partial diffusions were considered passing if 1/1 seeds passed (unlike Kemp elimination, which used the median of seeds); this is shown in columns 3 and 6. Column 3 corresponds to the heatmap in Fig. 4E. The total number of designs and hits in this analysis are 1,524 and 113, respectively.

### 2 Supplemental Tables

**Table S1. Michaelis-Menten parameters of best designs for model reactions and comparison with literature.** Errors represent standard deviation. \*KABLE1 uses a well-characterized existing scaffold but is the first Kemp eliminase designed in that scaffold.

| Design | Reaction | Substrate | Scaffold | $k_{\text{cat}}$ ($\text{s}^{-1}$) | $K_{\text{m}}$ ($\mu\text{M}$) | $K_{\text{i}}$ ($\mu\text{M}$) | $k_{\text{cat}}/K_{\text{m}}$ ($\text{M}^{-1}\text{s}^{-1}$) | Notes |
| --- | --- | --- | --- | --- | --- | --- | --- | --- |
| GDM_SE_2754 | Serine esterase | 4MU-Ac | Novel | $0.079 \pm 0.014$ | $16 \pm 5$ | $120 \pm 50$ | $4,900 \pm 700$ | non-NSE topology |
| GDM_SE_2663 | Serine esterase | 4MU-Ac | Novel | $0.44 \pm 0.22$ | $140 \pm 80$ | $7 \pm 4$ | $3,070 \pm 140$ | non-NSE topology |
| GDM_SE_2625 | Serine esterase | 4MU-Ac | Novel | $0.071 \pm 0.005$ | $18.7 \pm 2.1$ | $95 \pm 14$ | $3,820 \pm 180$ | non-NSE topology |
| GDM_SE_2845 | Serine esterase | 4MU-Ac | Novel | $0.0326 \pm 0.0029$ | $12.3 \pm 1.1$ | | $2,660 \pm 230$ | non-NSE topology |
| GDM_SE_2937 | Serine esterase | 4MU-Ac | Novel | $5.2 \pm 1.3$ | $14.7 \pm 2.8$ | | $360,000 \pm 70,000$ | NSE topology |
| GDM_SE_2729 | Serine esterase | 4MU-Ac | Novel | $0.113 \pm 0.008$ | $8.8 \pm 1.3$ | $180 \pm 40$ | $12,800 \pm 1,100$ | NSE topology |
| GDM_SE_1537 | Serine esterase | 4MU-Ac | Novel | $0.113 \pm 0.016$ | $9.9 \pm 2.7$ | $120 \pm 40$ | $11,400 \pm 1,600$ | NSE topology |
| GDM_SE_2878 | Serine esterase | 4MU-Ac | Novel | $0.19 \pm 0.06$ | $15.3 \pm 2.4$ | | $13,000 \pm 4,000$ | NSE topology |
| win1 [3] | Serine esterase | 4MU-Ac | Novel | 0.018 | 110 |  | 170 |  |
| win11 [3] | Serine esterase | 4MU-Ac | Recycled | 0.0197 | 5.0 |  | 3,900 |  |
| momi [3] | Serine esterase | 4MU-PA | Novel | 0.10 | 83 |  | 1,200 |  |
| momi120-103 [3] | Serine esterase | 4MU-PA | Recycled | 0.057 | 0.26 |  | 220,000 |  |
| GDM_KE_1483 | Kemp eliminase | 5-NBO | Novel | $18 \pm 3$ | $1200 \pm 300$ | | $14,600 \pm 1,100$ | pH 7.5 |
| | | | | $18.7 \pm 1.3$ | $940 \pm 30$ | | $19,900 \pm 2,100$ | pH 9.0 |
| | | | | $17.1 \pm 0.9$ | $1,170 \pm 280$ | | $15,300 \pm 2,800$ | pH 10.0 |
| | | | | $6.0 \pm 0.6$ | $500 \pm 90$ | | $12,300 \pm 1,400$ | pH 7.5 |
| | | | | $21.8 \pm 1.5$ | $610 \pm 50$ | | $35,700 \pm 400$ | pH 9.0 |
| GDM_KE_1872 | Kemp eliminase | 5-NBO | Novel | $29.9 \pm 2.5$ | $660 \pm 50$ | | $44,940 \pm 220$ | pH 10.0 |
| | | | | $9.1 \pm 2.0$ | $840 \pm 290$ | | $11,300 \pm 1,500$ | |
| | | | | $1.02 \pm 0.07$ | $160 \pm 30$ | | $6,800 \pm 1,500$ | |
|  |  |  |  | 0.3 | 1300 |  | 213 |  |
|  |  |  |  | 0.07 | 500 |  | 131 |  |
| GDM_KE_1902 | Kemp eliminase | 5-NBO | Novel | 9.1 ± 2.0 | 840 ± 290 |  | 11,300 ± 1,500 |  |
| GDM_KE_1770 | Kemp eliminase | 5-NBO | Novel | 1.02 ± 0.07 | 160 ± 30 |  | 6,800 ± 1,500 |  |
| Des61 [4] | Kemp eliminase | 5-NBO | Novel | 0.3 | 1300 |  | 213 |  |
| Des27 [4] | Kemp eliminase | 5-NBO | Novel | 0.07 | 500 |  | 131 |  |
| Des27.7 [4] | Kemp eliminase | 5-NBO | Recycled | 2.85 | 220 |  | 12,696 |  |
| KABLE1 [5] | Kemp eliminase | 5-NBO | Novel* | 0.030 | 170 |  | 176 | pH 7.5 |
|  |  |  |  | 0.39 | 120 |  | 3,300 | pH 9.0 |
|  |  |  |  | 1.3 | 350 |  | 3,700 | pH 10.0 |
|  |  |  |  | 65 | 380 |  | 170,000 | pH 7.5 |
|  |  |  |  | 210 | 400 |  | 530,000 | pH 9.0 |
| KABLE1.4 [5] | Kemp eliminase | 5-NBO | Recycled | 290 | 490 |  | 610,000 | pH 10.0 |
|  |  |  |  | 210 | 300 |  | 640,000 | pH 7.5 |
|  |  |  |  | 470 | 240 |  | 2,000,000 | pH 9.0 |
| KABLE2.5 [5] | Kemp eliminase | 5-NBO | Recycled | 570 | 270 |  | 2,100,000 | pH 10.0 |

**Table S2. Success rate in the Kemp eliminase multiple-resequencing facet (rounds 5 and 6) by motif.**

| base / oah | N | Q | S | T | Y | Total |
| --- | --- | --- | --- | --- | --- | --- |
| **D** | 2/9 (0.22) | 3/5 (0.60) | 1/9 (0.11) | 3/4 (0.75) | 9/32 (0.28) | 18/59 (0.31) |
| **E** | 0/3 (0.00) | 3/4 (0.75) | 3/4 (0.75) | 1/3 (0.33) | 7/13 (0.54) | 14/27 (0.52) |
| **H** | 0/8 (0.00) | 0/5 (0.00) | 1/14 (0.07) | 3/12 (0.25) | 3/28 (0.11) | 7/67 (0.10) |
| **HD** | 0/2 (0.00) | 0/4 (0.00) | 0/4 (0.00) | 0/2 (0.00) | 2/8 (0.25) | 2/20 (0.10) |
| **Total** | 2/22 (0.09) | 6/18 (0.33) | 5/31 (0.16) | 7/21 (0.33) | 21/81 (0.26) | 41/173 (0.24) |

**Table S3. Sequence novelty assessment.** We assessed the sequence novelty of a selection of our most active designs by conducting searches against the Uniref, Mgnify, and Small BFD databases ([6, 7, 8]). Using JackHMMer ([9]) with an E-value threshold of $10^{-3}$, we identified sequence hits which were then subjected to pairwise global alignment with the design sequences. Sequence identity was determined by calculating the ratio of identical amino acids to the total alignment length.

| Design ID | Max Seq. Identity (%) | Closest Hit |
| --- | --- | --- |
| **Nitrene Transferase** |  |  |
| GDM_NT_0121 | 12.1 | UniRef90_A0A7R9EA81/291-331 |
| GDM_NT_0270 | 18.7 | UniRef90_A0A8T2LSX5/170-244 |
| **DEHPase** |  |  |
| GDM_DEHP_0034 | 13.4 | UniRef90_UPI0028A2E68C/1625-1662 |
| GDM_DEHP_0078 | — | — |
| GDM_DEHP_0176 | 18.8 | UniRef90_A0A257XHU4/170-217 |
| GDM_DEHP_0187 | 15.3 | UniRef90_A0A9X4SQI1/1054-1097 |
| GDM_DEHP_0376 | — | — |
| GDM_DEHP_0382 | — | — |
| **Serine Esterase** |  |  |
| GDM_SE_1537 | — | — |
| GDM_SE_2619 | 19.3 | UniRef90_A0A8S9HB25/1295-1382 |
| GDM_SE_2625 | — | — |
| GDM_SE_2644 | — | — |
| GDM_SE_2663 | 17.2 | UniRef90_UPI001CE041EF/426-486 |
| GDM_SE_2729 | — | — |
| GDM_SE_2754 | — | — |
| GDM_SE_2785 | — | — |
| GDM_SE_2786 | — | — |
| GDM_SE_2800 | — | — |
| GDM_SE_2845 | — | — |
| GDM_SE_2878 | — | — |
| GDM_SE_2937 | — | — |
| GDM_SE_3019 | — | — |
| GDM_SE_3036 | — | — |
| GDM_SE_3043 | — | — |
| **Kemp Eliminase** |  |  |
| GDM_KE_1474 | — | — |
| GDM_KE_1483 | 29.0 | UniRef90_A0A9R0F2X0/3838-3912 |
| GDM_KE_1511 | — | — |
| GDM_KE_1515 | — | — |
| GDM_KE_1529 | 28.5 | ERR1039457_4090143/109-176 |
| GDM_KE_1770 | — | — |
| GDM_KE_1872 | — | — |
| GDM_KE_1902 | — | — |
| **Carbene Transferase** |  |  |
| GDM_CT_0103 | 24.6 | UniRef90_A0A5N5F7J9/1156-1221 |
| GDM_CT_0106 | 28.3 | SRR6516165_155164/13-77 |
| GDM_CT_0131 | 20.5 | A0A2E9QPM4_9DELT/712-789 |
| GDM_CT_0162 | 27.7 | SRR6516162_11924763/158-216 |

### 3 Design Process

The design process begins with the choice of a target reaction along with its known or hypothesised reaction mechanism. This informs the creation of a catalytic motif, or a set of coordinates of amino acid side chains and small molecules which represents a transition state or intermediate of the reaction. The motif is then input to the model along with additional conditioning information to form full inputs (see §3.1). The process of creating specific model inputs involves sampling of ligand conformers, design sequence length and optionally the indices of the motif residues in the sequence. These inputs are provided to a conditional diffusion model which generates a combined sequence and structure design candidate, where the generated structure includes side chain atoms. Optionally, the sequence can be redesigned by a "resequencing" model such as LigandMPNN.

Design candidates are evaluated on a variety of metrics, the majority of which are based on the AlphaFold 3 predicted structure of the designed sequence and target ligand, often across multiple seeds of such predictions. Based on these evaluations a batch is assembled from design candidates and tested *in vitro* for activity on the design goal reaction. The overall pipeline is illustrated in Figure S1.

#### 3.1 Design Inputs

The primary design inputs are coordinates and amino acid/atom types of partial proteins ("motifs") and of ligands. Ligand atoms are considered fixed during generation. The total length of the designed sequence is also included in the model inputs.

##### 3.1.1 Coordinate conditioning

Each residue in the motif can have all atom coordinates specified or only a subset, where the subset may include backbone atoms, sidechain atoms or both. Some specific variants of this where all residues in the motif have a similar setup are used commonly and we name them here:

• *Full Motif*: all residues in the motif have every atom specified other than possibly some residues where we specify only backbone coordinates.
• *Tip Atom*: all residues in the motif have only tip atoms (atoms towards the end of side chains) specified, again other than possibly some residues where we specify only backbone coordinates.
• *Tip Frame*: similar to Tip Atom conditioning, but with larger portions of each side chain specified.

##### 3.1.2 Amino acid type conditioning

Mostly, the amino acid types of residues in the motif are included in the conditioning. However for a subset of residues the conditioning can exclude the amino acid type, leaving the model to choose what type the residue should have. In particular, when only providing backbone coordinates for a residue, one can choose whether to fix the residue type or not. We call motif residues where the amino acid type is not fixed *resequence residues*.

##### 3.1.3 Sequence position conditioning

Finally, we also optionally condition on the positions (index) of each residue of the motif within the sequence. We name two important modes of conditioning:

• *Indexed*: position indices are provided for all residues in the motif.
• *Unindexed*: no position indices are provided for residues in the motif.

### 4 Diffusion Model

The model used to generate design candidates is a conditional diffusion model which accepts certain structural motifs and sequence constraints as conditioning and outputs a designed sequence and structure. The core of the model is a neural network "denoiser" which when sampling designs is run many times to gradually go from pure noise to a final design. In this section we describe the model in detail, including possible conditioning, diffusion sampling, model training and the denoiser network architecture.

### 4.1 Modalities and input representations

The model co-generates all-atom structures and sequences using a diffusion-based framework [10, 11, 12]. Structure diffusion is defined over the ambient coordinate space (i.e. not on any constrained subspace of plausible structures) and includes the coordinates of ligand atoms and sidechain atoms [13, 14].

Sequences are represented as discrete token types, with each protein residue as one token and each ligand atom as one token, similarly to [13, 15]. The vocabulary consists of the 20 canonical amino acid types plus the 44 most common elements represented in ligands in the PDB. Structures are represented by the coordinates of all heavy atoms in the protein and ligand structure. Coordinates are grouped together into tokens in the same way as for sequence, i.e. one token per amino acid residue and one token per ligand atom.

Per token, we represent coordinates using an array of three dimensional coordinates with length 37. For protein residue tokens, the 37 positions correspond to different possible atom types for the standard 20 amino acids (as in [16]); for ligand atom tokens only the first position is relevant, the remaining 36 are padding. For a protein residue with a given amino acid type, for atoms that are not present in that amino acid we introduce virtual atoms with coordinates that are included in the diffusion. During training, the target positions of these virtual atoms have the same coordinates as the $C_\alpha$ atom of that residue. During sampling, the model can place some of the 36 non-$C_\alpha$ atoms per residue near the $C_\alpha$ atom coordinates, then processing at the end of sampling removes atoms within 1Å of $C_\alpha$ and matches remaining atoms against standard amino acids to resolve the sequence. We call this "virtual-all-atom" and note that similar strategies have also been used elsewhere [17, 18, 19].

### 4.2 Notation

We define the following notation for the diffusion model description:

• $L$: The total sequence length, including all ligand atoms.
• $i \in \{1, \dots, L\}$: The sequence position index.
• $a \in \{1, \dots, 37\}$: The atom type index.
• $c \in \{1, 2, 3\}$: The 3D coordinate index.
• $x_{i,a,c} \in \mathbb{R}$: The coordinate/position of an atom at sequence position $i$, atom type $a$, and spatial dimension $c$.
• $\mathbf{x}$: The full structure (atom coordinates), including any conditioning motif and ligand atoms, represented as the collection $\{x_{i,a,c}\}$.
• $s_i \in \{1, \dots, 64, M\}$: The sequence token at position $i$, which is either one of the 20 canonical amino acid types, an unknown amino acid, one of 43 common ligand atom types, or a special mask token $M$.
• $\mathbf{s} := \{s_i\}_{i=1}^L$: The full sequence, including any conditioning motif and ligands.
• $\mathbf{m}^x$: atom mask indicating which atoms are present in data or active in training.
• $\mathbf{m}^s$: sequence mask indicating which motif or ligands are present.
• $\tilde{\mathbf{x}}_i$: generated structure at diffusion step $i$, to distinguish from a generic $\mathbf{x}$ above.
• $\tilde{\mathbf{s}}_i$: generated sequence at diffusion step $i$, to distinguish from a generic $\mathbf{s}$ above.
• $\mathbf{x}^{\text{motif}}$: the structure of the conditioning motif, including all ligand atoms.
• $\mathbf{s}^{\text{motif}}$: the sequence tokens fixed in the conditioning, including ligand atom types.
• $\mathbf{m}^{x,\text{motif}} \in \{0, 1\}^{L \times 37}$: a mask that specifies which atoms in $\mathbf{x}$ are in the structural motif, with $\mathbf{x}[\mathbf{m}^{x,\text{motif}}] = \mathbf{x}^{\text{motif}}$. For unindexed motifs this mask is empty when sampling, but not when training as losses make use of the ground truth index positions.
• $\mathbf{m}^{s,\text{motif}} \in \{0, 1\}^L$: a mask that specifies which elements in $\mathbf{s}$ are fixed, with $\mathbf{s}[\mathbf{m}^{s,\text{motif}}] = \mathbf{s}^{\text{motif}}$. For unindexed motifs this mask is empty when sampling, but not when training as losses make use of the ground truth index positions.
• $\tilde{\mathbf{m}}^{x,\text{motif}}$: motif mask at end of sampling (only relevant if motif is unindexed).

In algorithm boxes below, we use the following shorthands:

• $\{\cdot\}$: grouping a variable with its related tensors, typically masks. For example, $\{\mathbf{x}\}$ groups a structure $\mathbf{x}$ with $\mathbf{m}^x$; $\{\mathbf{s}\}$ groups sequence $\mathbf{s}$ with $\mathbf{m}^s$, and another mask that identifies ligands.
• $\{\mathbf{f}^*\}$: a collection of all motif conditioning tensors, including $\mathbf{x}^{\text{motif}}$, $\mathbf{s}^{\text{motif}}$, $\mathbf{m}^{x,\text{motif}}$, $\mathbf{m}^{s,\text{motif}}$.

• $D_\theta$: the denoiser network with parameters $\theta$.

Finally, we define one constant $\sigma_{\text{data}} := 14.4$ which is used to rescale data during the diffusion process.

#### 4.3 Diffusion process

We use a continuous diffusion process to model structure coordinates and a discrete diffusion process to model sequence tokens. For continuous diffusion we use the EDM framework [20] and for discrete diffusion we use the masked diffusion framework [21, 22, 23]. Briefly, the EDM framework corrupts data by adding Gaussian noise of increasing variance and the masked diffusion framework corrupts data by converting increasing proportions of the sequence to a special mask token.

We use a decoupled-time training framework [e.g., 24, 10, 25, 26] and assign distinct time steps to the sequence and structure modalities, denoted by $t^x$ and $t^s$, respectively. Compared to training with coupled $t^x$ and $t^s$, this approach allows us to optimise how fast we denoise sequence and structure after training, and can have additional generalisation benefits, since the model has seen more combinations of noisy structures and sequences. Structure and sequence space have separate diffusion time steps denoted by $x_{t^x}$ and $s_{t^s}$, respectively, with $x_0=x$ the clean structure and $s_0=s$ the clean sequence. For both modalities, in a single forward process transition, each element (coordinate or token) is sampled conditionally independent of the others: for all $l, a$ and $c$,

$$\begin{aligned} \vec{p}(x_{l,a,c,t^x} | x_{l,a,c,0}, t^x) &:= \mathcal{N}(x_{l,a,c,t^x}; x_{l,a,c}, \sigma_{t^x}^2), & \sigma_{t^x} &= h^x(t^x); \\ \vec{p}(s_{l,t^s} | s_{l,0}, t^s) &:= \begin{cases} s_l & \text{with probability } \alpha_{t^s} \\ M & \text{with probability } 1 - \alpha_{t^s} \end{cases}, & \alpha_{t^s} &= h^s(t^s) \end{aligned} \quad (1)$$

where $M$ is a mask token, and $h^x$ and $h^s$ are noise schedules mapping from a time step to a Gaussian noise standard deviation $\sigma_{t^x} \in (0, \infty)$ for the structure or an unmask probability $\alpha_{t^s} \in (0, 1)$ for the sequence. By convention of the discrete diffusion literature, the term $\alpha_{t^s}$ specifies the *signal* level [e.g., 21, 23], unlike the noise std $\sigma_{t^x}$ in the continuous structure diffusion. To avoid confusion, we refer to $\sigma_{t^x}$ and $1 - \alpha_{t^s}$ as the *noise levels*, and refer to $\sigma_{t^x}$ and $\alpha_{t^s}$ as the *noise terms* of the diffusion processes.

We now detail the schedule functions. Following [20], we set the time step as $t^x \in (0, \infty)$, and $\sigma_{t^x} = t^x$ ($h^x$ as the identity function). The sequence scheduling $h^s$ is defined with more nuance as the following. Let $S_\tau(z) = 1/(1+e^{-z/\tau})$ be the standard sigmoid parametrised by a temperature $\tau > 0$. Define a truncated sigmoid as

$$S_{\tau, z_{\text{start}}, z_{\text{end}}}(t) = \frac{S_\tau(z_{\text{end}}) - S_\tau(z(t))}{S_\tau(z_{\text{end}}) - S_\tau(z_{\text{start}})}. \quad z(t) = z_{\text{start}} + t(z_{\text{end}} - z_{\text{start}}). \quad (2)$$

This schedule follows the sigmoid schedule of [27, Algorithm 1]. This effectively takes a segment of $S$ supported on $[z_{\text{start}}, z_{\text{end}}]$, and rescales it to a function mapping from $[0, 1]$ to $[0, 1]$. We use $z_{\text{start}} = -4$, $z_{\text{end}} = 3$, and $\tau = 0.3$. We further pre-compose $S_{\tau, z_{\text{start}}, z_{\text{end}}}$ with a clipping function:

$$h^s(t) = S_{\tau, z_{\text{start}}, z_{\text{end}}}(\min\{t/0.93, 1.0\}). \quad (3)$$

The sigmoid function is chosen for best performance in our model evaluation from a wide range of noise schedules [27], and the reason for the clipping function is explained in §4.4.1.

### 4.4 Sampling

**Algorithm 1** Sampling given motif inputs, a trained denoiser $D_\theta$, design length $L$ and number of sampling steps $T$.

```
1: def SAMPLE($\{\mathbf{f}^*\}$, $L$, $T$, $D_\theta$):
2:   # Initialise $\tilde{\mathbf{x}}$ independently for each element and set and $\tilde{\mathbf{s}}$ to be all mask token M.
3:   $\tilde{\mathbf{x}} \sim \mathcal{N}(0, \sigma_{t^x}^2)$, $\tilde{\mathbf{s}} = \mathbf{M}$
4:   Initialise the masks in $\{\tilde{\mathbf{x}}\}$ and $\{\tilde{\mathbf{s}}\}$ from $\{\mathbf{f}^*\}$ and $L$.
5:   # Initialise an all-zero predicted structure for the initial self-conditioning.
6:   $\hat{\mathbf{x}}_0 \leftarrow \mathbf{0}$
7:   Setup timesteps $\{t_i^x\}_{i=1}^T, \{t_i^s\}_{i=1}^T$. §4.4.1, Equation (9)
8:   for $i = T : 1$ do
9:     # If motif indexed, replace sampled motif atom coordinates with noised conditioning coordinates. §4.4.2
10:    $\tilde{\mathbf{m}}^{\mathbf{x}, \text{motif}} \leftarrow \mathbf{x}^{\text{motif}} + \zeta$, $\zeta \sim \mathcal{N}(0, \sigma_i^2)$
11:    # If motif indexed, replace sampled motif sequence with conditioning sequence.
12:    $\tilde{\mathbf{s}}[\mathbf{m}^{\mathbf{s}, \text{motif}}] \leftarrow \mathbf{s}^{\text{motif}}$
13:    # Update noise terms.
14:    $\sigma_i \leftarrow t_i^x$, $\alpha_i \leftarrow h^s(t_i^s)$ Equation (3)
15:    # Call the protein denoiser to obtain predictions.
16:    $\hat{\mathbf{x}}_0, \hat{\mathbf{p}}_0 \leftarrow D_\theta(\{\tilde{\mathbf{x}}\}, \{\tilde{\mathbf{s}}\}, t_i^x, t_i^s, \{\hat{\mathbf{x}}_0\}, \{\mathbf{f}^*\})$ Equation (5), Algorithm 3
17:    # Sample the structure using the EDM algorithm.
18:    $\tilde{\mathbf{x}} \leftarrow \text{EDM}(\hat{\mathbf{x}}_0, \sigma_i, D_\theta, \tilde{\mathbf{x}}, t_i^x)$ Equation (6), [20, Algorithm 2]
19:    # Sample the sequence using the masked diffusion sampling algorithm.
20:    $\tilde{\mathbf{s}} \leftarrow \text{MD}(\hat{\mathbf{p}}_0, \tilde{\mathbf{s}}, \alpha_i)$ Equation (7), [21, Algorithm 2]
21:  end for
22:  # Final structure denoising without randomness.
23:  $\hat{\mathbf{x}}_0, \_ = D_\theta(\{\tilde{\mathbf{x}}\}, \{\tilde{\mathbf{s}}\}, t_1^x, t_1^s, \{\hat{\mathbf{x}}_0\}, \{\mathbf{f}^*\})$ Algorithm 3
24:  $\tilde{\mathbf{x}} \leftarrow \hat{\mathbf{x}}_0$
25:  If motif unindexed, identify $\tilde{\mathbf{m}}_i^{\mathbf{x}, \text{motif}}$ by spatial matching of input motif to sampled structure. §4.4.3
26:  If motif indexed, $\tilde{\mathbf{m}}^{\mathbf{x}, \text{motif}} \leftarrow \mathbf{m}^{\mathbf{x}, \text{motif}}$.
27:  return $\tilde{\mathbf{x}}, \tilde{\mathbf{s}}, \tilde{\mathbf{m}}^{\mathbf{x}, \text{motif}}$
```

Algorithm 1 summarises the sampling algorithm given a trained denoiser (training details in §4.5) and specific design inputs (input details in §3.1). Sampling dynamics is governed by a sequence of $T$ time steps $\{(t_i^x, t_i^s)\}_{i=1}^T$, where $t_i^x$ and $t_i^s$ decrease together as $i$ decreases from $T$ to 1. We couple $t^x$ and $t^s$ through a function $g : t^x \in (0, \infty) \mapsto t^s \in (0, 1)$ (see §4.4.1), and set up sampling time steps $\{t_i^x\}_{i=1}^T$ from some large value of $t_T^x$ to $t_0^x \approx 0$.

We denote the sampled structure and sampled sequence at the $i$'th sampling time step by $\tilde{\mathbf{x}}_i$ and $\tilde{\mathbf{s}}_i$, respectively. They approximate the forward diffusion distribution over $\mathbf{x}_{t^x}$ and $\mathbf{s}_{t^s}$. Note that the sampled $\tilde{\mathbf{x}}_i$ and $\tilde{\mathbf{s}}_i$ are indexed by integer indices $i$, while the $\mathbf{x}_{t^x}$ and $\mathbf{s}_{t^s}$ during training are indexed by continuous time steps. In other words, during sampling we identify $\tilde{\mathbf{x}}_i$ with $\tilde{\mathbf{x}}_{t_i^x}$, and $\tilde{\mathbf{s}}_i$ with $\tilde{\mathbf{s}}_{t_i^s}$.

To sample a protein, we randomly initialise structure from a zero-mean Gaussian distribution with a large variance, and initialise all sequence tokens to the mask token $\mathbf{M}$. We then iteratively diffuse backwards $T$ steps to obtain $\tilde{\mathbf{x}}_0$, which is approximately distributed as the clean data distribution over $\mathbf{x}_0$. Each denoising sampling step uses the outputs of the trained protein denoiser $D_\theta$, which predicts clean structure $\hat{\mathbf{x}}_0$ and the probabilities of the true amino acid types for masked tokens $\hat{\mathbf{p}}_0$ (shape $L \times 64$; log probabilities over 64 sequence alphabet, with length $L$) given noisy $\mathbf{x}_{t^x}$ and $\mathbf{s}_{t^s}$, conditioned on the time steps $t^x$ and $t^s$:

$$\hat{\mathbf{x}}_0, \hat{\mathbf{p}}_0, \dots = D_\theta(\mathbf{x}_{t^x}, \mathbf{s}_{t^s}, t^x, t^s, \dots). \quad (4)$$

For simplicity, here we omit other inputs and outputs of the denoiser (e.g., motif, self-conditioning, see §4.6).

At each sampling step, the predicted $\hat{\mathbf{x}}_0$ and $\hat{\mathbf{p}}_0$ based on $\tilde{\mathbf{x}}_i$ and $\tilde{\mathbf{s}}_i$ is used to parameterise the reverse sampling distribution:

$$\hat{\mathbf{x}}_0, \hat{\mathbf{p}}_0, \dots = D_\theta(\tilde{\mathbf{x}}_i, \tilde{\mathbf{s}}_i; t_i^x, t_i^s, \dots), \quad (5)$$

$$\tilde{\mathbf{x}}_{i-1} | \tilde{\mathbf{x}}_i, \tilde{\mathbf{s}}_i, i \sim \text{EDM}(\hat{\mathbf{x}}_0, \sigma_{t_i^x}, D_\theta, \tilde{\mathbf{x}}_i, t_i^x), \quad (6)$$

$$\tilde{\mathbf{s}}_{i-1} | \tilde{\mathbf{x}}_i, \tilde{\mathbf{s}}_i, i \sim \text{MD}(\hat{\mathbf{p}}_0, \tilde{\mathbf{s}}_i, \alpha_{t_i^s}), \quad (7)$$

where $\sigma_{t_i^x} = t_i^x$ and $\alpha_{t_i^s} = h^s(g(t_i^x))$ depend on the noise schedules in Equation (1) and the time coupling function $g$ to be detailed in Equation (8). The EDM sampler in Equation (6) uses Algorithm 2 by [20], and involves a churn step that requires making a prediction at slightly increased noise level, using $D_\theta$, $\tilde{\mathbf{x}}_i$, and $t_i^x$ in gray color, see [20]. The MD (masked diffusion) sampler implements Algorithm 2 by [21]. We additionally apply structure self-conditioning [28]: the denoised coordinate prediction from the previous time step serves as $\mathbf{x}^{\text{self}}$ for the current step, except for the first step where $\mathbf{x}^{\text{self}}$ are zeros.

##### 4.4.1 Sampling time steps and coupling

The sampling time steps $\{t_i^x\}_{i=1}^T$ and $\{t_i^s\}_{i=1}^T$ are chosen according to the training time step distributions $p_{\text{train}}^x$ and $p_{\text{train}}^s$ described in §4.5.2, as well as performance optimisation. We did not use the default schedule of EDM sampler, as we reasoned that it is important to follow the training time distributions in multi-modal diffusion modelling. Given that $p_{\text{train}}^x$ is log-normal, and $p_{\text{train}}^s$ is uniform, a natural choice of coupling is the CDF of $p_{\text{train}}^x$:

$$t^s = g(t^x) = \mathcal{F}_{\text{LN}(1.7, 2.0^2)}(t^x). \quad (8)$$

We still need to specify the starting time steps at $i = T$ and end time steps $i = 1$. During diffusion model development, we found that an initial $\sigma_{t_T^s} = 100$ and final $\sigma_{t_1^x} = 0.01$ to be optimal. The value $\sigma_{t_T^s} = 100$ implies that $t_T^s = \mathcal{F}_{\text{LN}(1.7, 2.0^2)}(100) \approx 0.926$, which requires the sequence noise term $\alpha_{t^s} = h^s(t^s)$ in Equation (1) to be zero when $0.926 \leq t^s \leq 1$, to correspond to a fully masked initial sequence. Since the noise schedule is a monotonically increasing function from $[0, 1] \rightarrow [0, 1]$, we use the clipping function in Equation (3): it makes sure that, during sampling when diffusion starts at $\sigma_{t_T^x} = 100$, the sequence noise level $1 - \alpha_{t_T^s} = 1$; and during training, the sequence is fully masked with $1 - \alpha_{t^s} = 1$ whenever $t^x > 100$. Finally, we discretise the noise level $\sigma_t$ by setting up uniform quantiles between $\sigma_{t_T^x}$ and $\sigma_{t_1^x}$ under the training log-normal distribution. Taken together, the sampling time steps are as follows:

$$\begin{aligned} \sigma_{t_1^x} &= 0.01, \quad \sigma_{t_T^x} = 100.0, \quad T = 1000, \quad \# \text{ Empirically optimised structure noise limits} \\ u_1 &= \mathcal{F}_{\text{LN}(1.7, 2.0^2)}(\sigma_{t_1^x}), \quad u_T = \mathcal{F}_{\text{LN}(1.7, 2.0^2)}(\sigma_{t_T^x}), \quad u_i = u_1 + (u_T - u_1)(i - 1)/(T - 1), \quad \# \text{ Coupling variable} \\ \sigma_{t_i^x} &= t_i^x = \mathcal{F}_{\text{LN}(1.7, 2.0^2)}^{-1}(u_i) \in [0.01, 100], \quad t_i^s = u_i \in [0, 1]. \quad \# \text{ Structure and sequence time steps} \end{aligned} \quad (9)$$

The time steps $t_i^x$ and $t_i^s$ are then coupled through the coupling variable $u_i$ with $t_i^s = g(t_i^x)$ for all $i$. The resulting trajectory of noise levels $\sigma_{t_i^x} = t_i^x$ and $1 - \alpha_{t_i^s} = 1 - h^s(t_i^s)$ during sampling are shown in Figure S31 (left).

##### 4.4.2 Replacement with indexed motif conditioning

When the motif is indexed, at each sampling iteration, we replace the motif parts of the sampled $\tilde{\mathbf{x}}$ and $\tilde{\mathbf{s}}$ with conditioning structure and sequence information before sending $\tilde{\mathbf{x}}$ and $\tilde{\mathbf{s}}$ to the denoiser. This makes sure that the motif is visible through the attention mechanism of the denoiser, in addition to through the conditioning embeddings, and ensures that the samples always contain the correct motif properties. For structure, we replace with atom coordinates with added Gaussian noise as done during training to avoid distribution shift. For sequence, we replace with the full true motif sequence.

##### 4.4.3 Unindexed motif conditioning

When the motif is unindexed, the model does not automatically assign residue indices for the generated motif. In fact, the final sample is not guaranteed to contain all residues of the input motif. To identify the motif, we geometrically match residues from the input motif to those in the sampled structure, constraining for matching sequence tokens and therefore atom naming. Specifically, we use the Hungarian algorithm to identify the optimal residue assignment based on the distances between the matched atoms as done in RFdiffusion2 [29].

#### 4.5 Training and data

The denoiser loss is defined in §4.5.1. The loss function measures how well the denoiser can predict clean structure and sequence in the training dataset, given noisy structures and sequences sampled at independent noise levels (§4.5.2). Algorithm 2 summarises the loss computation. The parameters are updated using gradients of the loss function with respect to model parameters using automatic differentiation. We apply gradient clipping and learning rate scaling as detailed in §4.5.3 before each parameter update. Our training data is a mixture of PDB and distillation dataset (§4.5.4), processed according to a set of criteria (§4.5.6). During each training step, each independently sampled protein goes through stages of preprocessing (§4.5.7 and 4.5.8) before being featurised into the model inputs described in §4.6.1.

**Figure S31.** Left, sampling noise levels down-sampled by a factor of 10; arrows indicate direction of decreasing noise levels during sampling. The gray cross indicates the initial noise levels of multi-partial-diffusion. Right, sample noise levels in training.

**Algorithm 2** Loss function for denoiser, used for iterative gradient update of model parameters $\theta$. Inputs are drawn from the data mixture, and have undergone preprocessing stages described in § 4.5.6 to 4.5.8; the resulting features are detailed in § 4.6.1.

```
1: def DENOISERTRAININGLOSS($\{\mathbf{x}\}, \{\mathbf{s}\}, \{\mathbf{f}^*\}; \theta$):
2:   # Sample decoupled diffusion time steps for training.
3:   $t^x \sim p_{\text{train}}^x = \text{LN}(1.7, 2.0^2), \quad t^s \sim p_{\text{train}}^s = \mathcal{U}(0, 1)$ Equations (15) and (16)
4:   # Sample noisy data given time steps.
5:   $x_{\ell,a,c,t^x} \sim \vec{p}(x_{\ell,a,c,t^x} | x_{\ell,a,c}, t^x), \quad s_{\ell,t^s} \sim \vec{p}(s_{\ell,t^s} | s_{\ell}, t^s), \quad \forall \ell, a, c,$ Equation (1)
6:   # Initialise an all-zero self-conditioning structure (no self-conditioning).
7:   $\mathbf{x}^{\text{self}} \leftarrow \mathbf{0}$
8:   # Call the denoiser to obtain predictions
9:   $\hat{\mathbf{x}}_0, \hat{\mathbf{p}}_0, \hat{\mathbf{m}}^{\text{mi}}, \beta^x, \beta^s \leftarrow D_{\theta}(\{\mathbf{x}_t\}, \{\mathbf{s}_t\}, t^x, t^s, \{\mathbf{x}^{\text{self}}\}, \{\mathbf{f}^*\})$ Algorithm 3
10:  $\mathbf{x}^{\text{self}} \leftarrow \text{STOPGRADIENT}(\hat{\mathbf{x}}_0)$
11:  if RAND < 0.8 then
12:    # Call the denoiser again to obtain self-conditioned predictions.
13:    $\hat{\mathbf{x}}_0, \hat{\mathbf{p}}_0, \hat{\mathbf{m}}^{\text{mi}}, \beta^x, \beta^s \leftarrow D_{\theta}(\{\mathbf{x}_t\}, \{\mathbf{s}_t\}, t^x, t^s, \{\mathbf{x}^{\text{self}}\}, \{\mathbf{f}^*\})$ Algorithm 3
14:  end if
15:  # Obtain motif and non-motif predictions using masks. § 4.5.1
16:  $\hat{\mathbf{x}}_0^{\text{motif}} \leftarrow \hat{\mathbf{x}}_0 \odot \mathbf{1}[\mathbf{m}^{\text{x}, \text{motif}} = 1], \quad \hat{\mathbf{p}}_0^{\text{motif}} \leftarrow \hat{\mathbf{p}}_0 \odot \mathbf{1}[\mathbf{m}^{\text{s}, \text{motif}} = 1]$
17:  $\hat{\mathbf{x}}_0 \leftarrow \hat{\mathbf{x}}_0 \odot \mathbf{1}[\mathbf{m}^{\text{x}, \text{motif}} = 0], \quad \hat{\mathbf{p}}_0 \leftarrow \hat{\mathbf{p}}_0 \odot \mathbf{1}[\mathbf{m}^{\text{s}, \text{motif}} = 0]$
18:  # Obtain ground-truth mask indicating if any motif is present at any residual index.
19:  $\mathbf{m}^{\text{mi}} \leftarrow \text{ANY}(\mathbf{m}^{\text{x}, \text{motif}}, \text{AXIS} = \text{atom type})$ $\mathbf{m}^{\text{mi}} \in \{0, 1\}^L$
20:  # Compute the loss Equation (14)
21:  $\mathcal{L}_{\theta}^{\text{total}} \leftarrow \mathcal{L}_{\theta}^x(\hat{\mathbf{x}}_0, \mathbf{x}) + 0.3\mathcal{L}_{\theta}^s(\hat{\mathbf{p}}_0, \mathbf{s}) + 1.0\mathcal{L}_{\theta}^{\text{mi}}(\hat{\mathbf{m}}^{\text{mi}}, \mathbf{m}^{\text{mi}}) + 0.75\mathcal{L}_{\theta}^x(\hat{\mathbf{x}}_0^{\text{motif}}, \mathbf{x}^{\text{motif}}) + 1.0\mathcal{L}_{\theta}^s(\hat{\mathbf{p}}_0^{\text{motif}}, \mathbf{s}^{\text{motif}})$
22:  return $\mathcal{L}_{\theta}^{\text{total}}$
```

##### 4.5.1 Losses

Given the structure and sequence of a cropped training example with sequence length $L$ (see Sections 4.5.6 and 4.5.7 for creation of training examples), we compose a training data tuple as

$$\mathcal{D}_{\mathbf{x}, \mathbf{s}} := \{t^x, \mathbf{x}_{t^x}, t^s, \mathbf{s}_{t^s}, \mathbf{x}, \mathbf{m}^x, \mathbf{s}, \mathbf{m}^s, L\} \quad \text{where } t^x \sim p_{\text{train}}^x, \mathbf{x}_{t^x} \sim \vec{p}(\mathbf{x}_{t^x} | \mathbf{x}, t^x), t^s \sim p_{\text{train}}^s, \mathbf{s}_{t^s} \sim \vec{p}(\mathbf{s}_{t^s} | \mathbf{s}, t^s). \quad (10)$$

The training time step distributions $p_{\text{train}}^x$ and $p_{\text{train}}^s$ are given by Equations (15) and (16), respectively, in § 4.5.2 later. The denoiser network $D_{\theta}$ takes this tuple and outputs the predicted structure $\hat{\mathbf{x}}_0$, predicted sequence $\hat{\mathbf{s}}_0$, predicted motif indices $\hat{\mathbf{m}}^{\text{mi}}$ and uncertainties $\beta^x$ and $\beta^s$ [30, 31]; see Algorithm 3 for details on the neural architecture of the denoiser. For clarity, in the loss functions below we make explicit the dependence of the predicted quantities on the input $\mathcal{D}_{\mathbf{x}, \mathbf{s}}$

and denoiser parameters $\theta$: The structure loss is

$$\mathcal{L}_\theta^x(\hat{\mathbf{x}}_0, \mathbf{x}, \mathbf{m}^x) := \frac{1}{L} \frac{\sigma_{\text{data}}^2 + \sigma_{t^x}^2}{\sigma_{\text{data}}^2 \cdot \sigma_{t^x}^2} \frac{1}{\beta^x(t^x, t^s; \theta)} \left( \sum_{\ell=1}^L \sum_{a=1}^{37} m_{\ell,a}^x \text{Huber}[(\hat{\mathbf{x}}_{0,\ell,a}(\mathcal{D}_{\mathbf{x},s}; \theta) - \mathbf{x}_{\ell,a})] \right) + \log(\beta^x(t^x, t^s; \theta)), \quad (11)$$

where $\sigma_{\text{data}} = 14.4$ is the standard deviation of the atom coordinates in the training data, the Huber loss has a quadratic-to-linear threshold of 20, and the structure uncertainty $\beta^x$ depends only on the time steps $t^x$ and $t^s$ [30, 31]. The structure loss is aggregated over atoms indicated by the atom mask $\mathbf{m}^x$. In the virtual-all-atom representation, we make all 37 atom types active (=1) for all residuals present in the training data, and make the first atom active but the rest 36 inactive for ligand atoms.

The sequence loss is

$$\mathcal{L}_\theta^s(\hat{\mathbf{p}}_0, \mathbf{s}, \mathbf{m}^s) := \frac{1}{L} \frac{\alpha'_{t^s}}{1 - \alpha_{t^s}} \sum_{\ell=1}^L m_\ell^s \cdot \delta[s_{\ell,t^s} = \text{M}] \cdot \text{SCE}[\hat{\mathbf{p}}_{0,\ell}(\mathcal{D}_{\mathbf{x},s}; \theta), s_\ell, \beta^s(t^x, t^s; \theta)], \quad (12)$$

where $\alpha'_t$ is the first derivative of the schedule $h^s$ evaluated at $t$, $\delta$ is the Kronecker delta function that restricts sequence loss to masked tokens only, SCE is the cross-entropy loss with scaled logits,

$$\text{SCE}[\hat{\mathbf{p}}, s, \beta] := -\log \text{softmax}(\hat{\mathbf{p}}/\beta)_s, \quad (13)$$

and $\beta^s$ is the sequence uncertainty that inversely scales the logits of the sequence prediction [30, Equation 8]. The denoised sequence $\hat{\mathbf{p}}_{0,\ell}$ is a vector of logits, and $s_\ell$ is an integer that indexes the log probability vector. The sequence mask $\mathbf{m}^s$ is a vector of $L$ 1's, but we still use this mask to help define the motif loss later.

To train the denoiser for self-conditioned structure sampling [28], we perform a first forward pass on the denoiser with $\mathbf{x}^{\text{self}}$ set to zeros, to obtain the first predictions. Then, with probability 0.2, these predictions are used for loss and gradient computations; with probability 0.8, we apply stop-gradient to the denoised coordinates $\hat{\mathbf{x}}_0$ to obtain $\mathbf{x}^{\text{self}}$, which is sent to the denoiser for a second pass, and we use the second prediction for loss and gradient computations (Algorithm 2).

The model additionally predicts $\hat{\mathbf{m}}_\theta^{\text{mi}} \in \mathbb{R}^L$, the binary logits (log probability ratio) of whether each residue is part of the motif. The target residue index mask $\mathbf{m}^{\text{mi}}$ is 1 if any atom in the corresponding residue is in $\mathbf{m}^{\text{x,motif}}$ and 0 otherwise. A cross-entropy loss $\mathcal{L}_\theta^{\text{mi}}(\hat{\mathbf{m}}_\theta^{\text{mi}}, \mathbf{m}^{\text{mi}})$ between the predicted logits and the true motif residue index mask serves as an auxiliary training signal that encourages the model to learn motif placement. These predicted indices are not used at sampling; instead, motif placement is determined by geometric matching after sampling (§4.4).

We also include auxiliary losses for the model predictions of motif coordinates and sequence tokens to improve performance. We first define $\mathbf{x}_{\text{expanded}}^{\text{motif}}$ to be an expanded version of $\mathbf{x}^{\text{motif}}$ that includes all atoms from sequence positions with any atom in $\mathbf{x}^{\text{motif}}$. Then we define $\hat{\mathbf{x}}^{\text{motif}}$ as the model prediction of all atom coordinates in $\mathbf{x}_{\text{expanded}}^{\text{motif}}$ and $\hat{\mathbf{p}}^{\text{motif}}$ as the model prediction of all sequence tokens in $\mathbf{s}^{\text{motif}}$. The losses $\mathcal{L}^x(\hat{\mathbf{x}}^{\text{motif}}, \mathbf{x}^{\text{motif}}, \mathbf{m}^{\text{x,motif}})$ and $\mathcal{L}^s(\hat{\mathbf{p}}^{\text{motif}}, \mathbf{s}^{\text{motif}}, \mathbf{m}^{\text{s,motif}})$ are computed in the same way as the denoising losses above, but restricted only to the motif sequence positions. Note that ligand atoms are included in the motif structure loss.

The final training objective loss is

$$\mathcal{L}_\theta^{\text{total}}(\mathcal{D}_{\mathbf{x},s}) := \mathcal{L}_\theta^x(\hat{\mathbf{x}}_0, \mathbf{x}, \mathbf{m}^x) + 0.3\mathcal{L}_\theta^s(\hat{\mathbf{p}}_0, \mathbf{s}, \mathbf{m}^s) + 1.0\mathcal{L}_\theta^{\text{mi}}(\hat{\mathbf{m}}_\theta^{\text{mi}}, \mathbf{m}^{\text{mi}}) + 0.75\mathcal{L}_\theta^x(\hat{\mathbf{x}}^{\text{motif}}, \mathbf{x}^{\text{motif}}, \mathbf{m}^{\text{x,motif}}) + 1.0\mathcal{L}_\theta^s(\hat{\mathbf{p}}^{\text{motif}}, \mathbf{s}^{\text{motif}}, \mathbf{m}^{\text{s,motif}}). \quad (14)$$

##### 4.5.2 Time sampling

As is typical for diffusion-based models we perform simulation-free training by sampling intermediate noisy samples from the marginal distributions in Equation (1) (instead of explicitly running some forward transition iteratively). The distributions from which diffusion time steps $t^x$ and $t^s$ are sampled during training has been shown to have a strong effect on training quality [e.g., 20, 32, 11]. The structure time is sampled according to a log-normal distribution with parameters mean = 1.7 and std = 2.0 (for the underlying Gaussian distribution).

$$t^x \sim p_{\text{train}}^x := \text{LN}(1.7, 2.0^2) \quad (15)$$

The sequence time step is sampled from a uniform distribution in (0, 1):

$$t^s \sim p_{\text{train}}^s := \mathcal{U}(0, 1). \quad (16)$$

During training, the structure and sequence time are sampled independently from each other, which we find to have a beneficial effect on performance and generalisation without sacrificing capacity. The time steps sampled this way map to structure and sequence noise terms through $\sigma_{t^x} = h^x(t^x)$ and $\alpha_{t^s} = h^s(t^s)$ in Equation (1), respectively.

#### 4.5.3 Optimisation

The model was trained on 32 Ironwood TPUs, each of which has 2 TensorCores, resulting in a total of 64 cores. We use a batch size of 8 per core, giving a total batch size per step of 512. Training runs last for $\sim 10^6$ steps, which takes $\sim 5$ days.

We use an Adam optimiser ($b_1 = 0.9$, $b_2 = 0.999$) [33], with learning rate modulated by a linear warm-up from 0 to 0.001 in 1000 gradient steps and an exponential decay $\exp(-\text{gradient\_step} \times 512 / (5 \times 10^8))$. The global norm of the gradient per core is clipped at 0.1 for stability. For each parameter update, we also update an exponential-moving-averaged parameter with decay exponent 0.999 and these parameters are used for inference.

#### 4.5.4 Data sources

Our training dataset comprises of Protein Data Bank (PDB) entries released prior to 12th January 2023 and AlphaFold2 distillation data derived from UniRef90 sequences (see Section 4.5.5 for details). PDB and distillation structures are sampled at ratios of 60% and 40% respectively.

#### 4.5.5 Distillation set creation

The Uniref90 distillation dataset consists of AF2 [16] predicted structures from a subset of the Uniref90 dataset. The version of Uniref90 used was 2021\_04, filtered to sequences with at least 16 and at most 1280 residues, resulting in $\sim 135\text{M}$ sequences. Structures were predicted using model the AF2 version “model\_2\_ptm”, with template search over PDB structures as of 2021-02-15, and with MSA search over Uniref90 version 2020\_03., MGnify version 2018/12 and “Small BFD” (described in [8] and available as part of the AF2 release[34]). Predicted structures were relaxed via constrained relaxation, following the same procedure as in AlphaFold2 (OpenMM v.7.3.1 with the Amber99sb force field, see [16]).

Following structure prediction and relaxation, further filters were applied, where SASA and core percentage calculations were performed using Rosetta [35, 36]:

• Mean pLDDT $> 80.0$
• (Sum of hSASA for hydrophobic residues) / (Sum of SASA for all residues) $< 0.25$
• (Sum of [Non-polar surface area minus hSASA] for hydrophobic residues) / (Sum of SASA for all residues) $> 0.44$
• Core layer percentage $> 1\%$

After application of these filters, $\sim 60\text{M}$ structures remained in the distillation set.

Values for the non-polar surface area of hydrophobic residues were taken from the supplementary material of [37] and are: {A: 111, F: 210, G: 92, L:183, M: 208, V: 157, W: 222, Y:177}.

#### 4.5.6 Data processing

PDB entries are sampled using the clusters defined in the PDB supplied file `clusters-by-entity-40.txt` downloaded on 16th October 2023 - a cluster is sampled uniformly then an entry is chosen uniformly from within the cluster. Distillation structures are sampled uniformly from the distillation set.

PDB entries are filtered, leaving only entries with less than 20 chains (pre-bioassembly), where every chain in the entry is either a protein chain or a non-polymer chain. A random bioassembly is chosen and the entry is expanded. Non-polymer chains with the following CCD codes are removed: {UNL, D8U, HOH, DOD}.

The training data processing includes some further chain-level filtering designed to allow for evaluating holdout performance of a much earlier version of the model, but was irrelevant for the development of the current model; for completeness, we describe this filtering now. For both PDB entries and distillation set structures, we remove chains which have $>40$ maximum sequence bit score in a JackHMMER ([9]) search against all chains in a set of monomer structures consisting of the monomers from CASP13 and CASP14 along with a more recent set of small monomers from PDB. Although we include PDB structures released up until 12th January 2023 in training, this filtering was only applied to PDB structures released prior to 29th August 2019. In total this filters out 7118 chains from PDB and 998873 chains from the monomer distillation set

Finally, if the resulting complex has no chains, greater than 20 chains, no protein chains or any protein chains with $>80\%$ of one amino acid type, we skip the complex and resample from the data source.

#### 4.5.7 Cropping strategies

Each protein structure undergoes spatial cropping to a crop size of 512 or less tokens during training (recall that a token is either a whole protein residue or a single ligand atom). We choose a token to center the crop around, take the 512 nearest tokens in space to form an initial crop, then potentially filter out certain chains from that crop and potentially reject the example because the crop does not meet certain conditions. Distances to determine the initial crop are defined using the single ligand atom for ligand tokens and the backbone nitrogen atom for protein residue tokens.

The crop center token is a random token from the whole example if there are no ligands in the example. If ligands are present, with probability 99% we choose a random ligand atom to be the center and with probability 1% we choose a random token from all available tokens instead. If any protein chain present in the initial crop of 512 tokens has less than 50 tokens remaining, we remove it, and if any ligand is not completely included in the initial crop of 512 tokens, we remove it. Finally, if the crop contains more than nine ligand chains or more than 64 ligand atoms, if using the ligand centered cropping we retry with the random spatially centered cropping, otherwise the training example is rejected and we resample from the data source.

#### 4.5.8 Training motif selection

To train the model on a diverse range of conditional generation tasks, we employ various strategies to select motifs from the ground-truth protein. When ligand atoms are present, they are always provided as conditioning. We proceed by selecting which subset of protein residues to include in the motif, then which atoms from those residues to include, then whether to include the residue types in the conditioning and finally whether to provide the motif through the unindexed pathway, the indexed pathway or both pathways.

**Selecting motif residues.** Motif residues for each cropped training example are chosen using one of the following sampling schemes. Schemes are selected with probability proportional to their sampling weight $w$. Strategies that are inapplicable to the current training example are excluded, such as strategies requiring ligand atoms when the current example is from the protein monomer distillation dataset, or the chain inpainting strategy for single chain protein structures.

• **Unconditional Generation** ($w = 2.0$): No residues are included in the motif, training the model to generate structures without conditioning.
• **Random Subset** ($w = 1.0$): Each residue is included in the motif independently with probability $p$, where $p$ is $2^{-U}$ with $U$ a uniform random variable over $(0, \log_2 L)$ where $L$ is the number of tokens in the crop. This is equivalent to $p$ having the loguniform distribution over $(1/L, 1)$.
• **Contiguous Segments** ($w = 1.0$): $p$ is chosen as for a random subset, then a single continuous segment of residues with length $\lceil pL \rceil$ is selected, where $L$ is again the number of tokens in the crop.
• **Chain Inpainting** ($w = 1.0$): A randomly selected protein chain is selected, then every residue not in that chain is fixed. *Requires $\geq 2$ protein chains.*
• **Discontiguous Spatial Triplets** ($w = 1.0$): A random triplet is selected from triplets of residues that are spatially proximal ($C_\beta$ distance $< 6 \text{ \AA}$) but separated by at least 10 residues in the sequence. If not such triplet exists, no fixed residues are chosen. This is similar to the triplet selection in [38].
• **Sparse Pocket Residues** ($w = 2.0$): A random subset of 0 to 4 residues is chosen from the set of residues where at least one atom in the residue is within $4 \text{ \AA}$ of any ligand. *Requires ligand atoms.*
• **Backbone Atoms** ($w = 0.2$): If this strategy is chosen, another strategy is sampled using the other $w$ values, and in addition to the chosen residues, all protein backbone atoms are set to be fixed as part of the later fixed atom sampling.

**Selecting motif atoms.** If the Backbone Atoms strategy was chosen as the initial residue sampling strategy, then all backbone atoms are selected. For four of the other motif residue selection strategies, with some probability we sub-sample the atoms which are fixed for each residue of the motif to simulate the tip-atom conditioning mode, otherwise all atoms for the chosen motif residues are included. Subsampling is done for 2% of Random Subset examples, 2% of Contiguous Segments examples, 10% of Discontiguous Spatial Triplets, and 10% of Sparse Pocket Residues. Subsampling first selects a number of atoms per motif residue then applies some sampled strategy to decide which atoms those should be. The number of atoms subsampled per residue is sampled independently – if the residue contains $n$ atoms then $k \sim \text{Binomial}(n, 0.2)$ atoms are subsampled. With 25% probability, per residue one of the following strategies is independently sampled to decide which $k$ atoms to select; otherwise, a single strategy is sampled and used for all residues.

• **Sidechain Tip** ($w = 12$): Prioritise outermost atoms based on IUPAC Greek-letter depth (e.g., $\gamma$, $\delta$, $\epsilon$), selecting randomly if more atoms at current depth than remain to be chosen to get a total of $k$ atoms.
• **Random Atom Subset** ($w = 3$): Choose atoms uniformly at random from the entire residue.
• **Ligand Proximity** ($w = 48$ if fixed residue strategy was *Sparse Pocket Residues* else $w = 0$): Choose the $k$ atoms closest to the nearest ligand atom.
• **Backbone Subset** ($w = 4$): Choose a random subset of backbone atoms (N, C$_{\alpha}$, C) by choosing a uniform number from 1 to 3 then choosing that many backbone atoms randomly (this strategy ignores the original sampled $k$ value).

**Sequence token conditioning.** Ligand atom types are always included in the conditioning information, but sometimes the amino acid type for protein residues in the motif are not fixed, with an independent random decision per motif residue. If any sidechain atom in the residue is included in the structure motif then the amino acid type is fixed, if not then with probability 50% the amino acid type is fixed and otherwise it is not fixed. This prepares the model for sampling with motifs where some of the residues are resequence residues (see §3.1).

**Selecting from indexed and unindexed pathways.** Motif conditioning can be provided to the model via an indexed token pathway (using the *scaffold* tokens described in §4.6) or via an unindexed token pathway (the *motif* tokens described in §4.6), or both. During training we always have access to the ground truth motif indices, so can choose which pathway to use, randomly selecting from one of the following three options (with one exception to the probabilities if the Backbone Atoms strategy was chosen as the initial residue sampling strategy).

• With probability 0.5, protein motif conditioning is provided only via the unindexed motif token pathway and ligand motif conditioning is provided via both pathways. This is the same as what we do when providing an unindexed motif when sampling.
• With probability 0.25, protein and ligand conditioning is provided via both scaffold and motif tokens. This is the same as what we do when providing an indexed motif when sampling.
• With probability 0.25, or always if the Backbone Atoms strategy was chosen as the initial residue sampling strategy, protein and ligand motif conditioning is provided only via the indexed pathway. This setup is not used when sampling.

Note that full indexed masks $\mathbf{m}^{\text{x,motif}}$ and $\mathbf{m}^{\text{s,motif}}$ are always retained during training for loss computation in Equation (14).

**Maximum motif token size during training and ordering** The number of input motif tokens for the unindexed pathway is at most 128 during training, with protein residue tokens going first, in random shuffle order, then ligand atom tokens, up to a maximum of 128 pathways.

### 4.6 Denoiser network

The denoiser neural network is a transformer based architecture outlined in Figure S32 and detailed in Algorithm 3 and Section 4.6.2. Table S4 lists the key architectural hyperparameters. The total number of trainable parameters in the denoiser network is 147.5 million.

Before describing the denoiser in detail, we define the concept of scaffold and motif token sets. The primary token set, which we call the *scaffold* token set, is the one containing the design complex comprising the main protein and ligand features. This token set is used to create the initial single and pair embeddings, which correspond to single tokens and pairs of tokens respectively. The single embedding is updated via the transformer stack and eventually projected to the denoised coordinates and sequence logits, and the pair embeddings are used to bias the transformer self-attention. Motif information is included in a second token set, which we call the *motif* token set. This information is used to update the single embeddings via cross-attention. The motif token set processing also uses both single and pair embeddings, again with the pair embeddings used to bias transformer self-attention for the single embeddings prior to their use in the cross-attention. Motif information can be included additionally in the initial single embeddings for the scaffold tokens if the motif is indexed (see §4.5.8 for how this is diversified during training).

**Table S4.** Architecture hyperparameters.

| Symbol | Description | Value |
| --- | --- | --- |
| $C$ | Transformer hidden dimension | 512 |
| $C_y$ | Single conditioning dimension | 192 |
| $C_z$ | Pair conditioning dimension | 16 |
| $H$ | Number of attention heads | 8 |
| $d_k$ | Attention key/query dimension per head | 32 |
| $d_v$ | Attention value dimension per head | 32 |
| $N$ | Number of transformer blocks | 32 |
| $n_f$ | SwiGLU transition expansion factor ($C_{\text{int}} = n_f \cdot C$) | 2 |
| $C_\epsilon$ | Noise embedding dimension (per noise type) | 256 |
| $d_{\text{max}}$ | Maximum relative position offset | 64 |
| $n_{\text{bins}}$ | Number of distogram bins | 15 |
| Dropout rate | Applied after attention and transition outputs | 0.3 |

##### 4.6.1 Denoiser inputs

The model operates on a set of features constructed from the tokenised design inputs. These features encode the identity, spatial coordinates, and conditioning state of each token, and are organised into several groups: token-level features describing sequence identity and chain membership; atom-level features providing all-atom 3D coordinates; conditioning features specifying which residues and atoms are fixed during generation; and, when applicable, unindexed motif features that supply the motif through the cross-attention pathway. Table S5 lists the complete set of model input features.

**Figure S32. Denoiser Network Overview.** Yellow boxes represent inputs, blue boxes represent network components (many of which are shown in more detail in later figures), white boxes represent intermediate representations and green boxes represent outputs. Note that the predicted motif indices and uncertainty outputs are only relevant for training, not for sampling.

**Table S5.** Input features to the model. Feature dimensions: $L$ is the number of tokens in the scaffold token set (max 512 during training), $M$ is the number of tokens in the motif token set (max 128 during training) and $A = 37$ is the number of atom types per token.

| Name | Symbol | Shape | Description |
| --- | --- | --- | --- |
| **Scaffold token features** |  |  |  |
| *Sequence token type* | **s** | $[L]$ | Token type index from $\{1, \dots, 64\}$. Values 1–21 for the 20 amino acids + one unknown amino acid; values 22–64 for the 43 ligand atom types. The mask token M in diffusion is not an available type here, instead masking is used to set related features to zero. |
| *Residue index* | **r** | $[L]$ | Index of residue within chain for protein tokens (1-indexed), or atom index within ligand for ligand tokens. |
| *Chain index* | **c** | $[L]$ | Unique integer for each distinct chain. |
| *Ligand mask* | **m<sup>l</sup>** | $[L]$ | Binary mask indicating whether the token is a ligand atom. |
| *Sequence mask* | **m<sup>s</sup>** | $[L]$ | Binary mask indicating valid (non-padding) token positions. |
| *Atom coordinates* | **x** | $[L, A, 3]$ | All-atom coordinates in Å. For protein tokens, up to 37 atoms per residue (padded with virtual atoms with positions at $C_\alpha$). For ligand tokens, only the first atom types is used. |
| *Atom mask* | **m<sup>x</sup>** | $[L, A]$ | Binary mask indicating which atom types are present. |
| *Ligand charge* | **g<sub>c</sub>** | $[L]$ | Formal charge for ligand tokens. Zero for protein tokens. |
| *Ligand hybridisation* | **g<sub>h</sub>** | $[L]$ | Hybridisation state for ligand tokens, i.e. sp, sp<sup>2</sup>, sp<sup>3</sup>, etc. , with a total of 8 categories. Is transformed into one-hot features prior to use, with all zeros for protein tokens. |
| *Indexed motif atom coordinates* | **x<sup>motif</sup>** | $[L, A, 3]$ | Atom coordinates of the indexed motif residues / atoms. |
| *Indexed motif atom mask* | **m<sup>x,motif</sup>** | $[L, A]$ | Binary mask indicating which atoms are provided as conditioning. |
| *Indexed motif sequence token type* | **s<sup>motif</sup>** | $[L]$ | Token type for each indexed motif position. |
| *Indexed motif sequence token mask* | **m<sup>s,motif</sup>** | $[L]$ | Binary mask indicating which tokens have their sequence type fixed. |
| *Self-conditioning atom coordinates* | **x<sup>self</sup>** | $[L, A, 3]$ | Self-conditioning atom coordinates: the denoised coordinate prediction from the previous forward pass (see §4.6). |
| **Motif token features** |  |  |  |
| *Unindexed motif atom coordinates* | **x<sub>unidx</sub><sup>motif</sup>** | $[M, A, 3]$ | Atom coordinates of the motif residues/atoms. |
| *Unindexed motif atom mask* | **m<sub>unidx</sub><sup>x,motif</sup>** | $[M, A]$ | Atom mask for the motif, restricted to fixed atoms. |
| *Unindexed motif sequence token type* | **s<sub>unidx</sub><sup>motif</sup>** | $[M]$ | Token type for each motif position. |
| *Unindexed motif sequence token mask* | **m<sub>unidx</sub><sup>s,motif</sup>** | $[M]$ | Mask indicating which motif positions have a known token type. |
| *Unindexed motif residue index* | **r<sub>unidx</sub>** | $[M]$ | Original residue index of each motif position (1-indexed). |
| *Unindexed motif chain index* | **c<sub>unidx</sub>** | $[M]$ | Original chain index of each motif position. |

### Transformer Block

**Figure S33. Transformer Block.** $n_{\text{block}}$ such blocks are stacked to form the full transformer. Each block applies self-attention independently to scaffold ($\mathbf{q}$) and motif ($\mathbf{k}$) token embeddings with pair bias ($\mathbf{b}_{qq}$, $\mathbf{b}_{kk}$), then cross-attention from scaffold to motif, and finally a SwiGLU transition. The output representation is for the scaffold token set, i.e. the first dimension has size $L$. All sub-layers have residual connections. Single conditioning ($\mathbf{y}_q$, $\mathbf{y}_k$) modulates normalisation via AdaLN.

### Self-Attention

**Figure S34. Self-Attention Detail.** Input $x$ is modulated by Adaptive LayerNorm conditioned on single representation $y$. Queries use biased linear projection; keys and values use bias-free projections. Scaled dot-product attention incorporates additive pair bias $b$. The output is element-wise gated and projected through a zero-initialised linear layer.

### Cross-Attention

**Figure S35. Cross-Attention Detail.** Queries come from the scaffold ($q$) and keys/values from the motif ($k$). Both are modulated by Adaptive LayerNorm. Unlike self-attention, cross-attention has no pair bias. The output is gated and projected through a zero-initialised linear layer.

#### 4.6.2 Denoiser Network

The denoiser network is detailed in Algorithm 3 with a pictorial outline shown in Figure S32. Given the input features, the conditioning embedding (Algorithm 4) constructs three representations: a per-token single representation $\mathbf{y} \in \mathbb{R}^{L \times C_y}$ encoding residue identity and local geometry, a pairwise representation $\mathbf{z} \in \mathbb{R}^{L \times L \times C_z}$ encoding relative positions and inter-residue distances, and a noise embedding $\epsilon$ encoding the current diffusion time steps. Both single and pair representations are refined through residual MLPs with LayerNorm and GELU activations.

After converting the pair embeddings into pair attention biases, these representations are passed to a transformer stack (Figure S33) that processes the single representation using multi-head self-attention (Figure S34) with pair bias and SwiGLU transition layers. For the unindexed motif, a separate embedding (Algorithm 5) computes motif-specific single and pairwise representations, and the transformer additionally applies cross-attention (Figure S35) from scaffold tokens to motif tokens. The conditioning representations ($\mathbf{y}$ and $\mathbf{z}$) modulate the transformer through Adaptive Layer Normalisation and attention logit biasing, respectively, following the adaLN-Zero approach of [39].

The processed activations are linearly projected to produce denoised sequence logits $\hat{\mathbf{p}}_0 \in \mathbb{R}^{L \times K}$ and a structure update $\mathbf{f}_\theta$, which is combined with the noisy input via a skip connection to yield the denoised coordinates $\hat{\mathbf{x}}_0$. In contrast to the comparable skip connection used in EDM, the skip weighting is learned rather than computed as a function of the noise level.

**Algorithm 3** Denoiser network. Predicts denoised coordinates and sequence token types from noisy inputs.

```
1: def $D_\theta(\{\mathbf{x}_t\}, \{\mathbf{s}_t\}, t^x, t^s, \{\mathbf{x}^{\text{self}}\}, \{\mathbf{f}^*\})$:
2: $\sigma_t \leftarrow t^x$, $c_{\text{in}} \leftarrow 1/\sqrt{\sigma_t^2 + \sigma_{\text{data}}^2}$
3: # Normalise noisy coordinates
4: $\mathbf{x}_t \leftarrow c_{\text{in}} \cdot \mathbf{x}_t$ $\mathbf{x}_t \in \mathbb{R}^{L \times A \times 3}$
5: # Compute conditioning embeddings (processes indexed motif masks and sequence types)
6: $\mathbf{y}, \mathbf{z}, \mathbf{f}_{\text{ld}}, \epsilon \leftarrow \text{CONDITIONINGEMBEDDING}(\{\mathbf{x}_t\}, \{\mathbf{s}_t\}, t^x, t^s, \{\mathbf{x}^{\text{self}}\}, \{\mathbf{f}^*\})$ $\mathbf{y} \in \mathbb{R}^{L \times C_y}$, $\mathbf{z} \in \mathbb{R}^{L \times L \times C_z}$
7: # Prepare input activations
8: $\mathbf{h}_{\text{in}} \leftarrow [\text{flatten}(\mathbf{x}_t); \text{flatten}(\mathbf{x}^{\text{self}}); \text{flatten}(\mathbf{x}^{\text{motif}}/\sigma_{\text{data}}); \mathbf{f}_{\text{ld}}]$
9: $\mathbf{h} \leftarrow \text{LINEAR}(\mathbf{h}_{\text{in}}) + \text{LINEARZEROINIT}(\text{LAYERNORM}(\epsilon))$ $\mathbf{h} \in \mathbb{R}^{L \times C}$
10: # Compute unindexed motif embeddings and key activations
11: $\mathbf{y}_{\text{unidx}}, \mathbf{z}_{\text{unidx}}, \mathbf{f}_{\text{ld,unidx}} \leftarrow \text{UNINDEXEDMOTIFCONDITIONINGEMBEDDING}(\mathbf{x}_{\text{unidx}}^{\text{motif}}, \mathbf{s}_{\text{unidx}}^{\text{motif}}, \{\mathbf{f}^*\})$
12: $\mathbf{k} \leftarrow \text{LINEAR}([\text{flatten}(\mathbf{x}_{\text{unidx}}^{\text{motif}}/\sigma_{\text{data}}); \mathbf{f}_{\text{ld,unidx}}])$ $\mathbf{k} \in \mathbb{R}^{M \times C}$
13: $\mathbf{m}_{\text{unidx}}^{\text{mi}} \leftarrow \text{ANY}(\mathbf{m}_{\text{unidx}}^{\text{x,motif}}, \text{AXIS} = \text{atom type})$ $\mathbf{m}_{\text{unidx}}^{\text{mi}} \in \{0, 1\}^M$
14: # Project pairwise features to attention logits
15: $\mathbf{b} \leftarrow \text{PAIRBIAS}(\mathbf{y}, \mathbf{z})$ $\mathbf{b} \in \mathbb{R}^{H \times L \times L}$
16: $\mathbf{b}_{\text{unidx}} \leftarrow \text{PAIRBIAS}(\mathbf{y}_{\text{unidx}}, \mathbf{z}_{\text{unidx}})$ $\mathbf{b}_{\text{unidx}} \in \mathbb{R}^{H \times M \times M}$
17: # Apply transformer blocks with cross-attention
18: for $b = 1, \dots, n_{\text{block}}$ do
19:   $\mathbf{h} \leftarrow \text{TRANSFORMERBLOCKWITHCROSSATTENTION}(\mathbf{h}, \mathbf{k}, \mathbf{m}^s, \mathbf{m}_{\text{unidx}}^{\text{mi}}, \mathbf{y}, \mathbf{y}_{\text{unidx}}, \mathbf{b}, \mathbf{b}_{\text{unidx}})$
20: end for
21: $\mathbf{h} \leftarrow \text{LAYERNORM}(\mathbf{h})$
22: # Predict denoised sequence
23: $\hat{\mathbf{p}}_0 \leftarrow \text{LINEAR}(\mathbf{h})$ $\hat{\mathbf{p}}_0 \in \mathbb{R}^{L \times K}$ (logits)
24: # Predict motif index mask
25: $\hat{\mathbf{m}}^{\text{mi}} \leftarrow \text{LINEAR}(\mathbf{h})$ $\hat{\mathbf{m}}^{\text{mi}} \in \mathbb{R}^L$ (binary logits, log odds ratio)
26: # Predict denoised coordinates via learned skip connection
27: $\mathbf{f}_\theta \leftarrow \text{LINEARZEROINIT}(\mathbf{h})$ $\mathbf{f}_\theta \in \mathbb{R}^{L \times A \times 3}$
28: $c_{\text{skip}} \leftarrow \text{sigmoid}(\text{LINEAR}(\mathbf{h}))$ $c_{\text{skip}} \in \mathbb{R}^{L \times A \times 1}$
29: $\hat{\mathbf{x}}_0 \leftarrow \sigma_{\text{data}} \cdot (c_{\text{skip}} \cdot \mathbf{x}_t + (1 - c_{\text{skip}}) \cdot \mathbf{f}_\theta)$ $\hat{\mathbf{x}}_0 \in \mathbb{R}^{L \times A \times 3}$
30: # Predict uncertainty parameters for dynamic loss weighting
31: $\beta^x \leftarrow \exp(\text{LINEAR}(\epsilon))$ $\beta^x > 0$
32: $\beta^s \leftarrow \exp(\text{LINEAR}(\epsilon))$ $\beta^s > 0$
33: return $\hat{\mathbf{x}}_0, \hat{\mathbf{p}}_0, \hat{\mathbf{m}}^{\text{mi}}, \beta^x, \beta^s$
```

#### 4.6.3 Conditioning Embeddings

The conditioning embedding (Algorithm 4) constructs the single and pairwise representations that condition the transformer stack. For the pairwise representation, we concatenate relative position encodings, a distogram of the noisy coordinates, and a distogram of the self-conditioning coordinates, then project and refine through two residual MLP blocks. For the single representation, we concatenate atom masks, one-hot amino acid types (including the mask state and motif conditioning types masked by $\mathbf{m}^{\text{s,motif}}$), the fixed atom mask $\mathbf{m}^{\text{x,motif}}$, flattened centered coordinates from both the noisy and self-conditioning structures, and ligand features (charge, hybridisation). A sinusoidal embedding of the structure noise level $\sigma/\sigma_{\text{data}}$ and the sequence time step $t^s$ is added to the single representation. Both representations are refined with residual MLPs using LayerNorm and GELU activations.

For the unindexed motif, a separate embedding (Algorithm 5) computes motif-specific single and pairwise representations using the motif atom coordinates and sequence token types, with a neighbour encoding and distogram for the pairwise features.

**Algorithm 4** Conditioning embedding. Computes single and pairwise conditioning representations from input features.

```
1: def CONDITIONINGEMBEDDING($\{\mathbf{x}_t\}, \{\mathbf{s}_t\}, t^x, t^s, \{\mathbf{x}^{\text{self}}\}, \{\mathbf{f}^*\}$):
2:   # Compute noise level / time step embeddings
3:   $\sigma_t \leftarrow t^x$
4:   $\epsilon_{\text{struct}} \leftarrow \text{RANDOMFOURIEREMBED}(\frac{1}{4} \ln(\sigma_t/\sigma_{\text{data}}))$ $\epsilon_{\text{struct}} \in \mathbb{R}^{C_\epsilon}$
5:   $\epsilon_{\text{seq}} \leftarrow \text{SINUSOIDALEMBED}(t^s)$ $\epsilon_{\text{seq}} \in \mathbb{R}^{C_\epsilon}$
6:   $\epsilon \leftarrow [\epsilon_{\text{struct}}; \epsilon_{\text{seq}}]$
7:   # Compute pairwise features
8:   $\mathbf{z}_{\text{rel}} \leftarrow \text{RELATIVEPOSITIONENCODING}(\mathbf{r}, \mathbf{c})$ $\mathbf{z}_{\text{rel}} \in \mathbb{R}^{L \times L \times C_{\text{rel}}}$
9:   $\mathbf{z}_{\text{dgram}} \leftarrow \text{DISTOGRAM}(\{\mathbf{x}_t\})$ $\mathbf{z}_{\text{dgram}} \in \mathbb{R}^{L \times L \times n_{\text{bins}}}$
10:  $\mathbf{z}_{\text{self}} \leftarrow \text{DISTOGRAM}(\{\mathbf{x}^{\text{self}}/\sigma_{\text{data}}\})$ $\mathbf{z}_{\text{self}} \in \mathbb{R}^{L \times L \times n_{\text{bins}}}$
11:  $\mathbf{m}^{\text{ss}} = \mathbf{m}^{\text{s}} \otimes \mathbf{m}^{\text{s}} \otimes \mathbf{1}$ $\mathbf{m}^{\text{ss}} \in [0, 1]^{L \times L \times 1}$
12:  $\mathbf{z}_{\text{pair}} \leftarrow [\mathbf{z}_{\text{rel}}; \mathbf{z}_{\text{dgram}}; \mathbf{z}_{\text{self}}] \odot \mathbf{m}^{\text{ss}}$
13:  # Project pairwise features with residual MLP
14:  $\mathbf{z} \leftarrow \text{LINEAR}(\text{LAYERNORM}(\mathbf{z}_{\text{pair}}))$ $\mathbf{z} \in \mathbb{R}^{L \times L \times C_z}$
15:  for $i = 1, 2$ do
16:    $\mathbf{z} \leftarrow \mathbf{z} + \text{LINEARZEROINIT}(\text{GELU}(\text{LINEAR}(\text{LAYERNORM}(\mathbf{z}))))$
17:  end for
18:  # Compute single (per-residue) features
19:  $\mathbf{f}_{\text{ld}} \leftarrow [(\mathbf{m}^{\text{x}})^{\otimes}; \text{ONEHOT}(\mathbf{s}_t); \mathbf{s}^{\text{motif}}; \mathbf{m}^{\text{x,motif}}; \text{CENTERFLATTEN}(\{\mathbf{x}_t\}); \text{CENTERFLATTEN}(\{\mathbf{x}^{\text{self}}\});$
20:    $\mathbf{g}_{\text{c}}^{\otimes}; \text{ONEHOT}(\mathbf{g}_{\text{h}})]$ $\mathbf{f}_{\text{ld}} \in \mathbb{R}^{L \times \cdot}$
21:  $\mathbf{f}_{\text{ld}} \leftarrow \mathbf{f}_{\text{ld}} \odot (\mathbf{m}^{\text{s}})^{\otimes}$
22:  $\mathbf{y} \leftarrow \text{LINEAR}(\text{LAYERNORM}(\mathbf{f}_{\text{ld}}))$ $\mathbf{y} \in \mathbb{R}^{L \times C_y}$
23:  # Add noise embedding to single representation
24:  $\mathbf{y} \leftarrow \mathbf{y} + \text{LINEAR}(\text{LAYERNORM}(\epsilon))$
25:  # Refine single representation with residual MLP
26:  for $i = 1, 2$ do
27:    $\mathbf{y} \leftarrow \mathbf{y} + \text{LINEARZEROINIT}(\text{GELU}(\text{LINEAR}(\text{LAYERNORM}(\mathbf{y}))))$
28:  end for
29:  return $\mathbf{y}, \mathbf{z}, \mathbf{f}_{\text{ld}}, \epsilon$
```

**Algorithm 5** Unindexed motif conditioning embedding. Computes single and pairwise conditioning for unindexed motif residues. Here, the inputs are all unindexed motif conditioning, i.e. $\mathbf{x} := \mathbf{x}_{\text{unidx}}^{\text{motif}}$, $\mathbf{m}^x := \mathbf{m}_{\text{unidx}}^{x, \text{motif}}$, and $\mathbf{s} := \mathbf{s}_{\text{unidx}}^{\text{motif}}$.

```
1: def UNINDEXEDMOTIFCONDITIONINGEMBEDDING($\mathbf{x}, \mathbf{m}^x, \mathbf{s}$):
2:   Normalise input atom coordinates
3:   $\mathbf{x} \leftarrow \mathbf{x} / \sigma_{\text{data}}$
4:   # Compute single (per-residue) features for motif
5:   $\mathbf{m}^a \leftarrow \text{ANY}(\mathbf{m}^x, -1)$ $\mathbf{m} \in \{0, 1\}^{M \times A}$
6:   $\mathbf{f}_{\text{ld}} \leftarrow [\mathbf{m}^a; \text{ONEHOT}(\mathbf{s}); \text{CENTERFLATTEN}(\mathbf{x})]$
7:   $\mathbf{y} \leftarrow \text{LINEAR}(\text{LAYERNORM}(\mathbf{f}_{\text{ld}}))$ $\mathbf{y} \in \mathbb{R}^{M \times C_y}$
8:   # Refine single representation with residual MLP
9:   for $i = 1, 2$ do
10:    $\mathbf{y} \leftarrow \mathbf{y} + \text{LINEARZEROINIT}(\text{GELU}(\text{LINEAR}(\text{LAYERNORM}(\mathbf{y}))))$
11:   end for
12:   # Compute pairwise features for motif
13:   $\mathbf{z}_{\text{neighbour}} \leftarrow \text{NEIGHBOURENCODING}(\mathbf{r}_{\text{unidx}}, \mathbf{c}_{\text{unidx}})$ $\mathbf{z}_{\text{neighbour}} \in \mathbb{R}^{M \times M \times 3}$
14:   $\mathbf{z}_{\text{dgram}} \leftarrow \text{DISTOGRAM}(\{\mathbf{x}\})$
15:   $\mathbf{z}_{\text{pair}} \leftarrow [\mathbf{z}_{\text{neighbour}}; \mathbf{z}_{\text{dgram}}]$
16:   # Project pairwise features with residual MLP
17:   $\mathbf{z} \leftarrow \text{LINEAR}(\text{LAYERNORM}(\mathbf{z}_{\text{pair}}))$ $\mathbf{z} \in \mathbb{R}^{M \times M \times C_z}$
18:   for $i = 1, 2$ do
19:    $\mathbf{z} \leftarrow \mathbf{z} + \text{LINEARZEROINIT}(\text{GELU}(\text{LINEAR}(\text{LAYERNORM}(\mathbf{z}))))$
20:   end for
21:   return $\mathbf{y}, \mathbf{z}, \mathbf{f}_{\text{ld}}$
```

##### 4.6.4 Transformer Stack

The transformer stack processes the token-level activations through $n_{\text{block}}$ blocks (Figure S33, Algorithm 6). Each block applies multi-head self-attention with pair bias followed by a SwiGLU transition, both with residual connections. Each block applies self-attention independently to scaffold and motif representations, followed by cross-attention from the scaffold queries to the motif keys, and a feedforward transition. This allows information from the motif to flow into the scaffold representation without requiring explicit scaffold index placement of the motif.

**Algorithm 6** Transformer block with cross-attention. Computes updates via self-attention, cross-attention, and transition layers with residual connections. The queries correspond to scaffold tokens and the keys correspond to unindexed motif tokens.

```
1: def TRANSFORMERBLOCKWITHCROSSATTENTION($\mathbf{q}, \mathbf{k}, \mathbf{m}_q, \mathbf{m}_k, \mathbf{y}_q, \mathbf{y}_k, \mathbf{b}_{qq}, \mathbf{b}_{kk}$):
2:   # Self-attention on queries
3:   $\mathbf{q}' \leftarrow \mathbf{q} + \text{SELFATTENTION}(\mathbf{q}, \mathbf{m}_q, \mathbf{y}_q, \mathbf{b}_{qq})$
4:   # Self-attention on keys
5:   $\mathbf{k}' \leftarrow \mathbf{k} + \text{SELFATTENTION}(\mathbf{k}, \mathbf{m}_k, \mathbf{y}_k, \mathbf{b}_{kk})$
6:   # Cross-attention from queries to keys
7:   $\mathbf{h} \leftarrow \mathbf{q}' + \text{CROSSATTENTION}(\mathbf{q}', \mathbf{k}', \mathbf{m}_q, \mathbf{m}_k, \mathbf{y}_q, \mathbf{y}_k)$
8:   # Feedforward transition
9:   $\mathbf{h}' \leftarrow \mathbf{h} + \text{TRANSITION}(\mathbf{h}, \mathbf{y}_q)$
10:  return $\mathbf{h}'$
```

##### 4.6.5 Attention Mechanisms

The attention layers (Algorithms 7 and 8, Figures S34 and S35) employ multi-head scaled dot-product attention with three key modifications. First, input activations are modulated by Adaptive LayerNorm conditioned on the single representation, providing per-residue conditioning. Second, the pairwise representation is linearly projected to produce per-head additive biases on the attention logits, allowing the network to directly incorporate structural and relational information into the attention pattern. Third, a gating mechanism multiplies the attention output elementwise by $\text{sigmoid}(\text{LINEARZEROINIT}(\mathbf{x}))$, initialised to pass zero at the start of training. The output projection is also zero-initialised following the adaLN-Zero formulation, ensuring that each block acts as an identity at initialisation.

**Algorithm 7** Multi-head self-attention. Attention is within scaffold tokens (as notated, with token dimension size $L$) or within unindexed motif tokens (replace $L$ by $M$).

```
1: def SELFATTENTION($\mathbf{x}, \mathbf{m}, \mathbf{y}, \mathbf{b}$):
2:   # Adaptive layer normalisation
3:   $\mathbf{x}' \leftarrow \text{ADAPTIVELAYERNORM}(\mathbf{x}, \mathbf{y})$
4:   # Project to queries, keys, and values
5:   $\mathbf{Q} \leftarrow \text{LINEAR}(\mathbf{x}')$
6:   $\mathbf{K} \leftarrow \text{LINEARNOBIAS}(\mathbf{x}')$
7:   $\mathbf{V} \leftarrow \text{LINEARNOBIAS}(\mathbf{x}')$
8:   # Scaled dot-product attention with bias and masking, repeated for each head.
9:   $\mathbf{A} \leftarrow \text{softmax} \left( \left( \frac{\mathbf{Q}\mathbf{K}^\top}{\sqrt{d_k}} + \mathbf{b} \right) \odot \mathbf{m}^a \right) \mathbf{V}$
10:  # Output gating
11:  $\mathbf{g} \leftarrow \text{sigmoid}(\text{LINEARZEROINIT}(\mathbf{x}))$
12:  $\mathbf{A} \leftarrow \mathbf{g} \odot \mathbf{A}$
13:  # Project back to hidden dimension
14:  $\mathbf{o} \leftarrow \text{DROPOUT}_{0.3}(\text{LINEARZEROINIT}(\mathbf{A}))$
15:  return $\mathbf{o}$
```

$\mathbf{Q} \in \mathbb{R}^{L \times H \times d_k}$
$\mathbf{K} \in \mathbb{R}^{L \times H \times d_k}$
$\mathbf{V} \in \mathbb{R}^{L \times H \times d_v}$
$\mathbf{A} \in \mathbb{R}^{L \times H \times d_v}$
$\mathbf{g} \in \mathbb{R}^{L \times H \times d_v}$
$\mathbf{o} \in \mathbb{R}^{L \times C}$

**Algorithm 8** Multi-head cross-attention from queries to keys. The queries correspond to scaffold tokens and the keys correspond to unindexed motif tokens.

```
1: def CROSSATTENTION($\mathbf{q}, \mathbf{k}, \mathbf{m}_q, \mathbf{m}_k, \mathbf{y}_q, \mathbf{y}_k$):
2:   # Adaptive layer normalisation for both sequences
3:   $\mathbf{q}' \leftarrow \text{ADAPTIVELAYERNORM}(\mathbf{q}, \mathbf{y}_q)$
4:   $\mathbf{k}' \leftarrow \text{ADAPTIVELAYERNORM}(\mathbf{k}, \mathbf{y}_k)$
5:   # Project to queries, keys, and values
6:   $\mathbf{Q} \leftarrow \text{LINEAR}(\mathbf{q}')$
7:   $\mathbf{K} \leftarrow \text{LINEARNOBIAS}(\mathbf{k}')$
8:   $\mathbf{V} \leftarrow \text{LINEARNOBIAS}(\mathbf{k}')$
9:   # Scaled dot-product attention with masking
10:  $\mathbf{A} \leftarrow \text{softmax} \left( \frac{\mathbf{Q}\mathbf{K}^\top}{\sqrt{d_k}} \right) \mathbf{V}$
11:  # Output gating
12:  $\mathbf{g} \leftarrow \text{sigmoid}(\text{LINEARZEROINIT}(\mathbf{q}'))$
13:  $\mathbf{A} \leftarrow \mathbf{g} \odot \mathbf{A}$
14:  # Project back and apply dropout
15:  $\mathbf{o} \leftarrow \text{DROPOUT}_{0.3}(\text{LINEARZEROINIT}(\mathbf{A}))$
16:  return $\mathbf{o}$
```

$\mathbf{Q} \in \mathbb{R}^{L \times H \times d_k}$
$\mathbf{K} \in \mathbb{R}^{M \times H \times d_k}$
$\mathbf{V} \in \mathbb{R}^{M \times H \times d_v}$
$\mathbf{A} \in \mathbb{R}^{L \times H \times d_v}$
$\mathbf{g} \in \mathbb{R}^{L \times H \times d_v}$
$\mathbf{o} \in \mathbb{R}^{L \times C}$

##### 4.6.6 Supporting Modules

Pair bias creation (Algorithm 9) follows the approach used in AlphaFold 3, applying an adaptive layer norm followed by a projection. The transformer transition layer (Algorithm 10) is a SwiGLU-based feedforward block that expands the hidden dimension, applies $\text{SiLU}(\mathbf{a}) \odot \mathbf{b}$ gating, and projects back, following [40]. All transition blocks in the model use SwiGLU rather than ReLU. The Adaptive LayerNorm (Algorithm 11) conditions activations on the single representation by learning per-residue scale and bias parameters, applying them as $\text{sigmoid}(\gamma) \odot \text{LAYERNORM}(\mathbf{x}) + \beta$. When no conditioning signal is provided (e.g. for pairwise representations), standard LayerNorm is used instead. The relative position encoding (Algorithm 12) computes pairwise features from residue indices and chain identifiers: within-chain residue offsets are clipped to $[-d_{\max}, d_{\max}]$ (with $d_{\max} = 64$) and one-hot encoded, while cross-chain pairs are assigned to a separate bin, yielding $2d_{\max} + 2 = 130$ features per pair. The distogram (Algorithm 13) computes a one-hot pairwise distance feature: for each token, the mean position of its valid atoms is computed, pairwise distances are calculated, and each distance is binned into $n_{\text{bins}} = 15$ bins with edges linearly spaced from 2 Å to 15 Å, yielding a 15-dimensional feature per token pair. The noise level is embedded via two different encodings: the structure noise level uses a random Fourier embedding (Algorithm 14), which projects $\frac{1}{4} \ln \sigma$ through fixed random frequencies, analogous to [41] and the Fourier embedding in AlphaFold 3 [15]; the sequence noise level uses a sinusoidal positional embedding (Algorithm 15) with geometric frequency spacing, following [42]. Both produce $C_\epsilon = 256$-dimensional features that are concatenated into a 512-dimensional noise embedding.

**Algorithm 9** Pair bias creation. Notated with token dimension size $L$, but uses token dimension size $M$ for unindexed motif pair bias.

```
1: def PAIRBIAS($\mathbf{y}$, $\mathbf{z}$):
2:   # Compute attention logits from pairwise conditioning
3:   $\mathbf{z}' \leftarrow \text{ADAPTIVELAYERNORM}(\mathbf{z}, \mathbf{y})$ $\mathbf{z} \in \mathbb{R}^{L \times L \times C_z}$
4:   # Project pairwise features to attention logits
5:   $\mathbf{b} \leftarrow \text{ROLLAXISRIGHT}(\text{LINEAR}(\mathbf{z}'))$ $\mathbf{b} \in \mathbb{R}^{H \times L \times L}$
6:   return $\mathbf{b}$
```

**Algorithm 10** SwiGLU transition layer with adaptive layer normalisation. Notated with token dimension size $L$, but uses token dimension size $M$ for unindexed motif self attention.

```
1: def TRANSITION($\mathbf{h}$, $\mathbf{y}$):
2:   # Adaptive layer normalisation
3:   $\mathbf{h}' \leftarrow \text{ADAPTIVELAYERNORM}(\mathbf{h}, \mathbf{y})$
4:   # Expand to intermediate dimension $C_{\text{int}} = n_f \cdot C$ ($n_f = 2$) and split for gating
5:   $[\mathbf{a}, \mathbf{b}] \leftarrow \text{split}(\text{LINEAR}(\mathbf{h}'))$ $\mathbf{a}, \mathbf{b} \in \mathbb{R}^{L \times C_{\text{int}}}$
6:   # SwiGLU activation
7:   $\mathbf{c} \leftarrow \text{SiLU}(\mathbf{a}) \odot \mathbf{b}$
8:   # Project back and apply dropout
9:   $\mathbf{o} \leftarrow \text{DROPOUT}_{0.3}(\text{LINEARZEROINIT}(\mathbf{c}))$ $\mathbf{o} \in \mathbb{R}^{L \times C}$
10: return $\mathbf{o}$
```

**Algorithm 11** Adaptive layer normalisation conditioned on single representation. Notated with token dimension size $L$, but uses token dimension size $M$ for unindexed motif related layers.

```
1: def ADAPTIVELAYERNORM($\mathbf{h}$, $\mathbf{y}$):
2:   # Normalise input without learnable parameters
3:   $\mathbf{h}' \leftarrow \text{LAYERNORMNOPARAMS}(\mathbf{h})$
4:   # Normalise conditioning and project to scale and bias
5:   $\mathbf{y} \leftarrow \text{LAYERNORM}(\mathbf{y})$
6:   $\boldsymbol{\gamma} \leftarrow \text{LINEARZEROINIT}(\mathbf{y})$ $\boldsymbol{\gamma} \in \mathbb{R}^{L \times C}$
7:   $\boldsymbol{\beta} \leftarrow \text{LINEARZEROINIT}(\mathbf{y})$ $\boldsymbol{\beta} \in \mathbb{R}^{L \times C}$
8:   # Apply adaptive modulation
9:   $\mathbf{h}'' \leftarrow \text{sigmoid}(\boldsymbol{\gamma}) \odot \mathbf{h}' + \boldsymbol{\beta}$
10: return $\mathbf{h}''$
```

**Algorithm 12** Relative position encoding. Computes pairwise relative position features from residue indices and chain identifiers. Only used for scaffold token embeddings, not unindexed motif.

```
1: def RELATIVEPOSITIONENCODING($\mathbf{r}$, $\mathbf{c}$):
2:   # Determine same-chain mask
3:   $\mathbf{M}_{\text{same}} \leftarrow \mathbf{1}[\mathbf{c}_i = \mathbf{c}_j]$ $\mathbf{M}_{\text{same}} \in \{0, 1\}^{L \times L}$
4:   # Compute pairwise residue index offset
5:   $\Delta_{ij} \leftarrow \mathbf{r}_i - \mathbf{r}_j$ $\Delta \in \mathbb{Z}^{L \times L}$
6:   # Clip offset to $[0, 2d_{\text{max}}]$
7:   $\bar{\Delta}_{ij} \leftarrow \text{clip}(\Delta_{ij} + d_{\text{max}}, 0, 2d_{\text{max}})$
8:   # Assign cross-chain pairs to a separate bin
9:   $\hat{\Delta}_{ij} \leftarrow \mathbf{M}_{\text{same}, ij} \cdot \bar{\Delta}_{ij} + (1 - \mathbf{M}_{\text{same}, ij}) \cdot (2d_{\text{max}} + 1)$
10:  # One-hot encode into $2d_{\text{max}} + 2$ bins
11:  $\mathbf{z}_{\text{rel}} \leftarrow \text{ONEHOT}(\hat{\Delta}, 2d_{\text{max}} + 2)$ $\mathbf{z}_{\text{rel}} \in \mathbb{R}^{L \times L \times (2d_{\text{max}} + 2)}$
12:  return $\mathbf{z}_{\text{rel}}$
```

**Algorithm 13** Distogram. Computes one-hot pairwise distance features from atom coordinates and masks. Notated with token dimension size $L$, but uses token dimension size $M$ for unindexed motif distogram.

```
1: def DISTOGRAM($\mathbf{x}$, $\mathbf{m}$):
2:   # Compute per-token centre (masked mean over atoms)
3:   $\bar{\mathbf{x}}_i \leftarrow \sum_a \mathbf{m}_{ia} \mathbf{x}_{ia} / \sum_a \mathbf{m}_{ia}$ $\bar{\mathbf{x}} \in \mathbb{R}^{L \times 3}$
4:   # Sequence mask: true for tokens with at least one valid atom
5:   $\mathbf{v}_i \leftarrow \mathbf{1}[\sum_a \mathbf{m}_{ia} > 0]$ $\mathbf{v} \in \{0, 1\}^L$
6:   # Pairwise squared distance
7:   $d_{ij}^2 \leftarrow \|\bar{\mathbf{x}}_i - \bar{\mathbf{x}}_j\|^2$ $d^2 \in \mathbb{R}^{L \times L}$
8:   # Bin edges: $n_{\text{bins}} = 15$, linearly spaced from $b_{\text{min}} = 2 \text{ \AA}$ to $b_{\text{max}} = 15 \text{ \AA}$, then squared
9:   $\ell_k \leftarrow \left( b_{\text{min}} + k \cdot \frac{b_{\text{max}} - b_{\text{min}}}{n_{\text{bins}} - 1} \right)^2, \quad u_k \leftarrow \begin{cases} \ell_{k+1} & k < n_{\text{bins}} - 1 \\ \infty & k = n_{\text{bins}} - 1 \end{cases}$
10:  # One-hot binning
11:  $\mathbf{z}_{ij,k} \leftarrow \mathbf{1}[\ell_k \leq d_{ij}^2 < u_k]$ $\mathbf{z} \in \{0, 1\}^{L \times L \times n_{\text{bins}}}$
12:  # Mask invalid tokens
13:  $\mathbf{z}_{ij} \leftarrow \mathbf{v}_i \cdot \mathbf{v}_j \cdot \mathbf{z}_{ij}$
14:  return $\mathbf{z}$
```

**Algorithm 14** Random Fourier embedding. Projects a scalar input through fixed random frequencies.

```
1: def RANDOMFOURIEREMBED($x$):
2:   # Fixed random projection (deterministic seed)
3:   $\mathbf{w} \sim \mathcal{N}(0, \mathbf{I}_D), \quad \mathbf{b} \sim \text{Uniform}(0, 1)^D$ (fixed at initialisation)
4:   $\mathbf{e} \leftarrow \cos(2\pi(x \cdot \mathbf{w} + \mathbf{b}))$ $\mathbf{e} \in \mathbb{R}^D$
5:   return $\mathbf{e}$
```

**Algorithm 15** Sinusoidal positional embedding. Encodes a scalar using geometric frequency spacing.

```
1: def SINUSOIDALEMBED($x$):
2:   $\omega_k \leftarrow \exp(-k \cdot \ln(10000) / (D/2 - 1)), \quad k = 0, \dots, D/2 - 1$
3:   $\mathbf{e} \leftarrow [\cos(1000x \cdot \omega_0), \dots, \cos(1000x \cdot \omega_{D/2-1}), \sin(1000x \cdot \omega_0), \dots, \sin(1000x \cdot \omega_{D/2-1})]$ $\mathbf{e} \in \mathbb{R}^D$
4:   return $\mathbf{e}$
```

**Algorithm 16** Neighbour encoding for unindexed motif. Computes a one-hot pairwise feature indicating immediate forwards or backwards sequence adjacency within the same chain.

```
1: def NEIGHBOURENCODING($\mathbf{r}$, $\mathbf{c}$):
2:   # Determine same-chain mask
3:   $\mathbf{M}_{\text{same}} \leftarrow \mathbf{1}[\mathbf{c}_i = \mathbf{c}_j]$ $\mathbf{M}_{\text{same}} \in \{0, 1\}^{M \times M}$
4:   # Compute pairwise residue index offset
5:   $\Delta_{ij} \leftarrow \mathbf{r}_i - \mathbf{r}_j$ $\Delta \in \mathbb{Z}^{M \times M}$
6:   # One-hot encode offsets $\{-1, 0, +1\}$ into 3 bins; all other offsets map to 0
7:   $\mathbf{z}_{ij} \leftarrow \text{ONEHOT}(\Delta_{ij} + 1, 3) \cdot \mathbf{M}_{\text{same}, ij}$ $\mathbf{z} \in \mathbb{R}^{M \times M \times 3}$
8:   return $\mathbf{z}$
```

### 5 Structure Prediction for Benchmarking and Filtering

We used AF3 or a fine-tuned variant, AF3-LeavingAtom (“AF3-LA”), for structure prediction based on single-sequence inputs (“without MSA”). The derived metrics were used for calculating *in silico* benchmark pass rates and to filter designs for experimental testing.

In some instances we ran structure prediction multiple times with different random seeds to produce an ensemble of predicted structures (“multi-seed” structure prediction). In general, AF3 uses a “trunk” seed to control RDKit conformer generation for obtaining ligand input features and to initialise the diffusion head. For a single trunk seed, five structures are predicted from different initialisations of the diffusion head and the prediction with highest ranking confidence is returned as the final prediction, resulting in one final prediction per trunk seed.

We also made “multi-state” predictions to evaluate active site conformations in different steps of a catalytic cycle. Each state is represented by the choice of a particular ligand and protein–ligand bonds. State and metric aggregations over multi-seed folds are denoted by prefixes and suffixes in the text below. For example, `apo/rmsd/mean` indicates the mean over multiple seeds of the RMSD between prediction and design in a protein-only state.

### 5.1 AlphaFold3 Leaving Atom (AF3-LA)

We noticed that AF3 predictions of ligands covalently bonded to proteins had a high incidence of geometry errors in the region of the inter-chain covalent bond. For example, serine esterase designs with the nucleophilic serine covalently bonded to a tetrahedral intermediate had bond angles of $\sim 180^\circ$ at $O_\gamma$, which is considerably bigger than the expected tetrahedral angle of $109^\circ$.

We hypothesised that AF3 makes these geometric errors because training incorrectly handled leaving atoms for covalently bonded ligands. To address this, we fine-tuned the AF3 weights for an additional 1.6M samples using the same architecture and losses (other than the covalent bond loss, see below) but with modifications to the handling of leaving atoms in the data pipeline. We refer to the modified model as AF3-LeavingAtom (AF3-LA).

• For non-glycan covalently-bound ligands, AF3 did not remove leaving atoms, causing the “ground truth” structure to often contain leaving atoms and protein residues within clash distance. AF3-LA now removes leaving atoms if they have a neighbour that is covalently bonded to another chain.
• For glycan covalently-bound ligands, AF3 did correctly remove $O_1$ leaving atoms during training. However, it also incorrectly removed $O_1$ from non-covalent glycans and did not remove non-$O_1$ leaving atoms from glycans. AF3-LA now removes leaving atoms from covalent glycans and does not remove them from non-covalent glycans.
• AF3 used a single token to represent protein residues with covalent bonds to ligands, leading to the need for a special loss to improve the position of the bond. AF3-LA now uses one token per atom for such residues, similar to RosettaFold-AllAtom [13] and AF3’s representation of post-translationally modified residues. This makes the standard bond conditioning specific to the correct atom in the residue, improving covalent bond geometries.

Note that AF3-LA still incorrectly handles examples containing multiple heavy leaving atoms, but these are a small fraction of the training data. AF3-LA also contains the following minor modifications, which we do not expect to affect prediction accuracy except potentially for large or DNA-containing structures:

• Fine-tuned at a crop size of 640 tokens (smaller than the 768 used for AF3).
• Removed the covalent bond loss (no longer necessary after using one token per atom for bonded residues).
• Removed DNA distillation data (as used in the final stage of AF3 training).
• Trained on covalent bonds that are not symmetric, even if the chains have symmetry.
• Removed the limit on the number of symmetric chains in training structures.
• Set eval selection cutoff at 1,536 tokens rather than 2,048.

AF3-LA had much-improved geometries in the region of the covalent bond between protein and ligand, and was thus used for *in silico* benchmarks and some of the design campaigns below. When running AF3-LA we always ran with three recycles instead of the default ten recycles that AF3 uses.

### 6 AME Benchmark

To evaluate the APN diffusion model *in silico*, we used the recently published Atomic Motif Enzyme (AME) benchmark, a suite of 41 active-site scaffolding problems specifying protein motif tip atoms and ligands [29]. However, we note that, in general, *in silico* benchmarks are highly simplified proxies for real-world design campaigns and consequently, benchmark improvements may not translate directly to experimental success.

Our implementation of the AME benchmark follows the published description [43] and code [29], with minor modifications. For each motif, we sampled 100 backbones, generated eight LigandMPNN sequences per backbone, predicted structures for each of those eight sequences with five different trunk seeds using AF3-LA, and considered a backbone a “pass” if for any of the eight sequences, any of the five predicted structures meet the following criteria:

• `expanded_motif/motif_bb_aligned_allatom_rmsd` $< 1.5$, defined as the RMSD between designed and predicted structures over all atoms of any residue containing at least one motif atom, after aligning on backbone (N, C$_{\alpha}$, C) atoms of those residues.
• `percent_ligand_bb_clashes_15` = 0, defined as the number of ligand atoms within 1.5 Å of any protein backbone atom.

Using AF3-LA in this benchmark gives similar overall results as AF3 (Figure S2E). However, AF3 incorrectly drops glycan O<sub>1</sub> atoms for three motifs, so instead we used AF3-LA to improve problem coverage.

To compare to the current state-of-the-art generative model for *de novo* enzyme design, RFDiffusion 3 (RFD3), we installed RFD3 and LigandMPNN via the RosettaCommons Foundry. We ran RFD3 with default settings (`num_timesteps` 200, `step_scale` 1.5, `noise_level` 0.6) and LigandMPNN with the `ligandmpnn_v_32_010_25.pt` checkpoint at temperature 0.1. We modified the AME benchmark as follows:

• We removed motifs M0058 (four islands) and M0110 (three islands), as RFD3 performs poorly on them due to technical reasons (truncation of residue numbers to three digits in the RFD3 data pipeline).
• We used AF3-LA instead of Chai-1 as the structure prediction model.
• We define a pass as a design for which any of eight LigandMPNN sequences passes filters, whereas RFD3 first selects a fold based on structure prediction confidence and then applies pass criteria.
• We align only on (N, C$_{\alpha}$, C) backbone atoms, while the previously published AME benchmark also includes O.

Our execution of the benchmark leads to very similar pass rates for RFD3 (“RFD3 rerun”) compared to the original paper (“RFD3 paper”) (Figure S2C).

We evaluated the APN diffusion model with both indexed (APN-I) and unindexed (APN-U) motif inputs. For APN-I, we randomly sample the permutation and design segment lengths until a total of 180 residues is reached. We find that the APN diffusion model has comparable or better *in silico* pass rates than RFD3 on the AME benchmark across different numbers of motif islands (Figure S2B) as well as individual motifs (Figure S2A,D). We also check that matching the number of diffusion steps to RFD3 (200) does not change our conclusions (Figure S2F). Therefore, we conclude that APN will be as or more sample-efficient than other similar models in generating candidate designs for experimental testing.

In order to score APN-U on AME, we had to make a modification to the outputs. Unlike RFD3 and APN-I, there is no enforced reproduction of input motif coordinates for APN-U (APN-I and RFD3 do this via replacement mechanisms during diffusion rollouts). APN-U can therefore generate motifs that do not match the input conditioning and does this in a way that tends to reduce RMSD between designed motif and folded motif. Because AME scores use this RMSD rather than the RMSD between input motif and folded motif, APN-U would have an unfair advantage if we used the default scoring setup. Instead, we override the default behavior of APN-U by replacing the final generated motif coordinates with the coordinates from the processed input motif. Additionally, as APN-U is not guaranteed to reproduce all residues of the input motif (e.g. it is possible that the generated design can contain no MET residues even if MET is present in the input motif) we consider a design non-passing if it is missing any residues from the input motif. This procedure can be applied for fair comparison of any similar model that does not enforce reproduction of input motif coordinates.

We hypothesised that if APN-U does not typically reproduce an input motif accurately, then that motif is challenging to scaffold well and other models would show signs of difficulty on the same motif. We found that designs from APN-I and RFD3 tended to have more broken motif intra-residue bonds (defined as $> 0.5$ Å from ideal values) for designs against such motifs, correlating with higher RMSD between input and folded motif in the APN-U (Figure S2H). This suggests that when APN-U deviates from the input motif, it is performing local structure prediction to avoid out-of-distribution bond geometries. Finally we note that the worst metrics occur for motifs with more residue islands, which are expected to be more difficult to scaffold.

### 7 Filtering Metrics

This section defines filtering metrics used in the design campaigns detailed in §8, where threshold values and reaction-specific metrics are given per campaign. Structure metrics are prefixed with something to indicate what structure they relate to: either “sampled” for the diffusion model sample structure (cannot be used for consistency metrics), “apo” for the protein only, “complex” for the protein and ligand combination if only one catalytic state is used or the name of the catalytic state if there are many involved.

### 7.1 Consistency metrics

Unless noted otherwise, these all compare a predicted structure and the structure output of diffusion model.

• `rmsd`: Root mean square deviation over all protein $C_{\alpha}$ atoms.
• `tm_score`: Template Modelling score measuring length-normalised global structural similarity.
• `mean_pocket_bb_aligned_ligand_rmsd`: Mean RMSD of ligand heavy atoms after superimposing the predicted structure onto the reference on backbone atoms within 10 Å of the ligand.
• `mean_pocket_bb_rmsd`: Mean RMSD of pocket backbone atoms as defined above.
• `interface_lddt`: LDDT computed between residues or atoms not on the same chain.
• `fixed_res_bb_aligned_allatom_sc_rmsd`: Backbone-aligned all-atom RMSD for residues with any atom part of a motif.
• `motif_allatom_rmsd`: All-atom RMSD between AF3 prediction and input motif structure.

### 7.2 Confidence metrics

• `protein_plddt_mean`: Predicted Local Distance Difference Test score averaged over the protein, a measure of confidence in the protein prediction.
• `ligand_plddt_mean`: Mean ligand pLDDT.
• `predicted_distance_error`: AF3 predicted distance error over all atoms.
• `chain_pair_pde_mean_sym`: Symmetrised mean predicted distance error between chains.

### 7.3 Structure topology metric

Only ever measured on the diffusion model output structure.

• `topology_string_len`: Number of secondary structure elements (e.g. three for 3-helical bundle).

### 7.4 Sequence metrics

• `seq_len`: Sequence length.
• `max_single_aatype_frac`: Max fraction of any single amino acid type.
• `max_consecutive_identical_aatypes`: Max consecutive identical residues.
• `num_blocklisted_motifs`: Number of subsequences matching anything from the following sequences. WSH PQFEK (Strep-tag), EQKLISEEDL (c-Myc-tag), ENLYFQ[SGAMCH] (TEV protease), LEVLFQGP (3C protease cleavage site), YPYDVPDYA (HA-tag), HHHHH (His-tag), DYKDDDDK (FLAG-tag), GKPIPNPLLGLDST (V5-tag), LVPRGS (Thrombin cleavage).

### 7.5 Multiple-resequencing and multiple-partial-diffusion filters

The multiple-resequencing filter scores a protein backbone. It uses LigandMPNN to obtain a pre-defined number of resequences for that backbone and then applies a set of filters to each sequence. A backbone passes the multiple-resequencing filter if a specific number of its resequences pass this filter setting. The multiple-partial-diffusion filter is similar, but instead of LigandMPNN, partial diffusion is applied to the design to generate a new set of sequences corresponding to similar backbones.

### 8 Design Campaigns

Unless otherwise noted below, all design campaign rounds were carried out by:

• Generating designs using a diffusion model with indexed motif residues, starting with randomly initialised coordinates.
• Generating one LigandMPNN (v\_32\_030\_25 weights, sampling temperature 0.1) sequence per design.
• Scoring by structure prediction and other metrics. We used AF3 as the structure prediction model for Kemp eliminase, carbene transfer, and nitrene transfer, and AF3-LA for serine esterase (both 4MU-Ac and DEHP).
• A single round of generation and filtering, without iterative partial diffusion or sequence design.

A note on naming of designs: the numbers used in our design names do not serve as a counter of our designs. Within facets used consecutive numbering, but when moving to a new facet or design round, we skipped to a larger round number.

#### 8.1 Kemp elimination

##### 8.1.1 Design rounds

We ran seven rounds of Kemp elimination design. The first two rounds were exploratory, testing different motifs and metrics. Rounds 3 and 4 evaluated these settings under various conditions, and round 5 demonstrated the added benefit of geometric metrics and our multiple-resequencing filtering strategy. We tested a total of 1,718 designs, with number of designs per round shown in Table S6.

**Table S6.** Number of designs in each round for Kemp elimination.

| Round | 1 | 2 | 3 | 4 | 5 | 6 | 7 | Total |
| --- | --- | --- | --- | --- | --- | --- | --- | --- |
| Number of designs | 268 | 188 | 180 | 182 | 364 | 173 | 363 | 1,718 |

##### 8.1.2 Input motifs

We used the theozyme geometries presented in [44], which are 2–4 residue motifs with the following roles:

• **Catalytic base:** Asp, Glu, His, His/Asp dyad, His/Glu dyad
• **Oxyanion hole:** Ser, Thr, Lys, Tyr, His, Arg, Asn, Gln
• **Aromatic interaction:** Phe, Tyr, Trp

Each input motif contains a catalytic base and an oxyanion hole, while the dyad and aromatic interaction are present in only a subset of motifs. While keeping the catalytic tip atoms fixed, we generated sidechain inverse rotamers for these motifs. We designed either against the substrate (5-nitrobenzisoxazole, PDB: H5J), or a transition state analog (6-nitrobenzotriazole, PDB: 6NT). During generation, we either fixed the full catalytic motif residues (full motif conditioning), or only the catalytic tip atoms (tip atom conditioning). Design rounds 1 and 2 were run on all motifs and included designs against the substrate and the transition state analog, while rounds 3–7 excluded the aromatic interaction residue, and were designed only against the transition state analog. For generation with an indexed model, catalytic residues were placed in a random order with intervening regions of randomly sampled length between 2–80 residues, and 10–100 residues at the N- and C-terminus.

##### 8.1.3 Kemp eliminase specific metric definitions

• `ligand_ca_neighbors_min/ligand_ca_neighbors_mean`: For each ligand atom, this metric counts how many protein Cα atoms are within each of three distance cutoffs (6, 8, 12 Å), then averages across cutoffs to get one “neighbor score” per ligand atom. `min` then indicates the minimum score among all ligand atoms, and `mean` the mean score. These metrics are intended to capture packing density. By requiring the mean to be above a certain threshold and the min to be below a certain threshold, we require a generally high packing density around the ligand, while retaining an opening.

• `cat_base_tip_ca_neighbors_mean`: This metric counts protein $C_{\alpha}$ neighbors within each of three distance cutoffs (6, 8, 12 Å) from the catalytic base tip atom, then averages across cutoffs. It measures how densely packed the catalytic site is.
• `protein_ligand_ipde_mean`: Structure prediction confidence: mean over the (protein × ligand) sub-matrix of the chain pair PDE matrix for structure prediction.
• `acid_base/distance`: Distance between the catalytic base's proton-abstracting atom and the substrate's heavy atom bonded to the abstracted proton.
• `oxyanion_oah/distance`: Distance between the oxyanion hole tip atom and the ligand atom developing a negative charge.
• `base_dyad/distance`: In some of our histidine catalytic base motifs, the histidine gets polarised by an additional motif residue. This metric measures their distance.
• `acid_base/dihedral2`: Out-of-plane angle of the catalytic tip atom relative to the substrate ring plane.
• `oxyanion_oah/dihedral2`: Out-of-plane angle of the oxyanion hole tip atom relative to the substrate ring plane.
• `base_dyad/dihedral1`: Out-of-plane angle of the histidine's nitrogen atom relative to the carboxyl group plane.

##### 8.1.4 Filters

The metrics in Table S7 were employed to filter designs in rounds 3 and 4, which can be viewed as a baseline. Metric distributions are shown in Figure S20. Depending on the subset, these were applied to either a single AlphaFold prediction or an aggregation of seven differently seeded predictions. In round 5, we added geometric filters (Table S8).

**Table S7.** Base filter thresholds for Kemp eliminase design (rounds 3 and 4).

| Metric | Threshold |
| --- | --- |
| *Sample metrics* |  |
| `sampld/ligand_ca_neighbors_min` | $\leq 18$ |
| `sampld/ligand_ca_neighbors_mean` | $\geq 18$ |
| `sampld/cat_base_tip_ca_neighbors_mean` | $\geq 15$ |
| *Apo structure prediction* |  |
| `apo/tm_score` | $\geq 80$ |
| `apo/motif_bb_aligned_allatom_rmsd` | $\leq 1$ |
| *Complex structure prediction* |  |
| `complex/tm_score` | $\geq 80$ |
| `complex/mean_pocket_bb_aligned_ligand_rmsd` | $\leq 2$ |
| `complex/protein_ligand_ipde_mean` | $\leq 1.9$ |

For computational efficiency, filtering was performed first on a single AlphaFold prediction ("seed\_0"), and then by aggregate metrics over multiple seeds. We applied these same filters with our multiple-resequencing critic, requiring that at least 6 out of 7 LigandMPNN resequences pass this filter set. Subsequently, we compiled facets to test APN sequences directly. We required these backbones to have 7/7 sequences pass the sequence ensemble filters (MR and MPD). For the MPD facet, we generated the scoring sequences using partial diffusion with backtracking by 575 sampling steps (out of 1000). The facets in rounds 1-6 used fixed motif residue indices. In round 7, we additionally tested our model with unindexed motifs. An overview, of screening activity by facet can be found in Figure S7.

**Table S8.** Filter thresholds for Kemp eliminase design (rounds 5-7, including geometric metrics). Aggregation indicates how metrics are summarised across 7 AlphaFold seeds.

| Metric | Threshold | Aggregation |
| --- | --- | --- |
| *Sample metrics* |  |  |
| sampled/ligand_ca_neighbors_min | $\leq 18$ | – |
| sampled/ligand_ca_neighbors_mean | $\geq 18$ | – |
| sampled/cat_base_tip_ca_neighbors_mean | $\geq 15$ | – |
| *Apo structure prediction (on seed_0 and as median-of-7)* |  |  |
| apo/tm_score | $\geq 80$ | median |
| apo/motif_bb_aligned_allatom_rmsd | $\leq 1$ | median |
| *Complex structure prediction (on seed_0 and as median-of-7)* |  |  |
| complex/tm_score | $\geq 80$ | median |
| complex/mean_pocket_bb_aligned_ligand_rmsd | $\leq 2$ | median |
| complex/protein_ligand_ipde_mean | $\leq 1.9$ | median |
| *Geometric metrics (mean-of-7 complex structure prediction)* |  |  |
| complex/acid_base/distance | $\leq 3.7$ | mean |
| complex/oxyanion_oah/distance | $\leq 3.7$ | mean |
| complex/acid_base/dihedral2 | 150–210 | mean |
| complex/oxyanion_oah/dihedral2 | 150–210 | mean |
| complex/base_dyad/distance | $\leq 3.7$ | mean |
| complex/base_dyad/dihedral1 | 150–210 | mean |

#### 8.1.5 Effect of motif choice

Further splitting up the success rate of the multiple-resequencing facet from rounds 5 and 6 by the input motif revealed that acid-based proton abstraction with aspartate and glutamate shows a higher combined success rate of 37% (32/86) compared with 10% (9/87) for histidine-based proton abstraction (Table S2).

### 8.2 Serine esterase (4MU-Ac and DEHP)

#### 8.2.1 Design rounds

We designed serine esterases for 4MU-Ac over 8 rounds. Round 1 used an early version of the input motifs and filters and did not yield any hits. Round 2 introduced the motifs described below. Rounds 3–6 prospectively tested various hypotheses but did not lead to conclusive insights or major improvements. However, data from these intermediate rounds were used to identify predictive metrics in retrospective analyses, which led to the large improvements afterwards. Round 7 prospectively tested the multi-resequencing filters. Round 8 tested sequences from the diffusion model, unindexed motif residue conditioning, and variations of the input motif (see below).

For DEHP, we performed three rounds of design. Round 1 established novel-scaffold hits. Round 2 included recycled-scaffold (partial diffusion) designs and novel-scaffold designs filtered using an initial version of multi-resequence filters. Round 3 used an improved implementation of partial diffusion and finalised multi-resequence filters as described below.

#### 8.2.2 Input motifs

To prepare input motifs for diffusion generation, we extracted catalytic residue coordinates from hydrolase structures (Table S9) co-crystallised with a tetrahedral intermediate analog [45, 46]. We included the catalytic Ser, His, and Asp/Glu, as well as the $n+1$ oxyanion hole H-bond donor residue, when present. We also included a second oxyanion hole residue, which was either a backbone amide or sidechain depending on the source enzyme. In one case there was a third oxyanion hole H-bond donor, which we included in a subset of designs derived from this motif. Into these motifs we placed RDKit conformers of 4MU-Ac and DEHP in a TI1 state (sp3 electrophile carbon) by superimposing them onto the crystal structure tetrahedral analog in both possible stereochemistries. Finally, motifs were filtered for clashes between ligand conformers and the catalytic residues.

In most cases, we defined the input motif to include all atoms of catalytic residues (full motif conditioning), except for backbone oxyanion hole residues, where we omitted sidechains from the motif and allowed the amino acid type to be designable (i.e. it was a resequence residue using the terminology of §3.1). In some cases we also included the C-term carbon of the residue N-term of a backbone oxyanion hole with the aim of improving motif scaffolding accuracy; however, this has a negligible effect. Finally, in a small subset of designs we allowed the backbone positions of catalytic residues to vary freely during design, either by using “tip atom” conditioning (motif includes only the functional atom on a sidechain), “tip frame” conditioning (motif only includes last rigid group on a sidechain), or inverse rotamer enumeration prior to input to the diffusion model as alternative full motif inputs. We obtained active enzymes with tip frame conditioning, but full motif conditioning using coordinates from PDB crystal structures had the highest *in silico* pass rates while still leading to novel scaffolds, so we used this approach in most designs. All of our most active 4MU-Ac designs come from full motif conditioning with 1XZM as the template PDB, while our best DEHPases come from 1R64.

Rounds 1–7 used indexed motif conditioning: catalytic residues were placed in a random order with intervening regions of randomly sampled length between 2–70 residues, and 5–100 residues at N- and C-terminus. The total length was restricted to 100–250 residues. A subset of round 8 used unindexed conditioning, where the total length was sampled from 100–250 and catalytic residues were allowed to be put in any residue position by the model. For both indexed and unindexed conditioning, motif residues within two residues of each other in the source PDB were kept together in the same relative position during design (i.e. as a single “motif island” [29]).

**Table S9.** PDB ID, chain ID, and residue number of catalytic motif residues used as input for serine esterase design. Amino acid type of oxyanion hole 2 or 3 is noted in parentheses. “BB” indicates backbone amide.

| PDB ID | Serine | Histidine | Asp/Glu | OAH1 | OAH2 | OAH3 |
| --- | --- | --- | --- | --- | --- | --- |
| 1R64 | B385 | B213 | B175 | B385 | B314 (Asn) | — |
| 1XZM | A120 | A188 | A175 | A121 | A42 (Ser) | A84 (Asn) |
| 1R50 | B77 | B156 | B133 | B78 | B12 (BB) | — |
| 1QFS | A554 | A680 | A641 | A555 | A473 (Tyr) | — |
| 2H5D | A195 | A57 | A102 | A195 | A193 (BB) | — |
| 1EAS | A195 | A57 | A102 | A195 | A193 (BB) | — |
| 6CHA | C195 | B57 | B102 | C195 | C193 (BB) | — |

#### 8.2.3 Multi-state structure prediction

We made AF3-LA predictions of designs in five catalytic states:

• **Apo:** Protein-only, no ligand.
• **ES:** Enzyme–substrate (Michaelis) non-covalent complex. Electrophile carbon on substrate is $sp^2$-hybridised and part of ester.
• **TI1:** First tetrahedral intermediate (after nucleophilic attack), with covalent bond between serine $O_\gamma$ and electrophilic carbon (which is $sp^3$-hybridised).
• **AEI:** Acyl-enzyme intermediate, with covalent bond between serine $O_\gamma$ and electrophilic carbon (which is $sp^2$-hybridised).
• **TI2:** Second tetrahedral intermediate (hydrolysis), with covalent bond between serine $O_\gamma$ and electrophilic carbon (which is $sp^3$-hybridised). Ligand has OH group (from water) instead of 4MU or ethyl-hexyl leaving group.

#### 8.2.4 Catalytic geometry metrics

Recent work showed that a strong predictor of serine esterase design success is whether certain atoms (representing important biochemical interactions) in predicted structures fall within certain distance, angle, and dihedral ranges. We adapted this approach to AF3 by defining “hydrogen bond” filters requiring that at least 1 out of 10 randomly seeded AF3-LA predictions satisfy the following constraints (defining $a_1$ and $a_2$ to be query atoms for the interaction, $b_1$ and $b_2$ as angle reference atoms, and $c_1$ and $c_2$ as dihedral reference atoms, respectively):

• Distance $a_1$–$a_2$ between 2.0 Å and 3.6 Å.
• Angle $b_1$–$a_1$–$a_2$ and $b_2$–$a_2$–$a_1$ are between 80°–160° if query atom is $sp^2$ and 69.5°–149.5° if middle atom is $sp^3$.
• Dihedral $c_1$–$b_1$–$a_1$–$a_2$ and $c_2$–$b_2$–$a_2$–$a_1$ are between 130° and 230° (or also in some cases –50° and 50°) if query atom is $sp^2$, and no constraint otherwise.

We evaluated interactions between:

• The triad histidine and the nucleophilic serine (“his\_nuc”).
• The triad histidine and triad carboxylic acid (“his\_acid”).
• First oxyanion hole hydrogen bond donor and the oxyanion (TI1, TI2) or carbonyl oxygen (ES, AEI) (“oah1\_oa”).
• First oxyanion hole hydrogen bond donor and the ester oxygen (“oah1\_ester”).
• Second oxyanion hole hydrogen bond donor and the oxyanion (TI1, TI2) or carbonyl oxygen (ES, AEI) (“oah2\_oa”).
• Second oxyanion hole hydrogen bond donor and the ester oxygen (“oah2\_ester”).

We defined “forward” H-bond filters that only consider the angle and dihedral around the first query atom $a_1$ rather than both directions (“hbond\_fwd”). For example, in the list of filters below, {apo, es, til, aei, ti2}/hbond\_fwd\_his\_nuc/mean indicates the mean occurrence (i.e. frequency) over 10 predictions, in each of the indicated states, of satisfying the distance and histidine $N_{\epsilon 2}$ angle and dihedral constraints. Requiring this value to be greater than zero just means that at least one of the 10 predictions satisfied the forward H-bond.

#### 8.2.5 Base filters

These are base filters for serine esterase design (Table S10), used exactly in and after 4MU-Ac round 6 and DEHP round 3, and with minor variations in earlier rounds. Metric distributions are shown in Figure S21.

Note that the TI2 state is filtered for OAH–ester H-bonds rather than OAH–OA as was intended. Because this led to active designs in round 2, it was maintained thereafter. Most designs satisfy both OAH–ester and OAH–OA H-bond filters in the TI2 state, likely due to OAH–OA filtering in other states.

#### 8.2.6 ES–TI1 consistency filters

We defined the following metrics to capture the ligand pose consistency between ES and TI1 states:

• es\_til\_min\_cross\_state\_rmsd: Minimum pocket-aligned ligand RMSD between any of 10 ES predictions and any of 10 TI1 predictions.
• es\_til\_cross\_seed\_ligand\_coord\_std: Root mean variance of ligand coordinates after aligning 10 ES predictions and 10 TI1 predictions on the ligand pocket.

These were not applied to rounds 1–6. In round 7, we filtered by:

• es\_til\_min\_cross\_state\_rmsd $\leq 5$
• es\_til\_cross\_seed\_ligand\_coord\_std $\leq 4.0$

In round 8, we filtered by:

• es\_til\_min\_cross\_state\_rmsd $\leq 1.0$
• es\_til\_cross\_seed\_ligand\_coord\_std $\leq 4.0$

#### 8.2.7 Multi-resequence (MR) and multi-partial-diffusion (MPD) filters

Round 7 introduced multi-resequence filtering. This inputs the backbone of a design to LigandMPNN and generates 10 “resequences”, predicts one seed for each resequence, and counts how many resequences out of 10 pass the following filters:

• {apo, til}/hbond\_fwd\_his\_nuc $> 0$
• {apo, til}/hbond\_his\_acid $> 0$
• til/hbond\_fwd\_oah1\_oa $> 0$
• til/hbond\_fwd\_oah2\_oa $> 0$
• {til, aei, ti2}/ligand\_plddt\_mean $\geq 80$

**Table S10.** Base filter thresholds for serine esterase design.

| **Metric** | **Threshold** |
| --- | --- |
| *Apo structure prediction* |  |
| apo/rmsd | $\leq 1.5$ |
| apo/protein_plddt_mean | $\geq 75$ |
| apo/motif_allatom_rmsd | $\leq 1.5$ |
| apo/tm_score | $\geq 80.0$ |
| apo/predicted_distance_error | $\leq 2.0$ |
| *T11 complex structure prediction* |  |
| ti1/interface_lddt | $\geq 50.0$ |
| ti1/ligand_plddt_mean | $\geq 80.0$ |
| ti1/fixed_res_bb_aligned_allatom_sc_rmsd | $\leq 3.0$ |
| ti1/motif_allatom_rmsd | $\leq 1.5$ |
| ti1/chain_pair_pde_mean_sym | $\leq 2.0$ |
| ti1/mean_pocket_bb_aligned_ligand_rmsd | $\leq 5$ |
| ti1/mean_pocket_bb_rmsd | $\leq 2.0$ |
| ti1/num_missing_atoms | $= 0$ |
| *Structure topology* |  |
| sampld/topology_string_len | $\geq 3$ |
| *Sequence quality* |  |
| num_missing_motif_residues | $= 0$ |
| max_single_aatype_frac | $\leq 0.25$ |
| max_consecutive_identical_aatypes | $\leq 5$ |
| num_blocklisted_motifs | $= 0$ |
| *Hydrogen bond geometry (mean across seeds)* |  |
| {apo, es, ti1, aei, ti2}/hbond_fwd_his_nuc/mean | $> 0$ |
| {apo, es, ti1, aei, ti2}/hbond_his_acid/mean | $> 0$ |
| {ti1, aei}/hbond_fwd_oah1_oa/mean | $> 0$ |
| {ti1, aei}/hbond_fwd_oah2_oa/mean | $> 0$ |
| ti2/hbond_fwd_oah1_ester/mean | $> 0$ |
| ti2/hbond_fwd_oah2_ester/mean | $> 0$ |
| *Substrate-specific* |  |
| ti1/ligand_plddt_mean/max (4MU-Ac) | $\geq 90.4$ |
| ti1/ligand_plddt_mean/max (DEHP) | $\geq 85$ |
| {aei, ti2}/ligand_plddt_mean/max (DEHP) | $\geq 88$ |

In retrospective analyses we also performed multi-seed prediction for each resequence (for 10 resequences x 10 seeds = 100 predictions per backbone), but this approach was not used to filter designs for experimental testing.

A facet of round 8 included diffusion model sequences with MPD filters. We applied the same filters as above for MR, but generated the scoring sequences using partial diffusion with backtracking by 575 sampling steps (out of 1000), which leads to outputs with 90% sequence identity to the input design (and negligible backbone difference). This number of steps was chosen after a hyperparameter sweep showed it to be the best compromise between creating diversity and maintaining high enough *in silico* pass rates to still filter for 10/10 MPD passing. The implied noise levels are indicated in Figure S31 (left) and shown next to training noise levels (right).

### 8.3 Styrene cyclopropanase (carbene transferase)

#### 8.3.1 Design rounds

We designed a single round of enzymes to catalyse the carbene transfer from ethyl diazoacetate (EDA) to styrene, resulting in ethyl 2-phenylcyclopropanecarboxylate S15A. Designable segment lengths from 10 to 100 residues were randomly sampled for either side of the heme-ligating histidine.

#### 8.3.2 Input motifs

For input motifs, we started with the density functional theory (DFT) theozyme geometries published in Kalvet *et al* [1], which contain a full heme cofactor, a heme-ligating histidine side chain, and a small molecule transition state in five different rotations. We then iterated through heme-ligating histidines in the PDB to identify suitable backbone coordinates for the histidine, clustering down to 64 conformations. We also represented the non-heme small molecules with two different representations: as two separate substrate molecules (PDB codes SYN and EEE); and as the actual product with a SMILES string. This results in $5 \times 64 \times 2$ input motifs to sample from for each of the two target products. Examples for both target *trans* products can be found in S15B.

#### 8.3.3 Metric Definitions

The following metrics are defined specifically for carbene transfer.

• `alanine_ratio`: proportion of alanine residues in the sequence.
• `transition_state_rmsd`: self-consistency RMSD computed on just the transition state excluding the haem cofactor.
• `heme_rmsd`: self-consistency RMSD on just the haem cofactor.

#### 8.3.4 Filters

The base filters applied to styrene cyclopropanase design are enumerated in Table S11. AF3 was used for structure prediction.

### 8.4 N-heterocycle synthase (nitrene transferase)

#### 8.4.1 Design rounds

We designed novel scaffolds for N-heterocycles in two rounds. For this reaction mechanism, first the nitrene intermediate is formed with the iron heme, ejecting dinitrogen in the process. This step is thought to be the rate-determining step, but for free heme will lead to an undesired reduction product. The regioselectivity for N-heterocycle formation is subsequently set by a hydrogen atom transfer step, where the nitrene interacts with the hydrogen of either the fourth or fifth carbon of the alkyl chain, forming the pyrrolidine or piperidine product, respectively.

In the first round, we designed for nitrene, pyrrolidine, or piperidine formation with the input motifs described below. In the second, we focused on piperidine formation, applying a stereoselectivity filter developed from the first round and also testing a theozyme obtained with DFT. Designable segment lengths from 1 to 278 residues were randomly sampled for either side of the heme-ligating histidine, with total lengths restricted between 60 and 280 residues.

**Table S11.** Base filter thresholds for styrene cyclopropanase design.

| Metric | Threshold |
| --- | --- |
| *Sample metrics* |  |
| alanine_ratio | < 0.225 |
| *Apo structure prediction* |  |
| apo/tm_score | > 80 |
| apo/rmsd | < 3 |
| apo/lddt | > 80 |
| *Complex structure prediction* |  |
| complex/mean_pocket_bb_aligned_ligand_rmsd | ≤ 2.25 |
| complex/mean_pocket_bb_rmsd | ≤ 1.1 |
| complex/predicted_distance_error | ≤ 1.5 |
| complex/interface_lddt | > 70 |
| complex/gdt_ha | > 60 |
| transition_state_RMSD | ≤ 2.25 |
| heme_RMSD | ≤ 2.25 |

#### 8.4.2 Input motifs

For the first round motif, we obtained motifs through AF3 structure prediction using the *TamPgb* and *ApePgb* sequences with three different input molecules as shown in Figure S3C: the nitrene intermediate, the pyrrolidine product, and the piperidine product. We started with eight structure predictions for each condition made with full MSA information, and selected a representative structure with all-atom RMSD on the motif atoms.

In the second round of design, we focused designs on motifs we obtained through DFT with the iron porphyrin, ligating histidine, and substrate in the regioselectivity-setting transition state. To complete the heme's atomic positions, we aligned heme conformers from the *TamPgb* structure prediction above as well as the optimised geometry from the Chemical Components Dictionary [47] to the core porphyrin present in the DFT-predicted geometry. When aligning, the core porphyrin contains 8-fold symmetry. We considered all permutations of the heme that do not result in a steric clash with the ligand.

#### 8.4.3 DFT for second round motif

DFT calculations were performed using an iron porphyrin model [48] to elucidate the reaction mechanism. All DFT calculations were carried out using the Gaussian 16 program [49]. Geometries of intermediates and transition states were optimised in the gas phase using the dispersion-corrected [50] B3LYP-D3(BJ) functional [51, 52] with a mixed basis set of SDD [53] (i.e., the Stuttgart/Dresden effective core potentials and the associated valence electron basis set) for Fe, and 6-31G(d) for other atoms. Vibrational frequency calculations were performed to confirm whether each structure is a local minimum or a transition state. Wavefunction stability of all structures was confirmed using the “stable=opt” keyword. Closed-shell singlet, triplet, quintet, and septet spin states of regioselectivity-determining hydrogen atom transfer (HAT) step were computed. Triplet was found to be the most stable spin state. We therefore based the motif on the triplet state structure of the HAT step.

#### 8.4.4 Metric definitions

For nitrene transfer samples, a potential H-bond partner is defined as any side-chain nitrogen or oxygen atom from residues Arg, His, Lys, Ser, Thr, Asn, Gln, or Tyr within 3.6 Å of the reference atom.

• `propionate_hbond_num_atoms`: the number of potential H-bond partner atoms to any of the heme propionate's oxygens (O1A, O2A, O1D, O2D), based on the structure prediction of the protein in complex with the heme and substrate
• `propionate_hbond_num_res`: the number of unique residues corresponding to the above

We also included two metrics to characterise the environment around the non-reactive end of the substrate, specifically as a proxy to solvent accessibility:

• `pocket_C9_min_atom_dist`: the minimum distance between any atom of the designed protein to the tail carbon atom ('C9') of the substrate's phenyl group, based on structure prediction with the substrate

• `pocket_C9_allatom_neighbors_mean`: the number of design atoms within 4, 6, or 8 Å, averaged across the different distance thresholds, of the substrate's 'C9' atom, based on structure prediction with the substrate

We also included metrics that quantify the variance of two relevant ligand atoms across structure prediction with five different representations of the small molecules and states involved: heme and substrate, heme and piperidine, heme and pyrrolidine, and the heme covalently bound to the substrate with two different SMILES representations. The SMILES representations are CC1=C(CCC(O)=O)C2=NC1=Cc1c(C=C)c(C)c3\C=C4/N=C(/C=C5\N([Fe])(CCCCC6=CC=CC=C6)N=N)n13)\C(=C/2)C(CCC(O)=O)=C5C)C(C)=C4C=C and [Fe]N(CCCCCC1=CC=CC=C1)N=N.CC1=C(CCC([O-])=O)/C2=C/C3=N/C(=C\C4=C(C)C(C=C)=C([N-]4)/C=C4\N=C(\C=C\1/[N-]\2)C(C=C)=C4C)/C(C)=C3CCC([O-])=O.

• `fe_self_rmsd`: the RMSD of the iron atom in the porphyrin across the five different structure predictions, after global alignment to the sampled structure
• `n_self_rmsd`: same as above, but for the ligating nitrogen of the substrate

Additionally, when sampling with tip atoms in coordinate replacement mode (APN-I), we noticed that some of the sampled histidine side-chains would not be ideally recapitulated. For this, we add the following metric to measure the deviation from the ideal geometry.

• `sample_his_rmsd`: the RMSD of the ligating His sidechain atoms from the ideal pentagonal plane

The above metrics were applied to both rounds of design. In the second round, we added an additional metric to quantify the pre-organisation of our designs in preference of pyrrolidine versus piperidine.

• `piperidine_propensity`: the difference in distance between nitrogen and the fifth carbon away of the substrate from structure prediction, and the distance between the nitrogen and the fourth carbon

##### 8.4.5 Filters

The metric thresholds in Table S12 were applied to inferences for nitrene transfer. Round 1 mostly used AF3 for structure prediction, while the second round used AF3-LA.

### 9 Experimental Methods

Unless noted otherwise, all chemicals, reagents, and consumables were obtained from commercial suppliers (Thermo Fisher, Sigma-Aldrich, or Merck) and used without further purification or treatment.

#### 9.1 Cloning and Transformation

DNA for clonal genes was purchased from Twist Bioscience (CA, USA). Genes encoding kemp eliminases were inserted into pET29b between the NdeI and XhoI cut site to contain a C-term 8×His tag. Genes encoding serine esterases, carbene transferases, nitrene transferases, and DEHPases were inserted into a modified pTriEx-4 vector to contain an N-term 8×His tag and HRV-3C cleavage site. For all genes, codons were optimised for *E. coli* using Twist Bioscience. For protein production, clonal genes from Twist Bioscience were directly transformed into chemically competent BL21(DE3) *E. coli* cells.

#### 9.2 Protein Production

##### 9.2.1 Protein expression and Ni-NTA-based purification

High-throughput protein production was performed at 37 °C, with shaking at 1,000 rpm (Infors shaking incubator with 3 mm throw) following inoculation of transformation mix into terrific broth (TB) with 0.4% (v/v) glycerol, autoinduction media (AIM; 0.05% glucose, 0.2% lactose), and the appropriate antibiotic, in sterile 2.2 mL deep-well plates.

Cultures were harvested by centrifugation at $2,568 \times g$ at 20 °C for 10 minutes using an Eppendorf 5810R centrifuge. Cell pellets were lysed with 40 µL Bugbuster (Merck) per well at 20 °C with shaking at 1,000 rpm. For kemp eliminases, Bugbuster was supplemented with cOmplete EDTA-free protease inhibitor (Roche). Following lysis, five replicate plates were pooled using an Integra ViaFlo and subsequently clarified by centrifugation at $2,568 \times g$ for 60 min at 20 °C. Clarified lysates were applied to pre-equilibrated 96-well HisPur™ Ni-NTA plates and eluted following the standard protocol in the relevant buffers, per enzyme type (Table S13).

**Table S12.** Nitrene transfer enzyme and specific filter thresholds.

| **Metric** | **Threshold** |
| --- | --- |
| *Sequence metrics* |  |
| seq_len | $\geq 90$ |
| max_consecutive_identical_aatypes | $\leq 5$ |
| max_single_aatype_frac | $\leq 0.25$ |
| num_blocklisted_motifs | $= 0$ |
| *Sample metrics* |  |
| sampld/percent_ligand_bb_clashes | $\leq 5.0$ |
| sampld_his_rmsd<sup>a</sup> | $\leq 0.04$ |
| *Nitrene-specific metrics* |  |
| n_self_rmsd | $\leq 0.8$ |
| pocket_C9_min_atom_dist | 2.8 – 3.8 |
| pocket_C9_allatom_neighbors_mean | 38 – 62 |
| propionate_hbond_num_atoms | 3 – 6 |
| propionate_hbond_num_res | 3 – 5 |
| his_heme_dist | $\leq 3.0$ |
| piperidine_propensity<sup>b</sup> | $\leq 0.35$ |
| *Apo structure prediction* |  |
| apo/tm_score | $\geq 80$ |
| apo/predicted_distance_error | $\leq 2.0$ |
| *Complex structure prediction* |  |
| complex/ligand_plddt_mean | $\geq 75$ |
| complex/interface_lddt | $\geq 30.0$ |
| complex/mean_pocket_bb_aligned_ligand_rmsd | $\leq 2.5$ |
| complex/mean_pocket_bb_rmsd | $\leq 2.0$ |
| complex/chain_pair_pde_mean_sym | $\leq 2.0$ |

<sup>a</sup> Relaxed to 0.08 for round 2

<sup>b</sup> Only applied to one facet of round 2.

**Table S13.** Buffers used in HTP protein purification.

| **Enzyme type** | **Ni-NTA equilibration buffer** | **Ni-NTA buffer** | **wash** | **Ni-NTA buffer** | **elution** | **SEC buffer** |
| --- | --- | --- | --- | --- | --- | --- |
| Kemp eliminases | 50 mM Tris pH 7.5, 500 mM NaCl, 0.005% Tween-20 | 50 mM Tris pH 7.5, 500 mM NaCl, 0.005% Tween-20, 20 mM imidazole |  | 50 mM Tris pH 7.5, 500 mM NaCl, 0.005% Tween-20, 250 mM imidazole |  | 50 mM Tris pH 7.5, 20 mM NaCl, 0.005% Tween-20 |
| Serine esterases | 50 mM HEPES pH 7.3, 500 mM NaCl | 50 mM HEPES pH 7.3, 500 mM NaCl, 20 mM imidazole |  | 50 mM HEPES pH 7.3, 500 mM NaCl, 250 mM imidazole |  | 20 mM HEPES pH 7.3, 50 mM NaCl |

### 9.2.2 Size exclusion chromatography (SEC)

SEC was performed in 96-well format using an ÄKTA Pure 25M system (Cytiva) with an ALIAS™ autosampler (Spark Holland). A Superdex 75 Increase 5/150 GL column (Cytiva) was connected directly to the injection module using 0.25 mm tubing, bypassing the column valve, as previously described [54]. 100 µL of Ni-NTA elution fractions were injected using the ALIAS' sample microlitre-pickup method in a 1 mL loop. SEC was performed at a flow rate of 0.5 mL/min, with 650 µL fractions collected into 2 mL deep well plates. To enable SEC of 192 samples overnight, injections were staggered while ensuring that no co-elution of salts and proteins occurred.

### 9.2.3 Protein concentration determination and normalisation

Protein concentrations were determined from SEC chromatograms using absorbance at 280 nm using custom Python code. For proteins with few or no aromatic residues and thus a negligible absorbance at 280 nm, concentrations were determined using absorbance at 205 nm with the $\epsilon_{205} = 31$ method on a NanoDrop One Spectrophotometer (Thermo Scientific).

Proteins were normalised to a standard concentration, typically 2 µM, using a Mantis liquid dispenser (Formulatrix). A continuous flow chip was used to dispense SEC buffer into 96-well deep well plates before purified proteins were added and mixed using a ViaFlo (Integra). Plates were briefly centrifuged at $500 \times g$ for 15 s at 20 °C.

### 9.3 Enzymatic activity screening assays (for $k_{obs}$)

#### 9.3.1 Kemp eliminase assay

Kemp eliminase activity was determined by monitoring the absorbance at 380 nm of 5-nitro-2-hydroxybenzonitrile product from 5-nitrobenzisoxazole (5-NBO, ChemScene) substrate. Assays were set up in 384-well clear-bottom absorbance plates (SpectraPlate-384 MB, Revvity) and monitored over time using a PheraStar FSX plate reader (BMG Labtech). Reactions were set up in 20 µL total volume composed of 10 µL of 2 µM enzyme solution in Kemp eliminase SEC buffer and 10 µL of 2 mM 5-NBO stock solution in Kemp eliminase SEC buffer supplemented with 10% DMSO. The 5-NBO stock solution was freshly prepared from powder in 100% DMSO to 0.5 M prior to diluting in the SEC buffer with 10% DMSO. Final assays thus contained Kemp eliminase SEC buffer, 1 µM of each designed protein, 1 mM 5-NBO, and 5% DMSO. Assays were initiated by the addition of enzyme solution to the reaction mixture using a 96-head Integra ViaFlo pipette (Z-height set to 33 mm to avoid scratching the bottom of the plate). Following a short centrifugation step ($\sim 15$ s at $500 \times g$), reaction plates were placed into the plate reader, and absorbance was monitored for 30 min at 25 °C. All reaction plates were set up with three in-plate controls such as buffer-only wells and positive control proteins to ensure that data remained consistent with previous screening rounds. Control proteins as well as the fastest enzymes were diluted to concentrations 1 µM or below to ensure linear reaction progress over the course of the measurement. Raw absorbance measurements were converted to product concentrations using the molar extinction coefficient of $15,800 \text{ M}^{-1} \text{ cm}^{-1}$ for product 5-nitro-2-hydroxybenzonitrile at 380 nm.

#### 9.3.2 Serine esterase assay

Serine esterase activity was determined by monitoring the fluorescence of 4-methylumbelliferone (4MU) product from non-fluorogenic 4MU acetate (4MU-Ac) substrate. Assays were set up in 384-well black shallow-well plates (Proxi-Plate, Revvity) and monitored using a PheraStar FSX plate reader equipped with a 350/450 optic module with a 3.5 OD attenuator, with gain set to 1535. Reactions were set up in 10 µL total volume composed of 5 µL of 2 µM enzyme solution in serine esterase SEC buffer and 5 µL of 0.2 mM 4MU-Ac stock solution, which was freshly prepared by diluting a 0.2 M stock solution in 100% DMSO into serine esterase SEC buffer containing 10% DMSO. Final assays thus contained 1 µM of each designed protein, 0.1 mM 4MU-Ac, and serine esterase buffer with 5% DMSO. The fastest enzymes were diluted to concentrations below 1 µM to ensure linear reaction progress over the course of the measurement. Assays were initiated by the addition of enzyme solution in duplicate to the reaction mixture using a 96-head Integra ViaFlo pipette. Following a short centrifugation step ($\sim 15$ s at $500 \times g$), the increase in fluorescence was monitored for 30 min at 30 °C. All reaction plates were set up with 4-6 in-plate controls such as empty wells, negative control proteins, positive control proteins, and best hits from previous design rounds, to ensure that data remained consistent with previous screening rounds. Wells containing only serine esterase SEC buffer and substrate were used to monitor the uncatalysed reaction. Raw fluorescence measurements were converted to product concentrations using a product standard curve.

#### 9.3.3 $k_{\text{obs}}$ fitting and hit calling

Custom Python code was used to determine initial rate constants $k_{\text{obs}}$ from progress curves of product concentration over time, after subtracting the uncatalyzed reaction progress from substrate-only wells (Figure S6). We found that accounting for the delay between assay initiation and the first measurement was important for reproducible results. We also computed a total turnover metric TTN@20min as the product concentration at 20 minutes normalised to the enzyme concentration. We defined a hit as any Kemp eliminase with a $k_{\text{obs}} > 0.038 \text{ s}^{-1}$ and TTN@20min $> 15$, or serine esterase with $k_{\text{obs}} > 0.003 \text{ s}^{-1}$ and TTN@20min $> 4$. We used the additional TTN@20min cutoff to ensure that hits can perform multiple turnovers. After automated hit-calling, we manually inspect progress curves to remove designs with poorly fitting data or bad replicates. Note that our $k_{\text{obs}}$ cutoffs for hit calling are much higher than the threshold of detection for non-zero activity, which is $k_{\text{obs}} = 0.0127 \text{ s}^{-1}$ for kemp eliminase (defined as 2-fold over buffer-only) and $k_{\text{obs}} = 0.0008 \text{ s}^{-1}$ for serine esterase (defined as the linear equivalent of TTN@20min = 1).

### 9.4 Michaelis–Menten kinetics

We quantified Michaelis–Menten parameters for a subset of designs against the substrates used for design. Assays were set up as described above but with an 8-fold serial dilution of substrate concentration. Serine esterases were thawed following storage at $-80^\circ\text{C}$, whereas Kemp eliminase proteins were assayed following storage at $4^\circ\text{C}$. Proteins were freshly diluted in their respective SEC buffers and dispensed into a 384-well assay plate. Assays were run in duplicate and always include a no protein control for background subtraction. For serine esterase, a freshly diluted product standard curve is run on the same plate as the samples.

For Kemp eliminases, substrate was added to the assay plate first, then protein was added to initiate the reaction. For serine esterases, protein-containing assay plates were briefly centrifuged for $\sim 15 \text{ s}$ at $500 \times g$ at $20^\circ\text{C}$ before the dilution series of substrate was transferred from a 384-well reservoir plate to the assay plate using an Integra ViaFlo (INTEGRA Biosciences, $384 \times 50 \mu\text{L}$ head). Plates were briefly centrifuged ($\sim 15 \text{ s}$ at $500 \times g$) then immediately transferred to a PheraStar FSX (BMG Labtech) plate reader to follow the reaction progress, as described in the rapid screening sections. To ensure low cycle times, only half of a 384-well plate was assayed at a time ($\sim 60 \text{ s}$ between datapoints) and the lowest substrate concentrations were measured first in each cycle.

Further kinetic characterisation at higher pH was done for Kemp eliminases to probe for an uplift in activity. The assay was done with the same buffer components except that Tris pH 7.5 was replaced with CHES pH 9.0 or CAPS pH 10.0 to maintain the corresponding pH. To account for the faster autoconversion of substrate to product at higher pHs, 5-NBO was prepared over eight dilution steps in 100% DMSO at 20X concentration before dispensing $1 \mu\text{L}$ to the assay plate and centrifuging briefly for $\sim 15 \text{ s}$ at $500 \times g$ at $20^\circ\text{C}$. Meanwhile, proteins were diluted to CHES or CAPS buffer at 1.05X concentration (buffers were also prepared at 1.05X concentration). The proteins used were purified using the standard protocol at pH 7.5 and diluted to the higher pH buffers with a dilution factor of at least 300X. To initiate the reaction, $19 \mu\text{L}$ of protein was added to $1 \mu\text{L}$ of substrate, briefly centrifuged ($\sim 15 \text{ s}$ at $500 \times g$) then immediately transferred to a PheraStar FSX plate reader to follow the reaction progress. Thus, the final reaction conditions contained, 50 mM CHES pH 9.0 or 50 mM CAPS pH 10.0, 20 mM NaCl, 0.005% Tween-20, 5% DMSO. Assays were set up in duplicate and run alongside a no protein control for background subtraction. To ensure low cycle times, only 1/8 of a 384-well plate was assayed simultaneously (30 s between datapoints) and the lowest substrate concentrations were measured first by the plate reader. The delay time between substrate addition and data acquisition was measured and used to correct initial rates. Analysis of the data was carried out as per the other kinetic measurements.

### 9.5 Thermal stability assays

The apparent thermal stability of designed enzymes, alongside natural protein human carboxylesterase 1 (CES1, BioTechne, 4920-CE-010), was determined using a thermal stability assay. Proteins were diluted in SEC buffer and $15\text{--}30 \mu\text{L}$ were added to a 384-well PCR plate (Biorad) in quadruplicate and sealed with a PCR microplate seal (Biorad). Samples were heated between $25\text{--}75^\circ\text{C}$ for Kemp eliminases and $30\text{--}80^\circ\text{C}$ for serine esterases in $10^\circ\text{C}$ increments for 10 minutes and cooled back to $20^\circ\text{C}$ using a CFX Opus 384 PCR machine (Biorad). The samples were then used to perform a rapid screening assay at $25^\circ\text{C}$ for Kemp eliminases or $30^\circ\text{C}$ for serine esterases. Technical repeats were pooled and $k_{\text{obs}}$ values calculated for each temperature and time point. Data were normalised to the mean $k_{\text{obs}}$ at $25^\circ\text{C}$ (Kemp eliminases) or $30^\circ\text{C}$ (serine esterases). The $T_{50}$ (the temperature at which 50% of the activity is lost) was determined by fitting normalised values to a shifted sigmoid function. Designs that did not exhibit loss of activity over the temperature gradient were reported as $T_{50} > 75^\circ\text{C}$ or $T_{50} > 80^\circ\text{C}$.

### 9.6 Nitrene transferase methods

#### 9.6.1 Protein expression

DNA clonal genes purchased from Twist Bioscience were directly transformed into chemically competent T7 express cells. Genes encoding nitrene transferases were inserted into a modified pTriEx-4 vector to contain an N-terminal 8×His tag and HRV-3C cleavage site.

**Protein expression in 96-well plates** To facilitate screening, 100 μL of T7 Express chemically competent cells were aliquoted into a PCR strip on ice. Using a multichannel pipette, 5 μL of cells and 1 μL of DNA construct were added to each well of a sterile, cone-shaped 96-well plate. Following a 30 minute incubation on ice, the plate was heat-shocked at 42 °C for 10 seconds and immediately returned to ice for 5 minutes. Cells were rescued with 100 μL of SOC medium and incubated at 37 °C for 30 minutes for outgrowth. Following outgrowth, 10 μL of each culture was inoculated into a 96-deep well plate preloaded with 400 μL/well of LB medium supplemented with ampicillin (100 μg/mL) (LB<sub>amp</sub>). These pre-cultures were incubated at 37 °C, 80% humidity, and 220 rpm shaking speed for 7–8 hours. Subsequently, 50 μL of each pre-culture was used to inoculate a secondary 96-deep well plate containing 950 μL/well of Terrific Broth supplemented with 100 μg/mL ampicillin (TB<sub>amp</sub>) per well. This expression culture was shaken at 37 °C and 220 rpm until an OD<sub>600</sub> of 0.8–1.0 was reached (typically 2–2.5 hours). The plates were then chilled on ice for 30–45 minutes and induced with 0.5 mM β-d-1-thiogalactopyranoside (IPTG) and 1.0 mM 5-aminolevulinic acid (ALA) (final concentrations). Protein expression was conducted at 22 °C and 220 rpm for 16–20 hours.

**Mid-scale protein expression** Single colonies from LB<sub>amp</sub> agar plates were picked using sterile toothpicks and cultured in a culture tube containing LB<sub>amp</sub> (5 mL/tube) at 37 °C with 220 rpm shaking speed for 7–8 hours. Subsequently, TB<sub>amp</sub> (50 mL) in a 125-mL Erlenmeyer flask was inoculated with an aliquot (1 mL) of these pre-cultures and allowed to shake for 2–2.5 hours at 37 °C, and 220 rpm until an optical density at 600 nm (OD<sub>600</sub>) of 0.8–1.0. The culture was then rested on ice for 30–45 min and induced with 0.5 mM IPTG and 1.0 mM ALA (final concentrations). Protein expression was conducted at 22 °C and 180 rpm for 16–20 hours.

#### 9.6.2 Analytical reaction setup and product quantification

Analytical reactions were performed in two formats: screening was conducted using whole-cell *E. coli* in 96-well plates, while quantitative validation was performed in individual vials using clarified lysates derived from protein-expressing *E. coli*.

**Reaction screening in 96-well plates** *E. coli* cells after expression in 96-well plates were pelleted by centrifugation (4,000 × g, 8 min, and 4 °C) and TB<sub>amp</sub> medium was removed. Harvested cell pellets were resuspended in 380 μL nitrogen-free M9 minimal medium (M9-N, pH 7.2). Resuspended cells in 96-well plates were introduced to an anaerobic chamber (coy chamber, oxygen level: <60 ppm). Substrate (5-azidopentyl)benzene was added using a multi-channel pipette (20 μL, 0.1 M stock in ethanol). The reactions were shaken at room temperature overnight at 700 rpm.

Following the overnight reaction, the 96-well plates were removed from the anaerobic chamber and the reactions were quenched with 800 μL acetonitrile. The resulting suspensions were mixed, then centrifuged (14,000 × g, 10 min, 4 °C). An aliquot of the supernatant (0.2 mL) was transferred to a 96-well high-throughput analytical plate and analysed by reverse-phase HPLC-MS. Products were identified by comparison with commercially available piperidine and pyrrolidine standards. Product quantification was determined using the selected ion monitoring (SIM) peak area ($m/z$ 162) and compared only within a single instrument run and experiment.

**Reaction validation in vials** *E. coli* cells after expression in 125-mL Erlenmeyer flasks were pelleted by centrifugation (4,000 × g, 8 min, and 4 °C) and TB<sub>amp</sub> medium was removed. Harvested cell pellets were resuspended in M9-N medium (pH 7.2) to OD<sub>600</sub> = 40. Lysate was obtained by sonication using a Qsonica Q500 sonicator (6 minutes, 1 second on, 2 seconds off, 35% amplitude, on wet ice). The cell debris was removed by centrifugation (14,000 × g, 10 minutes, 4 °C). Aliquots of the resulting clarified lysate (380 μL) were transferred to 2-mL screw cap vials and introduced to an anaerobic chamber (coy chamber, oxygen level: <60 ppm). Substrate (5-azidopentyl)benzene was added using a multi-channel pipette (20 μL, 0.1 M stock in ethanol). The reactions were shaken at room temperature overnight at 700 rpm.

Following overnight incubation, the reaction vials were removed from the anaerobic chamber and quenched with 800 μL of acetonitrile. The resulting mixtures were vortexed and centrifuged (14,000 × g, 10 min, 4 °C). An aliquot of the supernatant (0.2 mL) was transferred to a vial equipped with a glass insert for reverse-phase HPLC-MS analysis. Products were quantified using an ion-count calibration curve generated from a commercially available piperidine standard (selected ion monitoring, SIM $m/z$ 162); the curve was acquired on the same instrument and in the same sequence as the experimental samples.

**Analytical methods** Reverse-phase HPLC-MS was conducted using an Agilent Poroshell 120, EC-C18, 50 mm column, with acetonitrile (0.1 v% AcOH additive) and water (0.1 v% AcOH additive) as eluent. The enantiomeric excess (ee) was determined using polar ionic HPLC-MS method: single Agilent InfinityLab Poroshell 120 Chiral-V, 2.7 $\mu\text{m}$, 2.1 × 150 mm; eluent: 40% MeOH (0.2 w% $\text{NH}_4\text{CO}_2\text{H}$), 60% $\text{H}_2\text{O}$ (20 mM $\text{NH}_4\text{CO}_2\text{H}$, pH 4.0 adjusted with $\text{HCO}_2\text{H}$); flow rate: 0.2 mL/min.

#### 9.6.3 Hemochrome assay for the determination of hemoprotein concentration

*E. coli* cells after expression were pelleted by centrifugation ($4,000 \times g$, 8 min, and 4 °C) and $\text{TB}_{\text{amp}}$ medium was removed. Harvested cell pellets were resuspended in M9-N medium (pH 8.0) supplemented with D-glucose (25 mM final concentration) to $\text{OD}_{600} = 50$. The same cell suspension was used for the reaction set-up and determination of protein concentration. Protein concentration in the cell was determined using the hemochrome assay on cell lysate [55]. Lysate was obtained by sonication using a Qsonica Q500 sonicator (6 minutes, 1 second on, 2 seconds off, 35% amplitude, on wet ice). The cell debris was removed by centrifugation ($14,000 \times g$, 10 minutes, 4 °C). To a cuvette, 500 $\mu\text{L}$ of the lysate and 500 $\mu\text{L}$ of solution I [0.2 M NaOH, 40% (v/v) pyridine, 0.5 mM $\text{K}_3\text{Fe}(\text{CN})_6$] were added. The UV-Vis spectrum (380–620 nm) of the oxidised state Fe(III) was recorded immediately. Sodium dithionite (10 $\mu\text{L}$ of 0.5 M solution in water) was added, and the UV-Vis spectrum of the reduced state Fe(II) was recorded immediately. The protein concentration was calculated using the extinction coefficient, and accounting for the 2-fold dilution: $\varepsilon_{[557 \text{ reduced} - 540 \text{ oxidised}]} = 23.98 \text{ mM}^{-1}\text{cm}^{-1}$. The hemochrome assay detects total heme level, which is a good approximation of over-expressed heme enzyme.

### 9.7 DEHPase methods

#### 9.7.1 Protein expression

DNA clonal genes purchased from Twist Bioscience were directly transformed into chemically competent BL21(DE3) *E. coli* cells. Genes encoding DEHP esterases were inserted into a modified pTriEx-4 vector to contain an N-terminal 8×His tag and HRV-3C cleavage site.

**Protein expression in 96-well plates** Cells were grown overnight at a 1-mL scale in 2-mL 96-deep well plates, containing 500 $\mu\text{L}$/well of LB media supplemented with ampicillin at 100 $\mu\text{g}/\text{mL}$ ($\text{LB}_{\text{amp}}$), on a Multitron Infors shaker at 220 rpm and 37 °C. For expression, 50 $\mu\text{L}$ of overnight cultures were used to inoculate fresh 96-deep well plates containing 900 $\mu\text{L}$/well of TB media supplemented with ampicillin at 100 $\mu\text{g}/\text{mL}$ ($\text{TB}_{\text{amp}}$), then allowed to shake for 2 hours at 37 °C, 220 rpm, and 80% relative humidity. The plates were cooled on ice for 30 minutes and the cultures were induced with 0.5 mM IPTG. Expression was conducted at 22 °C, 220 rpm for 16–20 hours. Cells were then harvested by centrifugation for 10 min at $5000 \times g$ and 4 °C, then the supernatant was decanted. The cell pellets were then frozen at -20 °C for a minimum of 12 hours before reaction set-up.

**Mid-scale protein expression** Single colonies from $\text{LB}_{\text{amp}}$ agar plates were picked using sterile toothpicks and cultured in a culture tube containing $\text{LB}_{\text{amp}}$ (6 mL/tube) at 37 °C with 220 rpm shaking speed, grown overnight. A 1 mL aliquot of the starter culture was used to inoculate 50 mL of $\text{TB}_{\text{amp}}$ in a 125-mL sterile Erlenmeyer flask. The $\text{TB}_{\text{amp}}$ cultures were incubated at 37 °C (220 rpm) until the $\text{OD}_{600}$ reached 0.8–1.0. After cooling on ice for 30 minutes, protein expression was induced with 0.5 mM IPTG. The induction proceeded for 16–20 hours at 22 °C (220 rpm). Finally, cells were harvested in 50 mL Falcon tubes via centrifugation at $5000 \times g$ for 10 minutes at 4 °C. The supernatant was decanted, and the pelleted cells were frozen at -20 °C for a minimum of 12 hours until reaction set-up was conducted.

#### 9.7.2 Analytical reaction setup

Analytical reactions were performed using clarified *E. coli* lysates in two formats: screening in 96-well plates and validation in individual vials.

**Reaction screening in 96-well plates** Frozen cell pellets were thawed at room temperature for 15 min prior to lysis. Each well was resuspended in 500 μL of lysis buffer (10 mM potassium phosphate, pH 7.4, supplemented with 1 mg/mL lysozyme, 2 mM $\text{MgCl}_2$, and 0.07 mg/mL DNase I). The plates were agitated on an orbital shaker (800 rpm) and vortexed until pellets were fully resuspended, followed by a 1 hour incubation at 37 °C with shaking at 220 rpm. To clarify the lysate, the expression plates were centrifuged at $5000 \times g$ and 4°C for 10 min.

For the analytical reactions, 300 μL of clarified lysate was transferred to 2-mL 96-deep well plates containing 690 μL of 10mM potassium phosphate buffer (10 mM, pH 7.4) and 10 μL of a 1 M di(2-ethylhexyl) phthalate (DEHP) stock solution in DMSO, resulting in a final substrate concentration of 10 mM. The plates were foil-sealed and agitated on an orbital shaker at room temperature (800 rpm). After 4 hours, the reactions were quenched by transferring a 400 μL aliquot from each well into 400 μL of acetonitrile (MeCN) in a new 96-deep well plate. To facilitate protein precipitation, the mixtures were cooled at -20 °C for 30 minutes, followed by centrifugation at $5000 \times g$ for 10 minutes at 20 °C. Finally, 200 μL of the supernatant was transferred to a polypropylene 96-well microtiter plate (Agilent), heat-sealed with pierceable film (USA Scientific), and subjected to HPLC-MS analysis. The system was configured for selected ion monitoring (SIM) of mono-(2-ethylhexyl) phthalate (MEHP) at $m/z$ 277.0. Relative quantification was determined from the SIM peak area and compared only within a single instrument run and experiment.

**Reaction validation in vials** Frozen cell pellets were thawed at room temperature for 15 min, then resuspended in three volumes (w/v) of lysis buffer (10 mM potassium phosphate, pH 7.4, supplemented with 1 mg/mL lysozyme, 2 mM $\text{MgCl}_2$, and 0.07 mg/mL DNase I) relative to the pellet wet weight. The suspension was vortexed and incubated for 1 h at 37°C with shaking at 220 rpm. Following incubation, the mixture was transferred to microcentrifuge tubes and clarified via centrifugation at $14,000 \times g$ for 10 min at 4 °C. The resulting clarified lysate was collected into 15-mL conical tubes.

For the analytical validation, 300 μL of clarified lysate was added to 2-mL screw-cap glass vials (Agilent) containing 690 μL of potassium phosphate buffer (10 mM, pH 7.4) and 10 μL of a 1 M DEHP stock solution in DMSO, yielding a final substrate concentration of 10 mM. All reactions were performed in technical triplicate. The vials were capped and incubated on an orbital shaker (800 rpm) at room temperature for 4 hours. Reactions were then quenched by transferring a 400 μL aliquot into 300 μL of HPLC-grade MeCN and 100 μL of an internal standard solution (100 μM 4-butylbenzoic acid in 95:5 v/v MeCN:ddH<sub>2</sub>O). After mixing, the mixtures were cooled at -20 °C for 20 min to facilitate protein precipitation and subsequently centrifuged at $14,000 \times g$ for 15 min at 4 °C. Finally, 200 μL of the supernatant was transferred into glass inserts within 2-mL screw-cap vials for HPLC-MS analysis. Target compounds were quantified using the SIM peak area ratio of the analyte to the internal standard ($m/z$ 177.1).

**Analytical methods** Reactions were analysed by LC-MS using an Agilent 1260 Infinity II HPLC system. Chromatographic separation was achieved on an Agilent Poroshell 120 EC-C8 column (4.6 × 50 mm, 2.7 μm) maintained at 50 °C. The mobile phase consisted of ddH<sub>2</sub>O with 0.1% (v/v) acetic acid (Solvent A) and acetonitrile (MeCN) with 0.1% (v/v) acetic acid (Solvent B). A flow rate of 1.5 mL/min was used with the following step gradient: 5% B (0.0–0.5 min), 65% B (0.51–2.5 min), 95% B (2.51–3.0 min), and re-equilibration at 5% B (3.01–3.5 min). The mass spectrometer was operated in negative electrospray ionisation (ESI-) mode. For plate screening, the system was configured for selected ion monitoring (SIM) of mono-(2-ethylhexyl) phthalate (MEHP) at $m/z$ 277.0, with quantification estimated from the SIM peak area. For validation analysis, 4-butylbenzoic acid ($m/z$ 177.1) was employed as an internal standard; target compounds were quantified using the SIM peak area ratio of the analyte to the internal standard. Products were identified by comparing retention times and mass spectra to commercial standards. Data were processed using Agilent OpenLab CDS software.

### 9.8 Cyclopropanation methods

#### 9.8.1 Protein expression and purification protocol

Genes cloned in pET-29b were ordered from Twist Biosciences. Each construct included a C-terminal HRV-3C cleavage site followed by a Twin-Strep and 8×His affinity tags. For each construct, transformed BL21(DE3) *E. coli* cells were streaked onto LB agar wells from a 24-well plate supplemented with 50 μg/mL kanamycin, and incubated overnight at 37°C. Colonies were inoculated into 10 mL LB medium (Teknova) containing 0.5% glucose and 50 μg/mL kanamycin and grown overnight at 37°C with shaking at 200 rpm.

For each recombinant protein expression, 3 mL of preculture were diluted into 100 mL ZYM-5052 autoinduction medium (Teknova) containing 50 μg/mL kanamycin and 1 mM 5-aminolevulinic acid hydrochloride (Sigma-Aldrich; from a fresh 100 mM filtered stock solution) in 250 mL baffled flasks. Cultures were grown at 37°C with agitation at 200 rpm until an OD<sub>600</sub> of approximately 0.5 was reached. At this point, cultures were shifted to 18 °C for overnight expression. Cultures were split into two aliquots of 50 mL each and harvested by centrifugation ($4,000 \times g$, 20 min, 4°C). Pellets were stored at $-80^{\circ}\text{C}$ until purification.

#### 9.8.2 Small-scale protein purification

Purification was performed by batches of up to 48 constructs. 50 mL cell pellets were resuspended in 24-well plates with 5 mL lysis buffer (50 mM potassium phosphate pH 7.2, 200 mM NaCl, 0.5 mM TCEP) supplemented with benzonase (Millipore; 25 U/mL final) and EDTA-free protease inhibitor tablets (Roche). Samples were incubated at 300 rpm for 45 min before sonication (1.5 min total, 5 sec on/5 sec off, 40% amplitude, 24-tip probe). After lysis, hemin was added to a final concentration of 50 $\mu\text{M}$ and incubated for 30 min prior to centrifugation ($4,000 \times g$, 30 min, 4°C).

Clarified lysates were processed on a KingFisher™ Apex System using 50 $\mu\text{L}$ Pierce™ Ni-NTA magnetic agarose beads per construct. Following one hour incubation with lysates, the beads were washed three times with 1 mL Buffer A (50 mM potassium phosphate pH 7.2, 200 mM NaCl, 10 mM imidazole, 0.5 mM TCEP). Target proteins were eluted with Buffer B (50 mM potassium phosphate pH 7.2, 200 mM NaCl, 0.5 mM TCEP, 500 mM imidazole) in $\sim 90 \mu\text{L}$.

A second purification step was performed by SEC on a Superdex 200 Increase 5/150 column using an ÄKTA-Go system equipped with an ALIAS™ autosampler. For each construct (12 per session), 60 $\mu\text{L}$ of Ni-NTA eluate was loaded into a 250 $\mu\text{L}$ loop, 50 $\mu\text{L}$ of which were injected on the column. Elution was performed with 50 mM potassium phosphate pH 7.2 and 200 mM NaCl and 100 $\mu\text{L}$ fractions were collected. The main peak fractions (300 $\mu\text{L}$ per construct) were pooled, aliquoted in plates and/or vials, flash-frozen in liquid nitrogen and stored at $-80^{\circ}\text{C}$.

All purification buffers were degassed and purged with nitrogen prior to use. Protein concentrations were determined using Pierce™ 660nm Protein Assay. Purity and identity of purified targets were confirmed by SDS-PAGE, Western Blot and intact-mass spectrometry.

#### 9.8.3 Mid-scale protein expression and purification

Mid-scale expression and purification of selected constructs were performed as described for small-scale production, with the following modifications.

Culture volumes were increased to 2 L per construct and inoculated with 30 mL of preculture. Cell pellets were resuspended in lysis buffer, sonicated for 3 minutes (5 sec on/5 sec off, 40% amplitude), supplemented with 100 $\mu\text{M}$ hemin and clarified by centrifugation ($25,000 \times g$, 30 min, 4°C). Targets were purified using an automated Affinity-SEC workflow combining a 1 mL HisTrap™ FF Crude column with a Superdex™ 75 16/600 column operated in tandem. The affinity step included a 20 CV wash with Buffer A followed by step elution with 5.5 CV of Buffer B. SEC fractions corresponding to monomeric protein were pooled.

#### 9.8.4 Heme protein concentration determination

The concentration of the heme bound protein for the different constructs were determined using the Pyridine hemochromagen assay method [55]. The extinction coefficient for the Soret Band maxima ($157,000 \text{ M}^{-1} \text{ cm}^{-1}$) was used to calculate the protein concentrations [56].

#### 9.8.5 Carbene transferase assay protocol

Analytical scale biotransformations were performed in glass vials under anaerobic conditions ($\text{N}_2$ glove box). The total volume of the reaction was 348 $\mu\text{L}$. To 284 $\mu\text{L}$ of assay buffer (50 mM KPi, 200 mM NaCl, pH 7.2, $\text{N}_2$ degassed overnight), 40 $\mu\text{L}$ of the test protein was added, followed by the addition of styrene (10 $\mu\text{L}$, 40 mM stock in MeCN) and ethyl diazoacetate (EDA) (10 $\mu\text{L}$, 400 mM stock in MeCN). This was followed by addition of sodium dithionite (4 $\mu\text{L}$, 10 mM stock in assay buffer). The vials were sealed and incubated outside the glove box ($25^{\circ}\text{C}$, 200 r.p.m.) for 2h. Finally, the reactions were quenched with addition of 30 $\mu\text{L}$ HCl (3 M) and agitated on a shaker for 10 minutes. Subsequently, 500 $\mu\text{L}$ of *n*-butanol containing 0.1 mmol TMB (trimethoxybenzene) was added, and the plate was shaken for an additional 10 minutes. The mixture was then allowed to stand for 10 minutes to facilitate phase separation. The organic layer was collected, dried over anhydrous magnesium sulfate and transferred to a 96-well analytical plate. Samples were submitted for chiral liquid chromatography (LC) analysis.

#### 9.8.6 LC Method

Chromatographic analysis of the enzyme assay products was performed using a Waters ARC HPLC-MS system. The instrument configuration included an Acquity™ Arc Quaternary Solvent Manager-R pump and an Acquity Arc Sample Manager FTN-R injector. Detection was carried out using a Waters 2998 Photodiode Array (PDA) detector monitoring

UV absorbance at 215 nm, coupled in-line with a Waters QDa mass detector. Chromatographic separations were achieved on a Chiralcel OJ-H chiral column (4.6 mm × 250 mm, 5 µm particle size). The method utilised an isocratic mobile phase consisting of heptane and isopropanol (70:30, v/v) delivered at a constant flow rate of 1.0 mL/min. The total runtime for each injection was 12 minutes.

#### 9.8.7 Scale-up procedure

Under anaerobic conditions (N<sub>2</sub> glove box) 25.6 mL of assay buffer (50 mM KPi, 200 mM NaCl, pH 7.2, N<sub>2</sub> degassed) was added to a 250 mL bottle,. This was followed by the addition of styrene (2 mL, 200 mM stock in MeCN), followed by 48 mL of the enzyme (41.55 µM stock of heme-loaded enzyme). Afterward, sodium dithionite (2 mL, 100 mM stock in assay buffer) was added, followed by slow addition of ethyl diazoacetate (EDA) (2.4 mL, 500 mM stock in MeCN) over 30 minutes (by a pipette drop by drop over 1 minute, every 5 minutes) while shaking. The reaction bottle was incubated at 25°C, on a shaker at 200 rpm for 2 hours. Finally, the reactions were quenched with addition of HCl.

#### 9.8.8 Work up

The reaction mixture was quenched with 3 M HCl (50 mL) and agitated for 10 minutes followed by extraction with ethyl acetate (2 × 100 mL). The organic phases were combined, dried over sodium sulphate, filtered and concentrated under vacuum. The crude residue was purified by flash column chromatography (Biotage Selekt, 5 g Sfar column) eluting with diethyl ether/cyclohexane (0–100%). The fractions of interest were pooled and evaporated to yield *trans*-ethyl (1*S*,2*S*)-2-phenylcyclopropanecarboxylate (13.4 mg, 96% purity) as a colourless oil.

#### 9.8.9 Product Validation

To verify the identities of the two *trans* enantiomers in the reaction mixtures from GDM\_CT\_0103, standard co-injection experiments were performed using authenticated reference standards (Figure S15C). Below, DP1 refers to the *RR* product and DP2 refers to the *SS* product, which GDM\_CT\_103 was designed for.

To validate DP1 in the reaction mixture, initial chromatographic analysis of the reaction mixture from GDM\_CT\_103 exhibited a peak for the desired DP2 at a retention time ($t_R$) of 5.41 min, with trace amount of DP1 present at the expected time. To validate the elution profile of DP1, the sample in well C3 was spiked with 1 µL of a DP1 standard stock solution prepared in *n*-butanol (*n*-BuOH). Subsequent analysis of the spiked mixture revealed the emergence of a new distinct peak at the expected $t_R$ of 4.65 min, cleanly resolving from DP2 and confirming the specific retention time of the undesired product.

For DP2, initial analysis of the reaction mixture from well C4 confirmed the presence of DP2, eluting at $t_R$ 5.41 min with a baseline peak area of 5204 (representing 42% relative area). To verify this peak assignment, the sample in well C4 was spiked with 1 µL of a DP2 standard stock solution in *n*-BuOH. Following co-injection, the chromatogram showed an exact co-elution, with the target peak at $t_R$ 5.41 min exhibiting a proportional increase in area to 13,657 (64% relative area). No other changes to the chromatographic profile were observed, positively confirming the identity of the desired product DP2.

The identity of the product from the scale-up of GDM\_CT\_103 was further validated with a proton NMR in Figure S15D.
